# Division-resolved inference of flow and trajectories in proliferating cell populations

**DOI:** 10.64898/2026.09.03.747958

**Authors:** Geunwoo Shin, Teemu P. Miettinen, Joon Ho Kang

## Abstract

Interpreting cellular responses requires understanding how cellular states change over time, yet many scalable assays provide only population snapshots. In proliferating populations, changes in population distributions reflect both state progression within cells and redistribution caused by division. We introduce division-resolved inference of flow and trajectories (DRIFT), a computational framework that infers the dynamics of a measured cellular state from population distributions collected over time, without synchronizing or tracking individual cells. DRIFT solves a population-balance equation to separate state progression from the redistribution caused by cell division. In simulations of growth and division perturbations, DRIFT recovered the ground-truth mean volume trajectories across simulated single-cell lineages. Applied to live L1210 leukemia cells, DRIFT inferred perturbation-specific volume trajectories consistent with longitudinal single-cell measurements. Beyond volume, the framework captured cell area dynamics in live HeLa cells, with validation by continuous imaging. Its application to an endpoint molecular assay further yielded DNA-content dynamics from fixed-cell flow cytometry in L1210 cells, consistent with complementary analyses of DNA synthesis. Overall, DRIFT converts population measurements into division-resolved cellular dynamics, extending the utility of routine population assays to kinetic studies of cellular progression when longitudinal tracking is impractical.

## Introduction

Cellular processes, including growth, cell-cycle progression, and division, involve coordinated molecular activities and changes in physical state^1,2^. Transcription, translation, metabolism, and cell-cycle regulation collectively shape measurable cellular states, including physical quantities such as cell size and mass, and molecular quantities such as DNA-content, transcript and protein abundances, and the activities of signaling molecules and cell-cycle regulators^3^. Perturbations can alter biosynthesis, cell-cycle progression, division control, or combinations of these processes, with distinct mechanistic consequences^4,5^. Translation inhibition, for example, reduces biomass accumulation^6^, whereas Aurora B inhibition allows continued growth while blocking cytokinesis and driving cells toward higher ploidy^7^. Determining how these measured cellular states change within individual cells over time has traditionally relied on longitudinal measurements. At the molecular level, such measurements have revealed time-dependent changes in gene expression, protein abundance, and signaling activity^8,9^. At the physical level, such measurements have revealed size-control strategies in bacteria^10^, budding yeast^11^, and mammalian cells^4,12^, as well as cell-cycle-dependent mass accumulation in mammalian cells^6,13–16^.

Longitudinal single-cell platforms, including suspended microchannel resonators^13,14^, microfluidic mother machines^17^, quantitative phase imaging^15^, and fluorescence-exclusion microscopy^4^, can measure growth with high precision through changes in mass, cell length, area, or volume. Fluorescent reporters and biosensors extend longitudinal tracking to selected molecular quantities and activities but require compatible reporters and repeated live-cell imaging^18,19^. However, even when technically feasible, longitudinal measurements remain difficult to scale across diverse perturbation conditions. Moreover, several molecular measurements are inherently destructive. Stoichiometric DNA staining, fixed-cell immunofluorescence, and single-molecule RNA hybridization are performed on fixed cells and therefore cannot be repeated in the same cell^20,21^. Live-cell DNA dyes provide an alternative but induce a DNA damage response and perturb cell-cycle progression during prolonged exposure^22^. Consequently, total DNA-content and the abundances of many RNAs and proteins remain difficult to follow quantitatively within individual cells throughout a perturbation.

High-throughput population measurements provide a complementary source of kinetic information. A Coulter counter measures volume distributions from thousands of cells within seconds^23^, while flow cytometry and automated microscopy provide distributions of molecular and morphological states. Several computational frameworks infer cellular dynamics from population distributions without tracking individual cells. RNA velocity and scVelo estimate the direction of progression from intracellular kinetic relationships^24,25^, whereas Waddington-OT and probability-flow inference reconstruct transport between measured distributions using optimal transport or stochastic-process models^26,27^. Methods such as DeepVelo learn continuous transcriptomic velocity fields and integrate them to predict cell-state trajectories^28^. Other approaches, including pseudodynamics, dynamic unbalanced transport, and one-shot spatial reconstruction, incorporate changes in cell number by inferring transport together with source, sink, or net-growth terms^29–31^. However, changes in cell number alone do not specify how division redistributes cells along the measured state coordinate.

In proliferating populations, extensive cellular quantities that define state coordinates, including cell volume, dry mass, DNA-content, and ribosome abundance, increase over at least part of the cell cycle and are partitioned between two daughter cells at cell division^3^. Therefore, a state-resolved population model should represent binary division as the removal of one mother cell from its pre-division state and the entry of two daughter cells into their corresponding post-division states. For cell volume, a shift in the population distribution toward larger values can reflect an increased volume-growth rate, delayed division, or both. Time-series population distributions alone therefore do not uniquely determine the contributions of state progression and division. When neither the velocity field nor the division-state density is independently specified, the two must be inferred jointly, subject to a biologically defined partition rule.

Ergodic rate analysis (ERA) infers state-dependent progression in balanced, asynchronously proliferating populations at steady state. It represents binary division through a renewal boundary that links a mother cell leaving a single terminal state to two daughter cells entering a single initial state^32^. This ergodic principle has since been applied to infer transcriptional activity, cell-mass variation, and cell-cycle-resolved phosphoprotein dynamics and feedback from fixed-cell measurements^33–36^. Its steady-state formulation, however, does not directly address drug-perturbed populations undergoing transient changes in growth or division dynamics. Population-balance equations express the same conservation principle in a more general form, relating changes in a measured distribution to state progression, mother-cell division, daughter-cell births, and loss^37–39^. The Collins–Richmond method provided an early steady-state inverse formulation of this balance for cell growth^40^. Because the measured distributions alone do not determine a unique solution, previous inverse approaches have closed the problem primarily in two ways: by assuming or supplying birth and division terms, often using separate measurements^39–42^, or by prescribing a single-cell growth profile and inferring only the division rate^43,44^. Neither form of information is generally available from standard population measurements collected over time after drug treatment. Moreover, prescribing a single-cell growth profile would presuppose the perturbation response under investigation, because the perturbation may alter state progression, division control, or both. Transient inference therefore requires a computational procedure for selecting stable, biologically plausible velocity fields and division-state densities among the multiple solutions consistent with the population balance and the measured data^45^.

Here, we introduce division-resolved inference of flow and trajectories (DRIFT), a population-balance method that infers the progression of measured cellular states from population distributions collected over time, without synchronizing or tracking individual cells. Within a transient population balance, DRIFT jointly estimates the velocity field and division-state density as functions of state and time and reconstructs characteristic trajectories from the inferred fields (Fig. 1). In simulations of growth and division perturbations, the DRIFT-inferred growth kinetics, division-state densities, and characteristic trajectories matched their ground-truth counterparts (Figs. 2 and 3). We then applied DRIFT to Coulter-counter volume measurements in drug-treated L1210 leukemia cells (Fig. 4) and extended the framework to imaging-based projected-area measurements in live HeLa cells and DNA-content measurements from fixed L1210 cells (Fig. 5). Together, these applications extend the use of routine assays from measuring population distributions to inferring cellular dynamics.

**Fig. 1.**
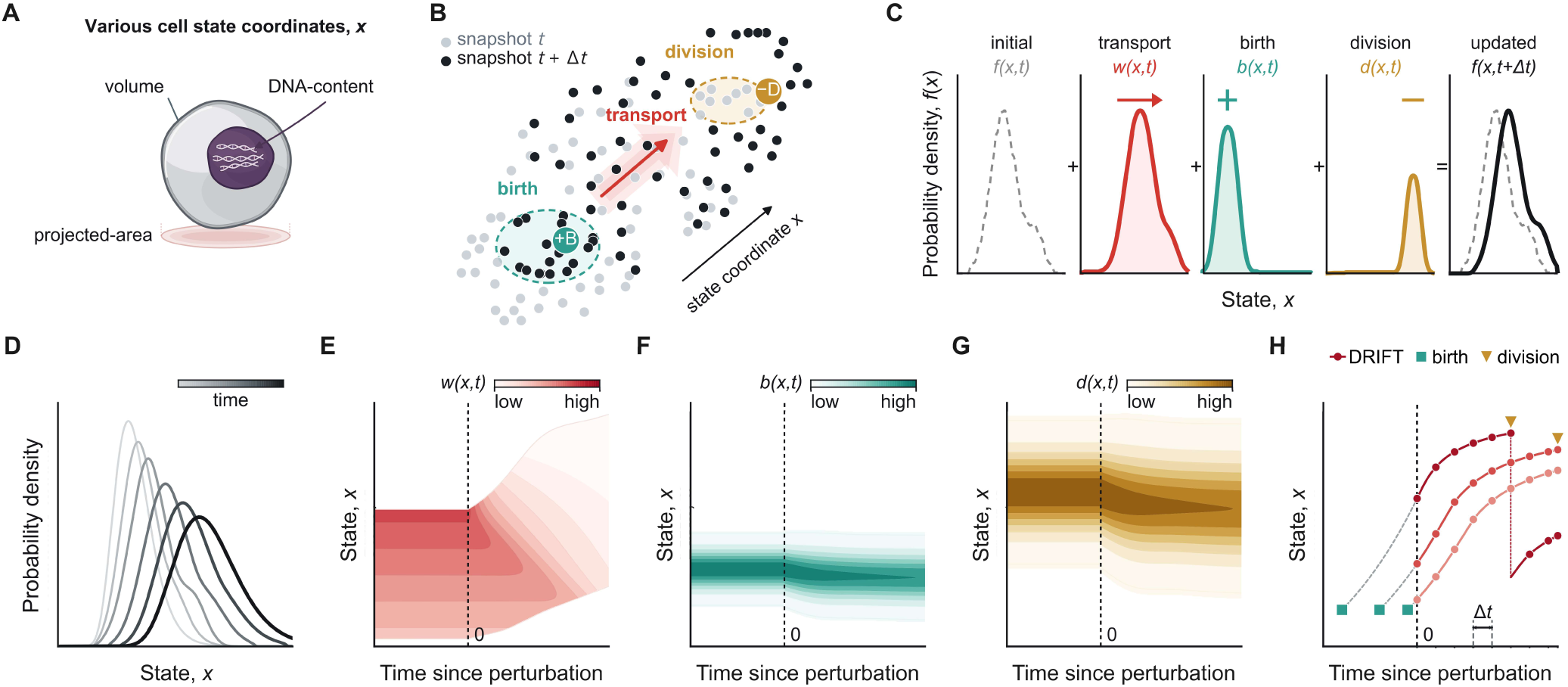
Overview of division-resolved inference of flow and trajectories (DRIFT). (**A**) Cellular state coordinates *x* considered in this study, including cell volume, projected-area, and DNA-content. (**B**) Population snapshots acquired at times *t* and *t + Δt*. State progression moves cells along the measured state coordinate, mother-cell division removes cells from their division states, and daughter-cell birth adds two cells at lower daughter states. (**C**) Schematic decomposition of the change in the normalized state probability density *f(x,t)* into state progression governed by the velocity field *w(x,t)*, birth-state density *b(x,t)*, and division-state density *d(x,t)*, producing the updated distribution at *t + Δt*. (**D**) Time-series normalized state probability densities *f(x,t)*; shading denotes successive observation times. (**E**–**G**) Illustrative state- and time-dependent fields for *w(x,t)*, *b(x,t)*, and *d(x,t)*, respectively. The vertical dashed line denotes perturbation at *t* = 0, and color intensity denotes field magnitude. (**H**) Illustrative DRIFT-inferred characteristic trajectories, which can be obtained by integrating the velocity field *w(x,t)*. Trajectories can be initialized from different states at the time of perturbation. Teal squares and ochre triangles denote birth and division events, respectively; gray dashed segments denote pre-perturbation progression, red segments denote post-perturbation progression, and *Δt* indicates the snapshot interval.

**Fig. 2.**
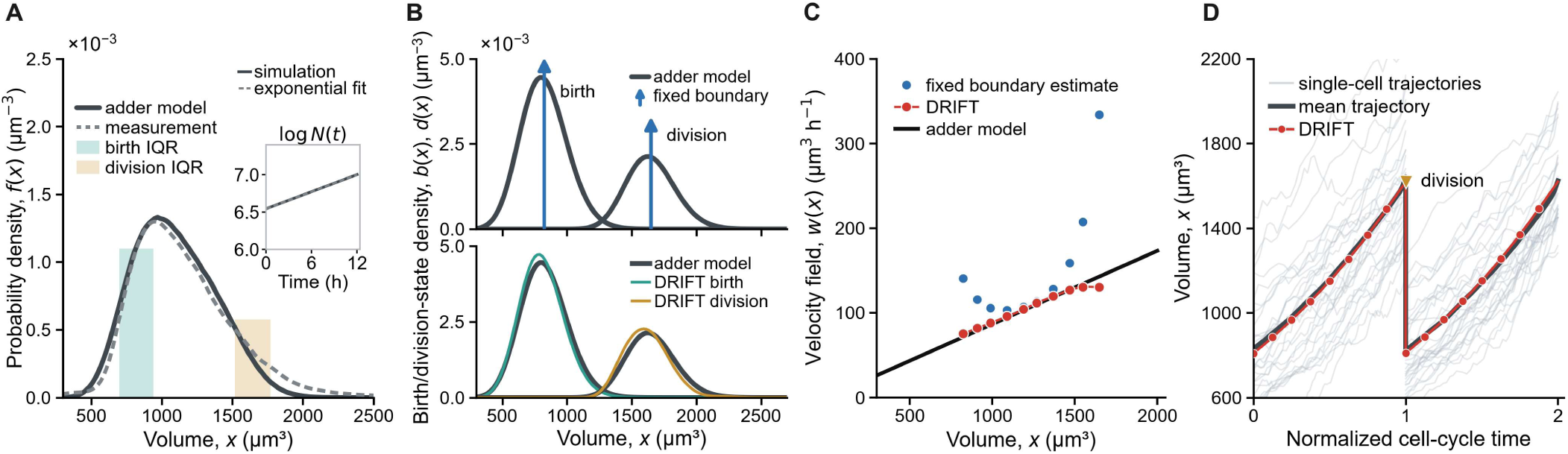
DRIFT infers distributed birth- and division-state densities and the volume-growth velocity field in a simulated adder population. **(A)** Stationary volume probability density *f(x)* from a simulation of adder-like size homeostasis with stochasticity (black solid line; Supplementary Note 3) and the Coulter counter measurement for comparison (dashed gray). Teal and ochre bands denote the interquartile ranges of the simulated birth- and division-state densities, respectively. The inset shows the total cell count *N(t)* on a logarithmic scale (solid) with an exponential fit (dashed). **(B)** Birth-state density *b(x)* and division-state density *d(x)* on a common volume coordinate. Dark curves in both panels are the adder-model densities. The upper panel shows the fixed-boundary formulation, which assigns daughter-cell birth and mother-cell division to single states (blue arrows), and the lower panel shows the corresponding distributed DRIFT estimates for birth (teal) and division (ochre). **(C)** Velocity field *w*(*x*) from the adder model (black solid), fixed-boundary estimate (blue), and DRIFT estimate (red). **(D)** Individual simulated cell-volume trajectories (light gray; 30 lineages shown), the ground-truth mean trajectory of the same 30 lineages (black), and the DRIFT-inferred characteristic trajectory (red), shown over two normalized cell cycles (Supplementary Table 3). The inverted triangle denotes the division event, and red circles mark positions at 1-h intervals. All panels show results from one representative simulation among *n* = 10 independent simulations.

**Fig. 3.**
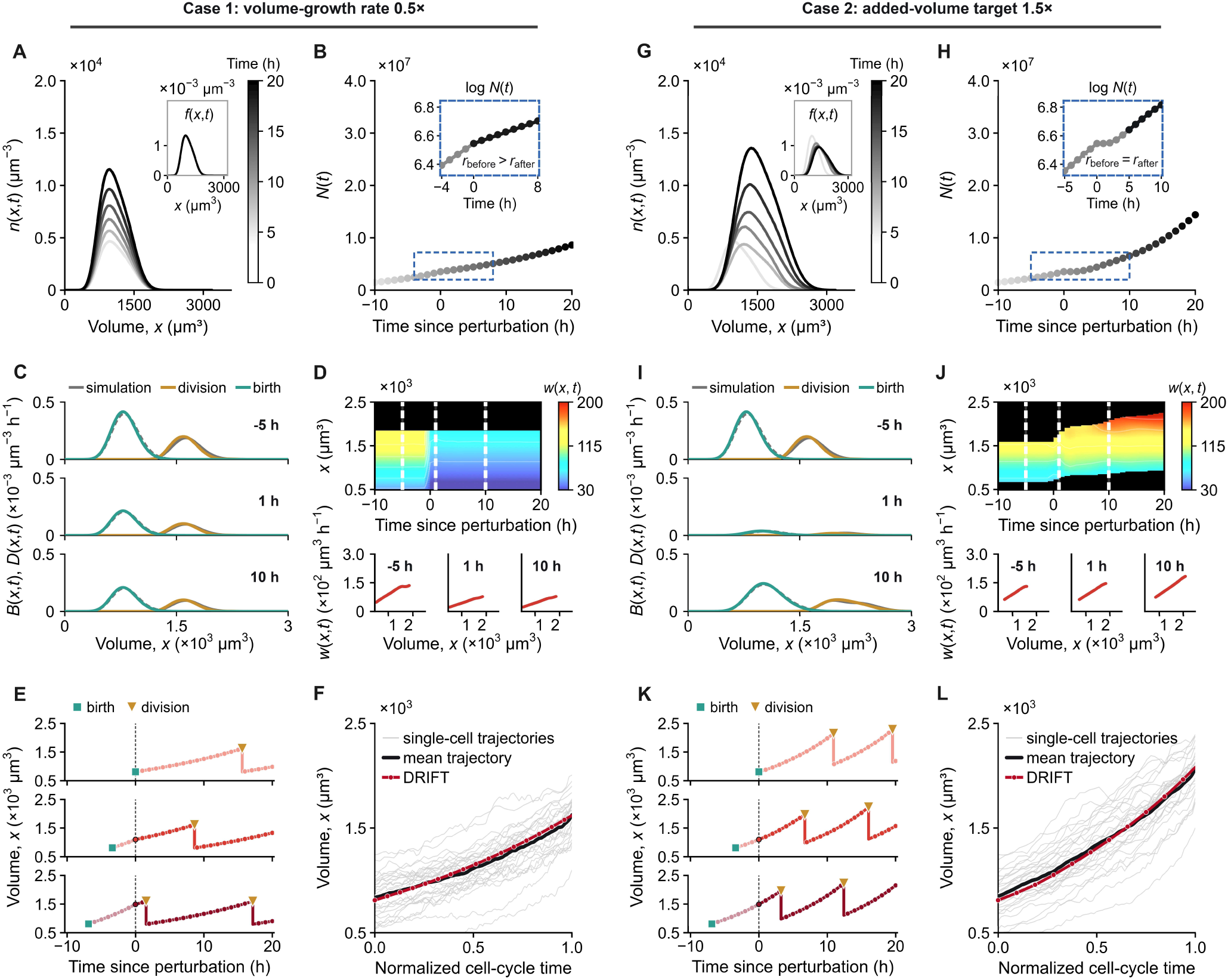
DRIFT recovers growth and division kinetics during perturbations to volume-growth rate and division-size control in simulated adder populations. **(A–F)** Case 1, in which the single-cell volume-growth rate in a stochastic-adder population was halved at *t* = 0 h while the division-size rule remained unchanged. **(G–L)** Case 2, in which the single-cell volume-growth rate remained unchanged and the added-volume target was increased to 1.5 times its pre-perturbation value at *t* = 0 h. (**A** and **G**) Time-series population distributions *n(x,t)*; grayscale denotes time, and the insets show the corresponding normalized state probability densities *f(x,t)*. (**B** and **H**) Total cell count *N(t)*. Dashed blue boxes mark the regions enlarged in the insets, which show log *N(t)* around the perturbation. The annotations compare the net population growth rate before and after the perturbation. (**C** and **I**) Daughter-birth and mother-division-loss densities *B(x,t)* and *D(x,t)* at −5, 1, and 10 h relative to the perturbation, shown for the simulation (gray) and DRIFT (teal for birth and ochre for division). (**D** and **J)** Inferred velocity fields *w(x,t)*. The lower panels show velocity profiles at −5, 1, and 10 h, corresponding to the dashed vertical lines in the heat maps; black regions denote state–time bins outside the displayed support. (**E** and **K)** Representative DRIFT-inferred characteristic trajectories initialized from three states (from top to bottom: newborn-like, mid-cycle, and near-division; Supplementary Note 2) at *t* = 0 h. Teal squares and ochre triangles denote birth and division events, respectively; pale red segments denote pre-perturbation progression, red segments denote post-perturbation progression, and the vertical dashed line denotes the perturbation at *t* = 0 h. (**F** and **L)** Simulated and DRIFT-inferred trajectories on a common normalized cell-cycle time since perturbation. Light gray, black, and red curves indicate 30 individual lineages, the mean trajectory of those 30 lineages, and the DRIFT-inferred characteristic trajectory, respectively, in each panel; red circles mark positions at 1-h intervals. All panels show results from one representative simulation among *n* = 10 independent simulations.

**Fig. 4.**
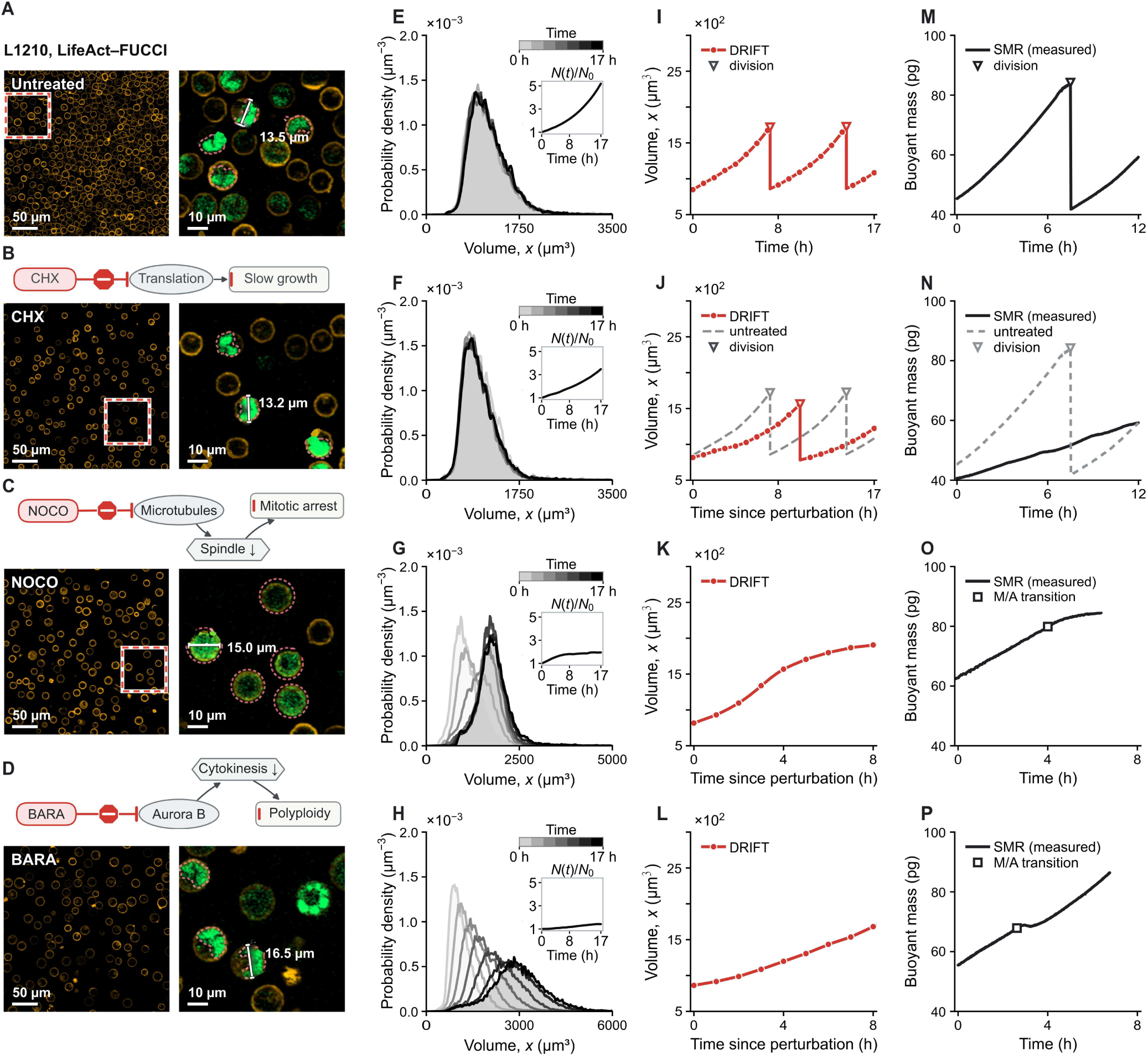
DRIFT reconstructs condition-specific volume dynamics consistent with longitudinal single-cell buoyant-mass traces in L1210 cells. Rows correspond to untreated (**A, E, I, and M**), cycloheximide-(CHX; B, F, J, and N), nocodazole-(NOCO; C, G, K, and O), and barasertib-(BARA; D, H, L, and P) treated L1210 leukemia cells. (**A–D**) Drug-target schematics and representative fluorescence micrographs of L1210 cells expressing LifeAct (red) and FUCCI (green) at *t* = 17 h since perturbation. Dashed boxes identify the enlarged regions shown at right, in which pink dashed outlines trace the FUCCI-positive region and white bars indicate representative cell diameters; scale bars, 50 µm and 10 µm for the field and enlarged images, respectively. (**E–H**) Time-series volume probability densities *f*(*x*,*t*) from Coulter counter measurements, averaged over the *n* = 1, 1, 2, and 3 independent experiments for untreated, CHX, NOCO, and BARA, respectively. Grayscale denotes treatment time, and the insets show the normalized cell count *N*/*N*_0_. (**I–L**) DRIFT-inferred characteristic trajectories initialized at the birth state at *t* = 0 h. Red circles mark positions at 1-h intervals. Inverted triangles denote division events; vertical segments denote daughter-state resets. The gray dashed curve in panel J denotes the untreated DRIFT reference from panel I. (**M–P**) Longitudinal single L1210 cell buoyant-mass traces measured using suspended microchannel resonators and reported previously^6,7^. Inverted triangles in panels M and N denote division. In panel N, the untreated trace is shown as a dashed reference. Each open square in panels O and P denotes the operationally assigned metaphase-to-anaphase (M/A) transition, following prior SMR studies of mitotic growth^6,7,16^.

**Fig. 5.**
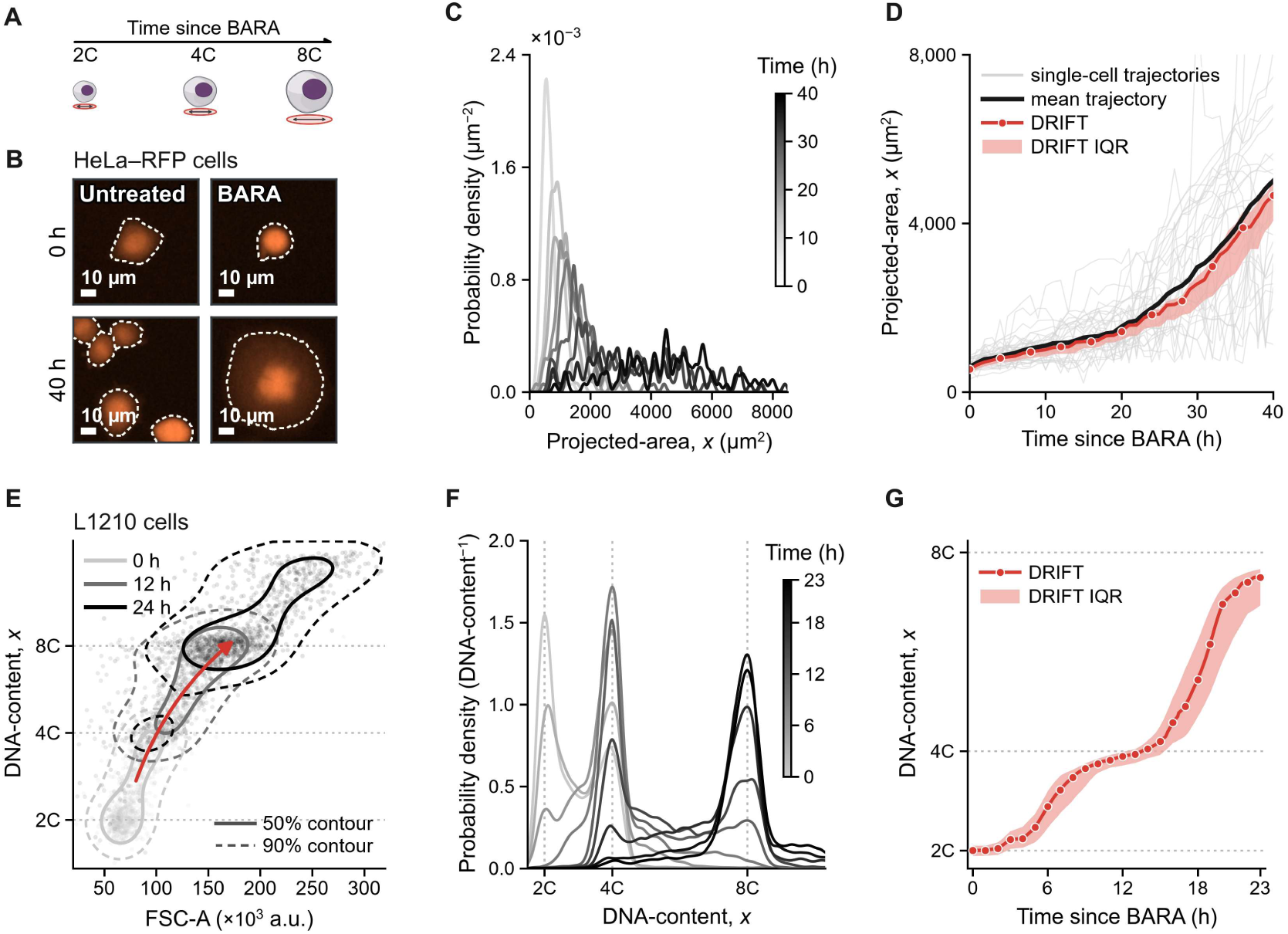
DRIFT extends to projected-area and DNA-content dynamics. (**A**) Illustration of cell size, projected-area and DNA-content changes after barasertib treatment. (**B**) Images of untreated and barasertib-treated (BARA) HeLa cells expressing free RFP at 0 and 40 h. Dashed lines mark segmented cell boundaries; scale bars, 10 µm. (**C**) Time-series projected-area probability densities *f(x,t)* calculated at each frame from the 100 retained complete tracks of BARA-treated HeLa cells. Data are shown for one representative replicate among the *n* = 5 biological replicates. Grayscale denotes treatment time. (**D**) Thirty representative single-cell projected-area trajectories (light gray), their mean trajectory (black), and DRIFT-inferred characteristic trajectories initialized at the mean (red) and the 25th and 75th percentiles (pink band) of the birth-state density at *t* = 0 h. All data are from the same replicate shown in panel **C**. Red circles mark positions at 4-h intervals. (**E**) Joint FSC-A and PI distributions from fixed-cell flow cytometry in L1210 cells treated with BARA for 0, 12, and 24 h. Solid and dashed contours enclose 50% and 90% of the plotted events, respectively. The red arrow indicates the direction of the population shift over time. (**F**) Time-series PI-derived DNA-content probability densities *f(x,t)* for BARA-treated L1210 cells from 0 to 23 h. Vertical guides denote the 2C, 4C, and 8C positions, and grayscale denotes treatment time. (**G**) Representative DRIFT-inferred DNA-content characteristic trajectories initialized at the mean (red) and the 25th and 75th percentiles (pink band) of the birth-state density at the time of perturbation (*t* = 0 h). Red circles mark positions at 1-h intervals. Panels **E**–**G** show data from one representative replicate among the *n* = 3 biological replicates.

## Results

### DRIFT infers cellular dynamics from time-series population measurements

DRIFT uses time-series population measurements to infer state- and time-dependent growth kinetics and the distribution of states at which cells divide, without tracking individual cells (Fig. 1A and B). At each time *t*, the inputs are the normalized state probability density *f*(*x*,*t*) of a measured state *x*, such as cell volume, projected-area, or DNA-content, and the total cell count *N*(*t*). Together, these define the time-dependent population distribution *n*(*x*,*t*) = *N*(*t*)*f*(*x*,*t*). Here, growth denotes progression along the measured state coordinate, whereas proliferation denotes changes in cell number.

Substituting *n(x,t)* = *N(t) f(x,t)* into the McKendrick–von Foerster population balance equation gives (refs. 37, 38; Fig. 1C):

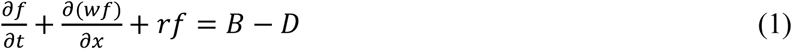

Here, *w*(*x*,*t*) is the velocity field, *r*(*t*) = (*dN*/*dt*)/*N* is the net population growth rate, and *B(x,t)* and *D(x,t)* are the daughter-birth and mother-division-loss densities, respectively. For binary division, *B* = 2*KD*, where the kernel *K* maps a mother state to the birth-state density (Supplementary Table 1).

Under zero transport flux at the lower state boundary, cumulative integration of equation (1) gives the following relation (Supplementary Note 1):

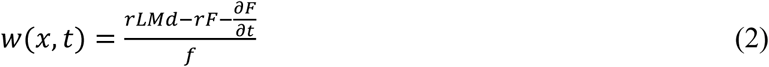

Here, *F(x,t)* = *Lf(x,t)* is the cumulative distribution, and *M* = 2*K* − *I*, where *L* is the integral operator and *I* is the identity operator. The lowercase source fields shown in Fig. 1 and equation (2) are the birth-state density *b*(*x*,*t*) = *B*(*x*,*t*)/*r*(*t*), and the unit-mass division-state density *d*(*x*,*t*) = *D*(*x*,*t*)/*r*(*t*), defined for *r(t)* > 0. In a balanced steady state, equation (2) reduces to the stationary relation underlying ERA if *f(x,t)* is time invariant (i.e., *f(x,t)* = *f(x)*), *N*(*t*) grows exponentially at a constant rate, and birth and division are confined to single boundary states (Supplementary Note 1).

In a balanced steady state, the velocity field and division-state density are time-independent functions, *w*(*x*) and *d*(*x*), respectively. Following a perturbation, DRIFT allows both fields to vary with state and time. However, the measured *f*(*x*,*t*) and *N*(*t*) alone do not uniquely determine how the observed redistribution is apportioned between progression and division^26,45^. To restrict the admissible solution space, we imposed two constraints: (i) non-divisional cell loss, such as cell death, was assumed to be negligible, allowing the net population growth rate derived from the measured cell counts, *N*(*t*), to constrain the total division flux; and (ii) the mother-to-daughter kernel linked the inferred distribution of mother-cell division states to the corresponding distribution of daughter-cell entry states.

Subject to these constraints, DRIFT fits the complete time-series population distributions *n*(*x*,*t*) jointly to estimate *w*(*x*,*t*) and *d*(*x*,*t*), while the count-derived *r*(*t*) sets the magnitude of *D*(*x*,*t*) (Fig. 1D–G). We used Tikhonov regularization to select biologically plausible and stable velocity fields and division-state densities consistent with the measured data and equation (2) (refs. 42, 46; Supplementary Note 1). The constrained formulation balances fidelity to the cumulative population-balance equation with smoothness of *w*(*x*,*t*) and *d*(*x*,*t*) across both state and time, thereby suppressing fluctuations without prescribing their shapes (Methods; Supplementary Note 1, Supplementary Fig. 1, and Supplementary Tables 1 and 2).

From the resulting velocity field *w(x,t)*, DRIFT reconstructs characteristic trajectories, *x̂*(*a*; *t*₀), where *x̂* is the state reached after an elapsed time *a* from an initial state *x*₀ at a start time *t*₀ (Fig. 1H; Supplementary Note 2). Each trajectory is obtained by integrating

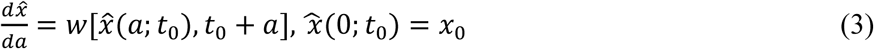

From a single inferred population-level velocity field, DRIFT generates characteristic trajectories initialized at different cellular states at the time of drug treatment. For cell volume, trajectories can be initialized at volumes corresponding approximately to newborn-like, mid-cycle, or near-division states (Fig. 1H). Comparing their subsequent volume growth allows DRIFT to estimate how the response to treatment depends on the initial cell volume. These trajectories are inferred from population measurements and should not be interpreted as directly tracked single-cell lineages or population-averaged single-cell trajectories. The starting-state labels describe relative positions along volume-growth progression rather than directly measured cell age. When the measured state coordinate approximately orders cell-cycle progression, these comparisons provide a way to infer cell-cycle-position-dependent responses without cell synchronization (Supplementary Note 2).

### DRIFT recovers growth kinetics in adder benchmarks

We first evaluated DRIFT using cell volume as the measured state coordinate. Volume can be measured directly and precisely at both single-cell and population levels, and models of cell growth and size homeostasis are well established^4,36,47–49^. We began with a deterministic asynchronous simulation in which every cell grew exponentially with the same volume-growth rate and divided at a single volume threshold. DRIFT reconstructed the characteristic volume trajectory with a relative root-mean-square error (RRMSE) of 0.62% and placed the inferred division-state density near the simulated threshold (Supplementary Notes 3 and 4, and Supplementary Fig. 2A–C).

Real proliferating populations exhibit cell-to-cell variation in growth history, instantaneous growth rates, birth size, and division size. Cultured mammalian cells show approximately log-normal size distributions and adder-like size homeostasis^4,12,36^. To model adder-like size homeostasis with cell-to-cell variability, we constructed a stochastic simulation (hereafter, the stochastic-adder population), in which the added-volume target, instantaneous growth rates, and daughter-cell volume partitioning varied among cells (Supplementary Note 3). At steady state, this population had an approximately log-normal volume distribution similar in range and shape to that measured for L1210 cells with a Coulter counter (Fig. 2A). The birth- and division-state densities of the simulated population spanned distinct volume ranges rather than forming sharp boundaries (Fig. 2B).

Using only the stationary volume probability density *f*(*x*) and the net population growth rate *r*(*t*) obtained from the total cell count *N*(*t*), DRIFT recovered birth- and division-state densities, *b*(*x*) and *d*(*x*), that closely matched the corresponding ground-truth distributions (Hellinger distance, 3.608 ± 0.007% and 7.787 ± 0.008% for the kernel-mapped birth and projected division-state densities, respectively; mean ± standard error of the mean (SEM), *n* = 10 independent simulations; Fig. 2B; Supplementary Fig. 3A and Supplementary Table 3). The DRIFT-inferred velocity field *w*(*x*) matched the linear velocity profile expected for exponential single-cell growth, in which the velocity increases in proportion to cell volume (Fig. 2C; RRMSE, 2.621 ± 0.003%; mean ± SEM, *n* = 10 independent simulations; Supplementary Fig. 3A and Supplementary Table 3). To examine how the representation of birth and division affects the inferred velocity field, we repeated the same steady-state inference using a fixed-boundary formulation in which daughter-cell birth and mother-cell division were each assigned to a single state (i.e., delta functions; Fig. 2B). Applied to the stochastic-adder population, the resulting velocity field deviated from the ground-truth profile, most prominently near the birth and division regions (Fig. 2C). This deviation follows from the boundary representation: the fixed-boundary formulation concentrates the entering and exiting fluxes at single volumes, whereas stochastic variation in birth size, added volume, and daughter-cell partitioning distributes birth and division across overlapping volume ranges (Fig. 2B; Supplementary Note 1). Under the steady-state assumptions used here, this fixed-boundary formulation is mathematically equivalent to ERA applied to a single scalar state coordinate^32^. However, this comparison specifically examined the consequence of the fixed-boundary representation in the stochastic-adder population. More generally, division events that are sharply defined in a higher-dimensional cellular state space may appear distributed when projected onto a single scalar coordinate.

The DRIFT-inferred characteristic trajectory matched the ground-truth mean trajectory of the simulated single-cell lineages when both curves were compared over one birth-to-division cycle on a common phase axis (Fig. 2D; RRMSE, 1.283 ± 0.095%; mean ± SEM, *n* = 10 independent simulations; Supplementary Note 4, Supplementary Fig. 3A, and Supplementary Table 3). To assess recovery beyond exponential single-cell growth, we repeated the same steady-state inference for three additional single-cell growth profiles—linear, sigmoidal Hill, and inverse exponential—under the same simulation settings. DRIFT recovered each growth profile and the corresponding characteristic trajectory in every case (Supplementary Note 3, Supplementary Fig. 4, and Supplementary Table 4), although distinguishing such growth modes is known to be statistically demanding in noisy single-cell measurements^50^.

We next assessed the effects of measurement noise and snapshot interval on recovery using the stochastic-adder population described in Fig. 2. We examined state-measurement noise in *f*(*x*,*t*) (e.g., volume-measurement noise from a Coulter counter), population-count noise in *N*(*t*), and the interval *Δt* between successive population measurements (hereafter, the snapshot interval; Supplementary Note 3). DRIFT was more sensitive to population-count noise than to state-measurement noise (Supplementary Fig. 5 and Supplementary Table 5). Across the tested noise levels, state-measurement noise left the trajectory RRMSE below 1.6%, whereas population-count noise increased it to 6.168 ± 1.074% at the highest noise level (Supplementary Notes 3 and 4, Supplementary Fig. 5C and G, and Supplementary Table 5). Population-count noise also amplified the effect of the snapshot interval: widening the interval from 0.4 to 4 h had little effect without population-count noise but increased the trajectory RRMSE by approximately 2.2-fold when population-count noise was present (Supplementary Notes 3 and 4, and Supplementary Fig. 6A). Thus, in these benchmarks, precise cell counting supported accurate trajectory recovery even at longer snapshot intervals, whereas shorter intervals improved recovery in the presence of population-count noise.

### DRIFT separates changes in volume-growth rate from changes in division size *in silico*

We next tested whether transient DRIFT inference could distinguish changes in volume-growth rate from changes in division size using two simulated perturbations. These perturbations represent two distinct changes relevant to drug-induced size responses. Translation inhibition can slow cell growth, whereas altered division control can shift the size at which cells divide^4,5^. Real drugs, however, can act through diverse molecular targets and combine both effects. Using the stochastic-adder population described in the preceding section (Fig. 2; Supplementary Fig. 7M), we simulated two cases in which either the volume-growth rate or the added-volume target was changed while the other was held fixed. In Case 1, the single-cell volume-growth rate was halved while the division-size rule was unchanged. The volume probability density *f*(*x*,*t*) remained nearly unchanged, whereas the total cell count *N*(*t*) increased more slowly (Fig. 3A–F; Supplementary Note 3). In Case 2, the added-volume target was increased 1.5-fold while the single-cell growth profile was unchanged. The population shifted toward larger cells, and the net population growth rate changed transiently as the population redistributed (Fig. 3G–L; Supplementary Note 3).

In Case 1, DRIFT recovered the decrease in the velocity field *w*(*x*,*t*) while the division-state density showed no appreciable shift, and the resulting characteristic trajectory retained its shape but progressed more slowly (Fig. 3C–F; Supplementary Fig. 7A–F and N). In Case 2, the inferred velocity field remained close to the control velocity profile while the division-state density shifted toward larger volumes, and the characteristic trajectory extended across the enlarged size range (Fig. 3I–L; Supplementary Fig. 7G–L and O). The characteristic trajectories were accurately recovered in both cases (RRMSE, 2.180 ± 0.163% and 3.502 ± 0.152% for Cases 1 and 2, respectively; mean ± SEM, *n* = 10 independent simulations; Supplementary Notes 3 and 4, Supplementary Fig. 3B and C, and Supplementary Table 3). The same inferred fields also generated distinct characteristic trajectories from newborn-like, mid-cycle, and near-division starting states (Fig. 3E and K; Supplementary Note 2).

We further varied the volume-growth rate and the added-volume target over broader ranges, both independently and in combination. Across these conditions, DRIFT continued to separate changes in volume-growth rate from changes in the division-state density and recovered the simulated characteristic trajectories (Supplementary Note 3 and Supplementary Fig. 8). We also examined how the snapshot interval, *Δt*, affected characteristic-trajectory recovery in Cases 1 and 2. DRIFT recovered the characteristic trajectories of both Cases 1 and 2 at all tested snapshot intervals, but the trajectory RRMSE increased as the interval widened in the presence of population-count noise (RRMSE, 9.18 ± 1.38% and 9.72 ± 1.99% for Cases 1 and 2, respectively, with population-count noise at 10% CV and 4-h snapshot intervals; mean ± SEM, *n* = 10 independent simulations; Supplementary Notes 3 and 4, and Supplementary Fig. 6B and C).

### DRIFT reconstructs condition-specific volume dynamics in L1210 cells

Having established DRIFT in simulation, we next asked whether it could recover transient volume dynamics from time-series population distributions of perturbed cells. We analyzed volume distributions and corresponding cell counts collected hourly from untreated L1210 mouse lymphocytic leukemia cells and cells treated with cycloheximide (CHX), nocodazole (NOCO), or barasertib (BARA), which perturb protein synthesis, mitotic progression, and cytokinesis, respectively (Fig. 4A–H; Supplementary Note 4). The CHX concentration was chosen to only partially inhibit protein synthesis so that cells continued to grow and divide at slower rates without completely arresting either process.

The DRIFT-inferred characteristic trajectories initialized at newborn-like volumes revealed distinct responses to each perturbation (Fig. 4I–L; Methods; Supplementary Notes 2 and 4). The characteristic trajectory for untreated cells showed two inferred divisions during the 17-h measurement window, whereas that for CHX-treated cells showed a single, delayed division. The first inferred division occurred approximately 40% later under CHX than in untreated cells (Supplementary Note 4). Under CHX, the inferred division volume also decreased over the measurement period, with the mean division state falling by approximately 10% between the first and last snapshots (Supplementary Note 4). Neither the newborn-initialized NOCO nor BARA characteristic trajectory shown exhibited a division event during the 17-h measurement window (Fig. 4K and L). Instead, the NOCO trajectory slowed and approached a plateau, whereas the BARA trajectory continued to increase throughout the measurement window. When initialized at near-division volumes, however, the NOCO and BARA trajectories each showed a single inferred division event (Supplementary Notes 2 and 4, and Supplementary Fig. 9). This dependence on initial state is consistent with stage-specific responses to both perturbations. Nocodazole treatment shortly after anaphase onset can permit furrow ingression and the formation of two daughter cells^51^, whereas Aurora B inactivation after furrow ingression can promote completion of cytokinesis through abscission^52^. Cells already sufficiently advanced through mitosis or cytokinesis at the time of treatment may therefore still complete one round of division. After this inferred division, the daughter trajectories showed the same qualitative responses as trajectories initialized at newborn-like volumes, with the NOCO daughter trajectory slowing toward a plateau and the BARA daughter trajectory continuing to grow without another division.

To compare the inferred volume dynamics with longitudinal single-cell measurements, we examined representative buoyant-mass trajectories measured with a suspended microchannel resonator (SMR) under the corresponding conditions (Fig. 4M–P). Because SMR measures buoyant mass rather than volume and the traces were obtained in separate experiments, these results provide an independent comparison of response direction and temporal form but do not test absolute agreement. The SMR-derived longitudinal single-cell traces showed response patterns similar to those of the DRIFT-inferred characteristic trajectories, including near-exponential accumulation in untreated cells, slower progression under CHX, a plateau-like response under NOCO, and sustained, near-exponential accumulation under BARA (refs. 6, 7; Supplementary Fig. 10).

### DRIFT extends to projected-area and DNA-content dynamics

We extended DRIFT beyond cell volume using the same population-balance framework with a mother-to-daughter kernel appropriate to each state coordinate. We first tested two other cellular state coordinates, projected-area and DNA-content, using deterministic benchmarks in an idealized cell model. In the projected-area benchmark, area scaled with cell volume and decreased upon division. In the DNA-content benchmark, DNA remained near 2C early in the cell cycle, increased sigmoidally to 4C, and returned to 2C in each daughter at division. For both state coordinates, DRIFT accurately recovered the velocity field *w*(*x*,*t*) and the characteristic trajectories (all RRMSEs < 3%; Supplementary Notes 3 and 4, and Supplementary Fig. 2D–I).

We next tested DRIFT using live-cell imaging of projected-area in adherent HeLa cells and fixed-cell flow cytometry of DNA-content in L1210 cells. Both state coordinates were examined in untreated steady-state populations and after BARA treatment. We selected BARA because Aurora B inhibition blocks cytokinesis while allowing continued cell growth and DNA replication, producing pronounced and sustained cell enlargement and DNA accumulation beyond 4C (ref. 7; Fig. 5A). BARA therefore provided a common perturbation with which to test whether DRIFT could reconstruct distinct physical and molecular responses from population distributions. We began with projected-area, analyzing population distributions obtained by imaging adherent HeLa cells expressing free red fluorescent protein (RFP; Fig. 5B). For untreated cells, we used the projected-area probability density *f*(*x*) measured at 20 h as the stationary input to DRIFT and compared the reconstructed characteristic trajectory with newborn-aligned trajectories obtained independently by direct single-cell tracking. The DRIFT-inferred characteristic trajectory agreed closely with the tracked cohort mean over the early 0–6-h window (RRMSE, 5.09%; Supplementary Note 4 and Supplementary Fig. 11E). After BARA treatment, the projected-area distribution shifted toward larger values over 40 h, and DRIFT reconstructed the corresponding sustained increase in projected-area (Fig. 5C and D; Supplementary Table 6). DRIFT-reconstructed characteristic trajectories captured the overall increase in the directly tracked mean trajectories of 100 retained cells per replicate (RRMSE, 10.87 ± 2.71%; mean ± SEM, *n* = 5 biological replicates; Supplementary Note 4, Supplementary Fig. 12, and Supplementary Table 7).

We then examined normal DNA-content progression in untreated L1210 cells. Using a steady-state analysis of propidium iodide (PI) distributions from fixed-cell flow cytometry, DRIFT reconstructed a closed DNA-content trajectory that progressed from 2C to 4C and returned to 2C at division (Supplementary Note 4 and Supplementary Fig. 13A–D). The state dependence of the inferred progression was independently evaluated using combined PI and 5-ethynyl-2′-deoxyuridine (EdU) measurements. Following separate unit-area normalization over the same DNA-content support, the EdU-incorporation profile and the stationary DRIFT velocity fields *w*(*x*) inferred separately from the PI series were closely aligned in shape and peak location (Pearson *r*, 0.975 ± 0.003; mean ± SEM, *n* = 3 biological replicates; Supplementary Note 4 and Supplementary Fig. 13E and F), supporting the state dependence of the DRIFT-inferred DNA-content progression.

Finally, we analyzed DNA-content progression in L1210 cells after BARA treatment. The measured population distributions shifted from the initial 2C and 4C peaks toward higher DNA-content and reached the 8C region (Fig. 5E and F). DRIFT reconstructed this progression as a staircase-like increase. DNA-content rose rapidly from 2C to 4C, slowed near 4C, and then resumed its increase toward 8C. This pattern was consistent with continued DNA replication without cytokinesis after Aurora *B* inhibition (ref. 7; Fig. 5G; Supplementary Note 4, Supplementary Fig. 14, and Supplementary Table 7). To evaluate the inferred kinetics independently, we compared the DRIFT-inferred velocity fields *w*(*x*,*t*) with estimates of DNA synthesis kinetics derived from Geminin and EdU measurements (Supplementary Fig. 15A and B). The DRIFT-inferred and independently estimated DNA kinetic profiles were positively correlated at both examined time points (Pearson *r* = 0.568 ± 0.016 at 11 h and 0.681 ± 0.151 at 22 h; mean ± SEM, *n* = 3 and 1 biological replicates for DRIFT-and Geminin-inferred profiles, respectively; Supplementary Note 4 and Supplementary Fig. 15C). Together, the projected-area and DNA-content analyses, supported respectively by direct tracking and independent estimates of DNA synthesis, show that DRIFT can infer cellular dynamics from both live-cell imaging and fixed-cell flow cytometry.

## Discussion

Many high-throughput assays quantify cellular states across large populations, but the resulting population distributions do not directly reveal how those states change within individual cells over time. DRIFT infers how an asynchronous, proliferating cell population progresses through a measured state coordinate without following the same cells over time. By representing state progression, mother-cell loss, and daughter-cell entry within a transient population balance, DRIFT captures kinetic information encoded in the evolution of cross-sectional distributions that is lost when the data are reduced to population means at each time point (Fig. 1).

DRIFT is particularly valuable for cellular quantities that are difficult or impossible to measure repeatedly in the same living cell. Using time-series flow-cytometry measurements of DNA-content in BARA-treated L1210 cells, DRIFT reconstructed DNA-content dynamics consistent with previous reports of continued DNA replication and polyploidization (ref. 7; Fig. 5E–G). Although several molecular reporters provide a proxy for DNA-content or synthesis dynamics, direct monitoring of absolute DNA amount or its synthesis rate over time in the same living cell remains challenging^53^. Moreover, prolonged exposure to live-cell DNA probes induces a DNA damage response and perturbs cell-cycle progression^22^. DRIFT therefore provides a way to infer time-resolved DNA-content dynamics under both unperturbed and perturbed conditions from conventional population endpoint assays without repeatedly measuring the same cells. DRIFT could also be extended to RNA, protein, lipid, or organelle-associated signals, provided that the measured state coordinate approximately orders the relevant cellular progression over the analysis window and the redistribution of that coordinate at division can be modeled. Recent steady-state ERA work using DNA and Geminin to order cell-cycle progression similarly inferred phosphoprotein dynamics and feedback from fixed-cell measurements^35^.

Because the inverse problem does not have a unique solution, DRIFT uses Tikhonov regularization to select a stable, representative velocity field and division-state density. This deterministic formulation is reproducible and computationally efficient, enabling the thousands of fits required for our benchmark and replicate analyses. However, the current implementation does not quantify uncertainty in the inferred field or distributions. Probabilistic extensions, including Bayesian inference, could provide uncertainty estimates but would require assay-specific models of measurement noise and biological stochasticity that may be difficult to verify^41,47,54,55^. Here, we instead assessed robustness by introducing known levels of noise into simulated measurements and repeating the complete analysis across independent simulation replicates (Supplementary Note 3). Extending DRIFT to quantify uncertainty while preserving its computational efficiency remains an important direction for future work.

Interpreting a DRIFT-inferred characteristic trajectory as representative of single-cell progression assumes that cells occupying the same state coordinate *x* at time *t* can be represented by a common conditional mean progression rate. Building on the population-to-lineage interpretation used by Kafri et al. for steady-state ERA^32^, DRIFT applies this assumption to transient, drug-perturbed populations. Accordingly, each trajectory describes the state progression implied by the population-level field from a chosen initial state, rather than a tracked or population-averaged single-cell path. Such a trajectory approximates lineage progression only when the population response is sufficiently coherent, with cell-to-cell variability appearing as deviations around the inferred trajectory. We evaluated this interpretation at several levels. In deterministic and stochastic-adder simulations, including those incorporating transient perturbations, DRIFT recovered the prescribed growth profiles and closely matched the corresponding mean simulated trajectories (Figs. 2 and 3). In measured populations, DRIFT resolved condition-specific volume responses that were consistent with single-cell buoyant-mass traces (Fig. 4), captured the overall projected-area progression relative to direct tracking of the same cohort, and inferred a DNA-content trajectory supported by independent measurements of DNA synthesis and ploidy (Fig. 5).

Yet DRIFT has several limitations arising from both the measurements and the current model formulation. First, DRIFT inherits the temporal limits of the input data. The total observation duration determines how long a response can be followed, whereas the measurement interval between successive snapshots sets the shortest response timescale that can be resolved. For example, a 1-h snapshot interval cannot capture the activation– adaptation dynamics of nuclear ERK2, which unfold within minutes^9^. In addition, because population growth enters the inference through *N*(*t*), sparse or noisy counts specifically distort the timing and magnitude assigned to division-related dynamics. Consistent with these limits, recovery degraded as snapshot intervals increased or *N*(*t*) measurement became less precise (Supplementary Note 3, Supplementary Figs. 5 and 6, and Supplementary Table 5).

Second, the current single-coordinate formulation assumes that the population responds coherently. DRIFT assigns one effective velocity and division profile to each state and time, and thus cells occupying the same measured state are summarized by a common population response. If drug-sensitive and resistant subpopulations coexist, or if distinct subpopulations respond at different onset times, their state progression and division dynamics are mixed in the observed distributions. If the measured state coordinate contains no information about subpopulation identity, resolving subpopulation-specific dynamics requires additional information, such as their proportions and expected response profiles. For example, if the resistant fraction were known independently and those cells were known to behave like untreated cells, their contribution could be separated using a two-class mixture model.

Third, DRIFT constrains the inferred velocity field *w*(*x*,*t*) to be nonnegative, and consequently, inferred progression cannot reverse along the measured state coordinate. Some cellular state coordinates nonetheless change nonmonotonically. For example, cyclin abundance rises and falls during subsequent recovery^56^. Such dynamics require negative velocity during decreasing segments and can cause the same coordinate value to be traversed in opposite directions. Because DRIFT assigns a single nonnegative velocity value to each state and time, decreasing segments can appear only as slowing or plateaus. Consistent with this limitation, the DRIFT-inferred projected-area characteristic trajectory in HeLa cells did not reproduce the transient decrease caused by mitotic rounding (ref. 57; Supplementary Note 3 and Supplementary Fig. 11E). Similarly, in simulations incorporating mitotic swelling^6,58^, DRIFT represented the resulting rise and rapid fall in cell volume as a smoothed plateau (Supplementary Note 3 and Supplementary Fig. 16).

Together, these considerations point toward a multiplexed DRIFT framework in which the state coordinate *x* of interest and additional variables are measured simultaneously in the same cells. For example, combining simultaneous viability measurements with absolute cell-counting standards could improve temporal inference by excluding dead cells and debris and providing more accurate estimates of *N*(*t*). Concurrent measurement of reporters that distinguish drug-sensitive and resistant subpopulations could enable subpopulation-conditioned or mixture-based inference, thereby resolving responses that are obscured in the aggregate population. Likewise, pairing *x* with a phase or state marker could distinguish cells that occupy the same value of *x* while progressing in different directions, supporting a multidimensional signed-transport formulation capable of recovering nonmonotonic dynamics. Thus, time-series multidimensional and multiplexed population endpoint assays could enable high-throughput drug-response screening that determines which subpopulations respond and through which kinetic mechanisms. More broadly, DRIFT is formulated in terms of the measured cellular state and its population balance, without specifying the molecular mechanism of a perturbation. The framework could therefore also be used to study how genetic perturbations or changes in nutrient availability affect state progression or division.

## Methods

### Basis of the DRIFT model

See Supplementary Note 1, for full details regarding the DRIFT model, including its mathematical derivations and numerical implementation. Briefly, observations from each condition were histogrammed on a state grid shared by all time points and normalized to the state probability density *f*(*x*,*t*); the total cell count *N*(*t*) was retained separately; for the experimental volume records, we used the fitted count curve described below. Interval growth rates were obtained from differences of log *N*. DRIFT then used the population balance in equation (1) and its cumulative form in equation (2), with binary division represented by *B* = 2*KD* and *D* = *rd*, where *d(x,t)* is a nonnegative unit-mass division-state density and *K* is a column-normalized single-daughter partition kernel following a symmetric Beta(15,15) daughter-fraction distribution for the volume analyses and a coordinate-specific map for the projected-area and DNA-content analyses. The transport flux at the lower boundary was set to zero.

The velocity field *w(x,t)* and division-state density *d(x,t)* were represented with piecewise-linear basis functions, using 64 velocity coefficients and 12 division coefficients defined on a common division support, and were estimated jointly over the full time series. The inverse was solved as a bound-constrained linear least-squares problem with *w* ≥ 0, *d* ≥ 0, and unit mass of *d* at every interval. Tikhonov penalties on the spatial and temporal derivatives of the two fields, together with a penalty on the spatial derivative of the specific rate *g* = *T₀w/x*, favored smooth fields and locally proportional growth without imposing an amplitude or a functional form. The penalty values were preset and identical across datasets. Each dataset was fitted twice, with a weak and a moderate specific-rate penalty. We retained the moderate fit only when its normalized residual exceeded the weak residual by no more than 25% in relative terms and 0.25 percentage points in absolute terms. Neither simulation truth nor experimental trajectories entered this choice. See Supplementary Note 1, Supplementary Fig. 1A–H, and Supplementary Tables 1 and 2, for the basis definitions, penalty values, and selection rule.

For the BARA-treated projected-area and DNA-content analyses, we assumed that no cell divided or was lost over the analysis window and set the birth, division, and count-source terms to zero (*r* = *B* = *D* = 0). For the projected-area analysis, the same retained tracks of viable, non-dividing cells were used to construct both the histograms for DRIFT inference and the directly tracked mean trajectory for comparison. For the DNA-content analysis, flow cytometry provided normalized DNA-content distributions, *f*(*x*,*t*), from the gated population at each time point, without corresponding absolute population counts, *N*(*t*). The coordinate-specific inference settings are given in Supplementary Note 4.

### DRIFT-inferred outputs

See Supplementary Note 2, for full details regarding DRIFT-inferred outputs, including numerical integration, division-event assignment, and characteristic trajectory initialization. Briefly, the inverse returned the population-level velocity field *w(x,t)* and the direct division-state density *d(x,t)*. We interpreted *w(x,t)* only over states covered by the measured distributions. For display and division-event assignment, we mapped every direct division-state density to the same fixed cubic B-spline projection targeted to the cumulative distribution function (CDF) and used the median of the projected density as the division-state criterion (Supplementary Fig. 1I–L, and Supplementary Table 1). We applied no Gaussian smoothing to inferred division curves. Simulated volume reference and event densities were smoothed with a mass-preserving Gaussian kernel (*σ* = 40 µm³) for display only. The primary division-state density error was computed between the fixed projection of the inferred density and the raw, unsmoothed simulated event density; the corresponding direct-density comparison was retained as an inverse-stage audit. See Supplementary Note 2, for the construction of the direct and projected outputs, and Supplementary Note 4, for the error metrics.

Characteristic trajectories were obtained by integrating the inferred velocity field from a specified starting state. The principal trajectory for each condition started at the mean of its inferred birth-state density. For the projected-area and DNA analyses, in which no division source was inferred, the trajectory instead started at the mean of a steady-state-proxy birth-state density inferred separately from the corresponding replicate- or series-specific 0-h distribution, and the plotted band was initialized at its 25th and 75th percentiles. For the mid-cycle and near-division starts in Fig. 3, the starting states were placed at phases 0.45 and 0.90, respectively, along the pretreatment birth-to-division trajectory. The pretreatment phase was preserved through *t* = 0: for initial phase *φ*₀, the first elapsed-time criterion used the remaining fraction (1 − *φ*₀)*T*_inst_(*t*), where *T*_inst_(*t*) denotes the instantaneous cycle duration, whereas subsequent cycles used the full *T*_inst_(*t*). See Supplementary Note 2, and Supplementary Table 1, for daughter-state initialization, pretreatment-phase assignment, and event assignment.

We assigned a division event only when two criteria were both satisfied: the elapsed time since treatment or the preceding reset reached the applicable phase-adjusted interval, and the trajectory reached the median of the projected division-state density. For the first event, this interval was (1 − *φ*₀)*T*_inst_(*t*); after a reset it was the full *T*_inst_(*t*). For the experimental volume records, the fitted count curve came from the same Coulter measurements used to define *N*(*t*). After an assigned event, the trajectory restarted at the condition-specific birth state with phase zero. We stopped the trajectory if the two criteria were not both satisfied before either the measured time range or the supported state range ended. See Supplementary Note 2, for numerical integration and division-event assignment, and Supplementary Note 4, for the experimental count-curve processing.

### Simulations

See Supplementary Note 3, for full details regarding simulation. Briefly, DRIFT was validated with simulated populations whose ground truth was not used during fitting and was compared with the inferred fields only afterward. Deterministic controls used an asynchronous population with an 8-h interdivision time, monotonic progression of volume, projected-area, and DNA-content, and coordinate-specific division resets (Supplementary Note 3). Stochastic-adder populations introduced cell-to-cell variability through an adder size-control rule with a mean added volume of 780 µm³, 5% variation of the added-volume target, multiplicative growth noise, and Beta(15,15) daughter partitioning, yielding an approximately log-normal steady-state volume distribution comparable to the measured L1210 Coulter distribution. All three conditions branched from the same steady-state population at *t* = 0. In Case 1, the single-cell volume-growth rate was halved at *t* = 0 while the division rule was unchanged. In Case 2, the added-volume target was increased 1.5-fold at *t* = 0, and the same rule was applied to subsequent daughters.

We varied the growth profile (Supplementary Fig. 4), basis resolution, regularization (Supplementary Fig. 1A– H), state-measurement noise, population-count noise, and snapshot interval. Recovery of the steady-state stochastic adder and Cases 1 and 2 was summarized as mean ± SEM across 10 independent simulations and complete refits; the four growth-profile benchmarks used 10 independent simulations per profile. Deterministic controls are point estimates. In the noise and snapshot-interval sweeps, every level was applied to the same 10 simulations, and the results are reported as mean ± SEM across those simulations. For the projected-area experiment, trajectory RRMSE was calculated separately for five biological replicates, each comprising 100 retained tracks, and reported as mean ± SEM across biological replicates (*n* = 5). See Supplementary Note 3, for the simulator definitions and perturbation construction, and Supplementary Note 4, for the error metrics and statistical units (Supplementary Tables 3–5).

### Cell culture and drugs

L1210 mouse lymphocytic leukemia cells (ATCC, CCL-219) were cultured in RPMI 1640 (Welgene, LM011-01) supplemented with 10% fetal bovine serum (Corning, 35-015-CF) and 1% penicillin–streptomycin (Gibco, 15140122) and were maintained at 37 °C in a humidified incubator with 5% CO₂. Before each experiment, cells were counted with a Multisizer 4e Coulter counter (Beckman Coulter) and diluted to 2 × 10⁵ cells/mL in fresh medium. The L1210 line expressing LifeAct-RFP and the FUCCI cell-cycle reporter was generated previously^16^ and was cultured under the same conditions. HeLa cells expressing free RFP from a CMV-driven pRFP-N1 plasmid with neomycin/kanamycin selection were cultured in DMEM (Welgene, LM001-05) supplemented with 10% fetal bovine serum (Corning, 35-015-CF) and 1% penicillin–streptomycin (Gibco, 15140122) and were maintained at 37 °C in a humidified incubator with 5% CO₂. For imaging experiments, HeLa cells were seeded in six-well plates at 10–20% confluency and were allowed to attach for 4 h before treatment.

For the Coulter volume experiments, L1210 cells were treated with 15 ng/mL cycloheximide (CHX), 1 µg/mL nocodazole (NOCO), or 100 ng/mL barasertib (BARA). For the DNA-content experiments, wild-type L1210 cells were treated with 100 ng/mL barasertib. For the projected-area experiments, HeLa cells were treated with 20 ng/mL barasertib. DNA was stained with FxCycle PI/RNase staining solution (Invitrogen, F10797). EdU was applied with the Click-iT EdU Alexa Fluor 647 Flow Cytometry Assay Kit (Thermo Fisher Scientific, C10419).

### Coulter counter volume measurements

Volume distributions were measured with a Multisizer 4e Coulter counter (Beckman Coulter) using a 100-µm aperture. Seven records acquired in this study were analyzed: one untreated, one cycloheximide, two nocodazole, and three barasertib files, each containing 18 nominal hourly snapshots. Finite positive diameters were converted to spherical-equivalent volume, *V* = π*d*³/6. For each replicate and time point, we removed debris and dead cells by placing a lower volume threshold at the valley between the low-volume peak and the cell peak of the histogram (Supplementary Fig. 17). Smoothing was used only to locate the valley. The threshold was then applied to the unbinned raw events, and no upper threshold was applied. Retained events from each replicate were converted to a density and averaged across replicates. To suppress snapshot-to-snapshot counting fluctuations in the interval growth rates, we represented the count record by a monotone penalized fit to log *N*(*t*), and the fitted curve was then combined with the retained fraction to give the *N*(*t*) used in the inference. The inference assumed that cell loss other than division was negligible over the fitted window. See Supplementary Note 4, for full details regarding data processing and statistics.

### Single-cell buoyant-mass measurements

Single-cell buoyant-mass trajectories were from previously published suspended microchannel resonator (SMR) measurements of L1210 cells^6,7^, acquired as described previously^13,14^, in which individual cells were passed repeatedly through the SMR and buoyant mass was calculated from the resonant-frequency shift. The cycloheximide and nocodazole data were published in *ref. 6*, and the untreated and barasertib data in *ref. 7*. These traces were compared with the volume-based characteristic trajectories only in terms of response direction and temporal form.

### Microscopy and imaging

Projected-area distributions were obtained from time-lapse images of the HeLa free-RFP cells. Five barasertib-treated biological replicates and one untreated acquisition were imaged in the orange fluorescence channel using an Incucyte SX5 (NFEC-2025-08-307698, Sartorius) with a 20× objective; 81 fields per biological replicate were acquired at 1-h intervals over a 0–43-h record; the reported analysis used the 0–40-h portion of the retained tracks. Cell masks were generated with the Cellpose cyto3 model^59^ and linked across frames within imaging fields that were sparse at 0 h and showed stable object motion. Tracks that spanned the whole record were then ranked separately within each biological replicate by a score that rewarded long, spatially stable trajectories with a sustained and internally consistent increase in area, and the 100 highest-scoring complete tracks from each of the five BARA-treated biological replicates were retained (500 tracks in total; Supplementary Table 6). See Supplementary Note 4, for image processing, track linking, track ranking, and biological-replicate repeatability. Each biological replicate’s retained tracks provided its own histograms for the corresponding inference and its own directly tracked mean for comparison. See Supplementary Note 4, for full details regarding data processing and statistics.

The fluorescence micrographs in Fig. 4A–D were acquired from the L1210 line expressing LifeAct-RFP and the fluorescent ubiquitination-based cell cycle indicator (FUCCI) at 0.137 µm per pixel on a ZEISS Apotome-equipped fluorescence microscope (Carl Zeiss Microscopy) with a 40× air objective (numerical aperture, 0.95) and structured-illumination optical sectioning. All conditions were acquired with identical illumination and exposure settings, and the same lookup table was applied to every displayed image of a given channel. These micrographs illustrate cell morphology and cell-cycle reporter state only and do not contribute to any DRIFT inference. See Supplementary Note 4, for full details regarding data processing and statistics.

### Flow cytometry

Wild-type L1210 cells were treated with barasertib and sampled at 1-h intervals over the 0–23-h record, giving three serial PI time series. At each time point, cells were fixed overnight in 70% ethanol at −20 °C, washed twice, stained in 50 µL of undiluted FxCycle PI/RNase staining solution for 30 min at room temperature in the dark, and analyzed immediately on a BD FACSymphony A5 SE flow cytometer (BD Biosciences) with BD FACSDiva v9.6, recording 10,000 events per sample. PI fluorescence was excited with the 561-nm yellow-green laser and collected in the YG585-A detector. DNA-content was quantified from PI fluorescence alone. The joint forward-scatter area (FSC-A)/PI distributions in Fig. 5E came from a separate PI acquisition at 0, 12, and 24 h in which forward scatter was recorded together with PI. Separate pretreatment PI datasets were used for the stationary DNA control (Supplementary Note 4 and Supplementary Fig. 13). PI–EdU data were used for independent validation of the stationary DNA field, whereas PI–Geminin data were used for the treated-series cross-modal consistency comparison; neither dataset entered DRIFT fitting. The PI–EdU dataset is described under EdU labeling below; see Supplementary Note 4, for the PI–Geminin dataset and transfer analysis, PI calibration, interpolation of missing samples, initial-state construction, and the inference settings. See Supplementary Note 4, for full details regarding data processing and statistics.

DNA synthesis rates were evaluated using EdU labeling with the Click-iT kit described above. Cells were labeled with 20 µM EdU for 30 min, washed twice with cold PBS, and fixed with 4% paraformaldehyde for 10 min. The cells were then washed with PBS, permeabilized with 0.1% Triton X-100 for 10 min, washed with PBS, and the incorporated EdU was labeled according to the supplier’s instructions. Following EdU labeling, the cells were stained with FxCycle PI/RNase. The PI–EdU data represent one untreated experiment with four technical replicates derived from two biological replicates (separate cultures), plus two EdU-negative controls. The PI–Geminin data comprise two experiments, one untreated and one barasertib-treated (Supplementary Note 4). The EdU comparison settings are described in Supplementary Note 4.

## Supporting information

Supplementary Informations

