## Supplementary Informations for "Division-resolved inference of flow and trajectories in proliferating cell populations"

**This Supplementary Information file includes:**

Supplementary Notes 1–4
Supplementary Figures 1–17
Supplementary Tables 1–7
Supplementary References

**Contents**

Note 1. Basis of the DRIFT model (pp. 2–5)
Note 2. DRIFT-inferred outputs (pp. 6–7)
Note 3. Simulations (pp. 8–10)
Note 4. Data processing and statistics (pp. 11–18)
Supplementary Figures 1–17 (pp. 19–35)
Supplementary Tables 1–7 (pp. 36–38)
Supplementary References (p. 39)

### Supplementary Notes

#### Note 1. Basis of the DRIFT model

**Snapshot preparation.** For each snapshot, we normalized the retained events to obtain the state probability density *f(x,t)*, written $f_q$ at snapshot time $t_q$. For the experimental volume records, retained events were those above the lower threshold defined in Supplementary Note 4. From the total cell count $N(t)$, with log denoting the natural logarithm, we calculated the net population growth rate of each snapshot interval $q$ as

$$r_q = \frac{\log N(t_{q+1}) - \log N(t_q)}{t_{q+1} - t_q},$$

and assigned the count-derived net population growth rate to that snapshot interval.

**Derivation of the cumulative form.** Let $n(x,t)$ be the population distributions of cells in state $x$ on the retained support $\Omega_x = [x_{\min}, x_{\max}]$, $N(t) = \int_{\Omega_x} n(x,t)dx$ the total cell count, and $f = \frac{n}{N}$ the state probability density. Let $w(x,t)$ denote the velocity field, $D(x,t)$ the mother-division-loss density, and $B(x,t)$ the daughter-birth density, each per cell per unit time and per unit state. The unnormalized balance is

$$\partial_t n + \partial_x(wn) = N(B - D).$$

Substituting $n = Nf$ and $r = \frac{\dot{N}}{N}$ gives

$$\partial_t f + \partial_x(wf) + rf = B - D.$$

Define $F(x,t) = \int_{x_{\min}}^{x} f(\xi,t)d\xi$. With zero flux at the lower state boundary, $w(x_{\min},t)f(x_{\min},t) = 0$, integration gives

$$\partial_t F + wf + rF = L(B - D).$$

For binary division with a single-daughter partition kernel $K$, $B = 2KD$. In a division-only proliferating population, $\int_{\Omega_x} Ddx = r$. We divide by this rate to define the unit-mass division-state density $d$ and the birth-state density $b$. This gives $D = rd$ and $B = rb$. Here $L$ integrates a field from $x_{\min}$ up to $x$, and the operator $M = 2K - I$, where $I$ is the unit (identity) operator, combines the two daughter births with mother loss, giving

$$wf = rLMd - rF - \partial_t F \qquad \text{(S1)}.$$

The $rf$ term is required because $f$ is normalized at every time. Omitting it, or imposing $\partial_t F = 0$ during a transient, reallocates population growth or movement of the distribution to $w$. Cumulative integration avoids direct spatial differentiation of noisy histograms, but temporal differentiation remains noise-sensitive.

On the finite grid, each column of the single-daughter partition kernel $K$ was normalized to carry the mass of exactly one daughter. This renormalization redistributed any daughter mass falling below the retained support edge rather than discarding it. The numerical $L$ integrated each field from the lower edge of the retained state grid up to each state, treating as zero any flux below the retained support edge $x_{\min}$. For the experimental volume records, the partition kernel followed a symmetric Beta(15,15) daughter-fraction distribution; the projected-area and DNA-content analyses used the coordinate-specific kernels described in Supplementary Notes 3 and 4.

For a spatial error field $\eta$, cumulative integration damps high-frequency components, scaling the Fourier magnitude approximately as $|\hat{\eta}(k)|/|k|$, which stabilizes the inverse, whereas differentiation amplifies it. Because two densities with similar cumulative distributions can still differ in peak position, width, or the number of peaks, we judged division recovery against the raw, unsmoothed density of recorded division events.

The closure $B = 2KD$ with $D = rd$ assumes that interval count change is dominated by binary division. Nondivisional loss or death that contributes to $\frac{dN}{dt}$ is not separately identifiable from snapshot densities and total counts and can bias the count-derived net population growth rate and the physical division-loss density. The inferred $D$ should be interpreted conditionally on the division-only closure, in which division dominates.

**Bases and scaling.** The inverse with the birth and division terms represented $w(x,t)$ with $K_w = 64$ spatial coefficients and $d(x,t)$ with $K_d = 12$ coefficients. On the division support $\Omega_D$,

$$\int_{\Omega_D} d_q dx = 1.$$

For display and for the division criterion in Supplementary Note 2, $d$ was additionally projected onto a smooth density $\tilde{d}$, defined in Supplementary Note 2. With $r_0$ denoting the reference net population growth rate, taken as the median of the interval rates, we expanded the velocity field in the nonnegative piecewise-linear (triangular) basis functions $H$ and the division-state density in the basis functions $\Phi$:

$$w_q(x) = r_0 x \sum_{j=1}^{K_w} \beta_{jq} H_j(x), d_q(x) = \sum_{j=1}^{K_d} \gamma_{jq} \Phi_j(x).$$

The $H_j$ were centered on the observed state-grid centers, and we call these centers the velocity basis points. If $r_0$ was nonpositive, we replaced it for scaling only by the median positive interval rate. Consequently, $\beta_{jq}$ and $\gamma_{jq}$ were dimensionless, whereas each $\Phi_j$ had unit discrete mass and units of inverse state. Coefficients obeyed

$$\beta_{jq} \geq 0, \gamma_{jq} \geq 0, \sum_j \gamma_{jq} = 1.$$

The population-balance equation entered the fit as a normalized least-squares residual rather than as an exact constraint.

The division basis used a common 200–4,000-µm³ physical support, corresponding to 210–3,990 µm³ on the 20-µm³ bin-center grid, for all seven volume records. This division support differs both from the range of the measured-state histogram and from the narrower range of states covered by the measured distributions, on which we scored $w$. The lower volume threshold was applied before the fit and did not change the common division support.

**Objective and regularization.** For adjacent snapshots, we define

$$\Delta t_q = t_{q+1} - t_q, \overline{f}_q = \frac{f_q + f_{q+1}}{2}.$$

The interval residual is

$$e_q = r_q L M d_q - w_q \overline{f}_q - L\left[\frac{f_{q+1} - f_q}{\Delta t_q} + r_q \overline{f}_q\right] \qquad \text{(S2)}.$$

With $r_0$ as defined above, we define

$$L_x = x_{\max} - x_{\min}, Q_x = \operatorname{diag}\sqrt{\frac{\Delta x}{L_x}}, \omega_q = \frac{\Delta t_q}{\sum_k \Delta t_k}.$$

The denominator that normalizes each interval residual is

$$s_q = \max\left\{\left\|Q_x\left[r_0 x\bar{f}_q + L\left(\frac{f_{q+1}-f_q}{\Delta t_q} + r_q\bar{f}_q\right)\right]\right\|_2, 10^{-10}\mathrm{h}^{-1}\right\} \quad \text{(S3)}.$$

The normalized variables are

$$\xi = \frac{x - x_{\min}}{L_x}, \tau = \frac{t}{T_0}, u = \frac{T_0 w}{L_x}, g = \frac{T_0 w}{x}, F_d(x,t) = \int_{x_{\min}}^{x} d(y,t)dy.$$

The minimized objective is

$$J = \sum_q \omega_q \|Q_x e_q\|_2^2 / s_q^2 + \lambda_{w,x} R_{w,x} + \lambda_g R_g + \lambda_{d,x} R_{d,x} + \lambda_{w,t} R_{w,t} + \lambda_{d,t} R_{d,t}.$$

The penalties act on values at these basis points. They are discrete derivative norms rather than classical $L_2$ derivatives. Let $\mathcal{D}^{(1)}_{\xi,w}$ and $\mathcal{D}^{(2)}_{\xi,w}$ denote nonuniform first- and second-derivative matrices on the $w$ basis points, let $\mathcal{D}^{(1)}_{\xi,D}$ act on the dimensionless division-state density $L_x d$ over the division support $\Omega_D$, and let $\mathcal{D}^{(1)}_{\tau}$ and $\mathcal{D}^{(2)}_{\tau}$ act on time values at the interval midpoints. Throughout, $\|\cdot\|_2$ denotes the Euclidean norm of a grid vector, $\|\cdot\|_F$ the Frobenius norm of a matrix, and $\langle\cdot\rangle_q$ the average over snapshot intervals; $\Delta x$ is the vector of state-bin widths and $T_0 = 1\,\mathrm{h}$ is the reference time scale. The diagonal matrices Q hold the integration weights: histogram-cell widths for terms on the state grid, the spacing between neighboring basis points for terms on those points, and the time weights $\omega_q$ for terms across intervals. The penalties are

$$R_{w,x} = \left\langle\left\|\mathrm{Q}_{w,2}\mathcal{D}^{(2)}_{\xi,w}\mathbf{u}_q\right\|_2^2\right\rangle_q, R_g = \left\langle\left\|\mathrm{Q}_{g,1}\mathcal{D}^{(1)}_{\xi,w}\mathbf{g}_q\right\|_2^2\right\rangle_q, R_{d,x} = \left\langle\left\|\mathrm{Q}_{D,1}\mathcal{D}^{(1)}_{\xi,D}L_x\mathbf{d}_q\right\|_2^2\right\rangle_q$$

$$R_{w,t} = \left\|\mathrm{Q}_{t,2}\mathcal{D}^{(2)}_{\tau}\mathbf{U}\right\|_F^2, R_{d,t} = \left\|\mathrm{Q}_{t,1}\mathcal{D}^{(1)}_{\tau}\mathbf{F}_d\right\|_F^2.$$

Here $\mathbf{u}_q$ and $\mathbf{g}_q$ are $u$ and $g$ at the $w$ basis points, $\mathbf{U}$ contains $u$ on the state grid across intervals, and $\mathbf{F}_d$ contains the division cumulative distribution function (CDF) on the state grid across intervals. The $w$ and $g$ penalties span the fixed state support $\Omega_x = [x_{\min}, x_{\max}]$, whereas $R_{d,x}$ spans only $\Omega_D$. The division penalty is the discrete integral of the squared first spatial derivative of the dimensionless division-state density, which is equivalent to penalizing the squared second derivative of its cumulative distribution on that support. Normalized time $\tau$ was evaluated at interval midpoints. The fixed settings were $T_0 = 1$ h, $\lambda_{w,x} = 10^{-6}$, $\lambda_{d,x} = 10^{-12}$, $\lambda_{w,t} = 10$, and $\lambda_{d,t} = 10^{-2}$. The histogram derivative in Supplementary equation (S2) was the forward difference between adjacent snapshots. The specific rate $g = \frac{T_0 w}{x}$ was evaluated for $x > 0$, with its lower-edge value defined by the finite basis limit when the support edge was zero. The inferred velocity field and division-state density were returned to physical units after solution.

**Choice between the weak and moderate penalties.** An additional penalty was placed on the spatial derivative of the dimensionless specific rate $g = \frac{T_0 w}{x}$. The specific-rate penalty is zero when $w$ is proportional to $x$, but neither fixes the proportionality constant nor prevents departures from exponential growth that the data support. Every dataset with the birth and division terms was solved twice: a weak fit with $\lambda_g = 3 \times 10^{-4}$ and a moderate fit with $\lambda_g = 3 \times 10^{-3}$. To compare the two fits, we measured how well each satisfied the population balance. For every interval we divided the residual of Supplementary equation (S2) by the size of the terms it balances, averaged this ratio over the intervals, and expressed it as a percentage:

$$\rho_q = \frac{\|e_q\|_2}{\max\left\{\left\|r_0 x\bar{f}_q + L\left[\frac{f_{q+1} - f_q}{\Delta t_q} + r_q \bar{f}_q\right]\right\|_2, \epsilon\right\}}, R = 100\langle\rho_q\rangle_q.$$

Here $\epsilon$ is machine precision, distinct from the fixed $10^{-10}$ $h^{-1}$ floor in Supplementary equation (S3). $R$ is therefore the mean population-balance residual in percent; across the four condition-level volume fits it ranged from 3.5% to 19.3% (Section Table 2). The interval average is unweighted and includes intervals with finite $\rho_q$. The $\frac{\Delta t_q}{\sum_q \Delta t_q}$ weights enter the fitted objective only, not $R$. Writing $R_{\text{weak}}$ and $R_{\text{mod}}$ for the two fits, we kept the moderate fit only when the moderate penalty raised the population-balance residual by at most one quarter in relative terms and by at most 0.25 percentage points in absolute terms:

$$\frac{R_{\text{mod}}}{R_{\text{weak}}} \leq 1.25, R_{\text{mod}} - R_{\text{weak}} \leq 0.25.$$

The penalty acts on the spatial roughness of $g = \frac{T_0 w}{x}$ rather than on the distance from a prescribed constant. Thus, $w = \alpha(t)x$ is the reference shape, with the amplitude *α(t)* set by the data. The fit keeps a nonexponential shape when it reduces the data residual by more than the smoothness penalty adds, and projected-area and DNA are therefore not forced to follow a volume-growth profile. The moderate coefficient is tenfold the weak one. The two conditions bind in different regimes: the ratio is the stricter test when $R$ is below 1%, and the 0.25-percentage-point cap is stricter above it. Thus, the cap governed for the measured records. The rule was applied only after both fits converged.

**Connection to ergodic rate analysis.** At balanced steady state, the state distribution is time invariant ($\partial_t F = 0$), and the population grows as $N(t) = N_0 e^{rt}$ with constant $r = \ln(2)/T_d$, where $T_d$ is the population doubling time. Supplementary equation (S1) then becomes

$$w(x)f(x) = r[LMd - F(x)].$$

ERA assumes fixed boundary states. A mother cell divides only at $x_d$, and both daughters enter at $x_b$. The corresponding division-state density and source mapping are

$$d(x) = \delta(x - x_d), \qquad [Kd](x) = \delta(x - x_b), \qquad [Md](x) = 2\delta(x - x_b) - \delta(x - x_d).$$

Let $H$ denote the Heaviside step function. Cumulative integration from the lower support boundary gives

$$[LMd](x) = 2H(x - x_b) - H(x - x_d).$$

Substitution into the stationary balance gives $w(x)f(x) = r\{2H(x - x_b) - H(x - x_d) - F(x)\}$. On the occupied cell-cycle interval $x_b < x < x_d$, the two step functions equal 1 and 0, respectively. The velocity therefore reduces to

$$w_{\text{ERA}}(x) = r\frac{2 - F(x)}{f(x)} = \frac{\ln 2}{T_d}\frac{2 - F(x)}{f(x)} \qquad \text{(S4)}.$$

Supplementary equation (S4) is the ERA occupancy–progression relation. DRIFT therefore reduces to ERA when the distribution is stationary, population growth is exponential, and birth and division occur at single boundary states.

**Note 2. DRIFT-inferred outputs**

**Division-event output and display.** The inverse estimated the direct unit-mass division-state density $d(x,t)$; the corresponding physical division-loss density $D = rd$ is defined in Supplementary Note 1. For display we evaluated $D$ with the projected density $\tilde{d}$ defined next. Because the fitted count curve does not decrease, the interval rate is nonnegative and $D$ is nonnegative. Each inferred division curve was mapped to a nonnegative cubic B-spline density $\tilde{d}(x,t)$ by fitting its cumulative distribution function with 24 coefficients and a smoothing penalty of 0.01. The projected division-state density was used as the plotted DRIFT curve and supplied the median division state used for event assignment. When simulated volume truth was shown, the raw 20-μm$^3$ event histogram was convolved with a mass-preserving Gaussian kernel with $\sigma = 40$ μm$^3$ for display only. We quantified division-state recovery with the Hellinger distance between the raw, unsmoothed simulated event density and the fixed projected inferred density.

**Condition-specific birth-state initialization.** For initialization, we ran a separate steady-state fit for each condition in which that condition's first distribution was held fixed. We repeated $f(x,0)$ over three snapshots one hour apart, imposed exponential growth at the matched pretreatment count rate, and applied the inverse defined in Supplementary Note 1. For the experimental Coulter records, $f(x,0)$ was the thresholded and renormalized density defined in Supplementary Note 4. We mapped the resulting division-state density through the single-daughter kernel $K$, renormalized the birth-state density, and initialized the central characteristic trajectory at the mean of the birth-state density. Each condition therefore used its own daughter state.

**Assignment of division events.** We fitted $\log N(t)$ for each condition with a monotone penalized curve and selected one common smoothness parameter by leave-one-out cross-validation pooled across the four conditions. The derivative of the fitted log-count curve defined the instantaneous count rate $r_{\text{fit}}(t)$ and the count-derived instantaneous cycle duration $T_{\text{inst}}(t)$. We assigned a division event only when the phase-adjusted elapsed-time threshold and the median of the projected division-state density were both reached. For a start at pretreatment phase $\varphi_0$ (defined below), the first threshold was $(1-\varphi_0)T_{\text{inst}}(t)$; after an assigned division event, the threshold was the full $T_{\text{inst}}(t)$.

$$r_{\text{fit}}(t) = \frac{d\ln[N_{\text{fit}}(t)]}{dt}, \quad T_{\text{inst}}(t) = \frac{\ln(2)}{r_{\text{fit}}(t)}.$$

The division-state boundary was the time-dependent median $x_{D,0.5}(t)$ of the projected unit-mass division-state density $\tilde{d}(x,t)$, the state at which its cumulative distribution first reaches 0.5.

For a birth-started path, $\varphi_0 = 0$. For a path initialized above the pretreatment birth-state mean $x_B$ at $t = 0$, the initial phase was the pretreatment travel time from $x_B$ to $x_0$ normalized by the full pretreatment birth-to-division travel time:

$$\varphi_0(x_0) = \frac{\left\{\int_{x_B}^{x_0} \frac{d\xi}{w_{\text{pre}}(\xi)}\right\}}{\left\{\int_{x_B}^{x_{D,\text{pre}}} \frac{d\xi}{w_{\text{pre}}(\xi)}\right\}},$$

where $w_{\text{pre}}$ is the pretreatment velocity field and $x_{D,\text{pre}}$ the pretreatment division-state median; first event when $a \geq [1-\varphi_0(x_0)]T_{\text{inst}}(t)$ and $\hat{x}(t) \geq x_{D,0.5}(t)$.

Each characteristic trajectory satisfies $\frac{d\hat{x}}{da} = \hat{w}[\hat{x}(a;t_0), t_0 + a], \hat{x}(0;t_0) = x_0$. The integration used 0.005-h Heun (RK2) steps that preserved nonnegativity, held the velocity constant within each interval, and interpolated linearly in state. We linearly resampled the dense solutions only for display and for the phase comparison with the simulations described below. After an assigned event, we returned the trajectory to the condition-specific birth state and set the phase to zero. Each later event required the elapsed time since the previous event to reach the full $T_{\text{inst}}(t)$ and the state criterion to be met. When the two criteria were not both satisfied before the record or supported state range ended, we truncated the trajectory and recorded that no division event had been assigned.

For synthetic comparisons, truth and inference used matched daughter starts and observation windows but reached their own division endpoints. We independently registered each complete truth and inferred birth-to-division trajectory to phase 0–1 and linearly resampled it to a common 1,001-point grid for the shape metric of the characteristic trajectory. We reported cycle-duration error separately. The simulator's adder target defined the simulation truth only and was not used to assign a DRIFT division event.

Mid-cycle and near-division starts corresponded to pretreatment phases 0.45 and 0.90, respectively, along the birth-to-division trajectory (Fig. 3E and K).

**Scope of application.** We applied the phase-adjusted temporal and state criteria to assign division events in all division-resolved volume characteristic trajectories, including steady-state and transient volume simulations, experimental volume analyses, and stationary-input controls (Figs. 2–4). We did not apply these criteria to the projected-area or transient DNA fits, which assumed no division, or to the measured suspended microchannel resonator (SMR) traces and independent 5-ethynyl-2′-deoxyuridine (EdU) and Geminin comparisons.

A trajectory can end without a division event because the phase-adjusted temporal criterion is not reached, the state criterion is not reached, or the supported state or time range ends first. Thus, the absence of an assigned division event indicates only that no division was inferred within the fitted record, not that division would never occur.

**Interpretation of inferred velocity fields and characteristic trajectories.** The reconstructed $w(x,t)$ is a population-level velocity that depends only on the measured state and time. A characteristic trajectory generated from the reconstructed velocity field is an inferred path rather than a tracked or population-averaged single-cell path. If the measurement cannot separate subpopulations that progress at different rates at the same measured state, one-dimensional snapshots recover only the combined flux of those subpopulations. Resolving such subpopulations requires an additional label or coordinate.

For the projected-area and transient DNA analyses, DRIFT inferred progression along the measured state coordinate under the assumption that no cell divided or was lost. The birth-state density from that steady-state fit was used only to initialize the central trajectory and its interquartile band; the birth-state density did not determine the transient velocity field. By contrast, the stationary-input controls tested whether the full DRIFT inverse remained internally consistent when the input distribution was held stationary (Supplementary Figs. 11 and 13).

**Note 3. Simulations**

**Deterministic benchmarks (Supplementary Fig. 2).** We used a deterministic asynchronous population with an 8-h interdivision time to test whether DRIFT reconstructs volume, projected-area, and DNA-content on their native coordinates. The balanced age density was $p(a) = 2r\mathrm{e}^{-r}$ for $0 \leq a < T$, with $T = 8$ h and $r = \frac{\ln(2)}{T}$. Normalized volume followed $V(a) = 2^{a/T}$, projected-area followed $A(a) = V(a)^{2/3}$, and terminal division applied $V \mapsto \frac{V}{2}$ or $A \mapsto 2^{-\frac{2}{3}}A$. These transformations give the analytic balanced distributions $f_V(V) = 2V^{-2}$ on $[1,2]$ and $f_A(A) = 3A^{-\frac{5}{2}}$ on $[1, 2^{2/3}]$.

The DNA-content control used a strictly increasing near-staircase profile designed to approximate long G1 and G2 periods of nearly constant DNA-content separated by rapid DNA synthesis. With normalized age $u = \frac{a}{T}$, $T = 8$ h, $\varepsilon = 0.08$, $\kappa = 18$, and $\sigma(z) = \frac{1}{[1+\mathrm{e}^{-z}]}$, we defined $L_\kappa(u) = \frac{\{\sigma[\kappa(u-\frac{1}{2})]-\sigma(-\frac{\kappa}{2})\}}{\{\sigma(\frac{\kappa}{2})-\sigma(-\frac{\kappa}{2})\}}$ and $x(u) = 1 + \varepsilon u + (1 - \varepsilon)L_\kappa(u)$. The positive $\varepsilon$ term keeps the velocity small but nonzero near 2C and 4C, whereas the logistic term concentrates most DNA accumulation near mid-cycle, giving a mid-cycle DNA-content velocity approximately 50-fold higher than near 2C and 4C. A zero-velocity plateau would give an infinite crossing time in this one-dimensional formulation; the positive $\varepsilon$ therefore preserves a finite birth-to-division trajectory. The balanced state distribution was generated exactly by transforming the same balanced age density through this monotone age-to-state map. In the simulation, cells divided at $x = 2$ and each daughter started at $x = 1$; the inverse was given only a broad division-state support and the deterministic halving kernel.

We applied DRIFT to hourly state distributions that were identical under balanced growth, together with an exponentially increasing total cell count, $N(t) = N_0\mathrm{e}^{rt}$. The velocity field $w$ and unit-mass division-state density $d$ were estimated jointly; the endpoint division state was not provided. The physical division loss was mapped to the daughter-birth density through the coordinate-specific deterministic single-daughter kernel. Both truth and inference were evaluated over the prescribed 8-h cycle. Supplementary Figure 2 reports the recovery errors for all three coordinates. DRIFT recovered the near-staircase DNA characteristic trajectory (Supplementary Fig. 2G–I).

**Stochastic-adder population (Figs. 2 and 3).** Figures 2 and 3 share a single simulated population, which we call the baseline population. Every cell advanced on a 0.2-h update grid according to $x(t + \Delta t) = x(t)\exp(k\Delta t + \eta)$, with $k = k_0 = \frac{\ln(2)}{8\text{ h}} = 0.08664\ \text{h}^{-1}$ and $\eta$ sampled independently at each update from a zero-mean Gaussian distribution with a standard deviation of 0.02. The mean growth profile is exponential, and the instantaneous growth rate varies from cell to cell and from update to update. At birth, each cell received an added-volume target with a mean of $\Delta_0 = 780\ \mu\text{m}^3$ and a relative standard deviation of 5%, and it divided at the first update for which the volume added since its birth reached that target. The added-volume target alone determined division; the absolute volume, the net population growth rate $r(t)$, and every DRIFT-inferred quantity were excluded from the simulation. At division we replaced the mother cell with two daughters whose volume fractions were $R$ and $1 - R$, where $R \sim \text{Beta}(15,15)$ was drawn independently at each division. Each daughter's volume became its birth volume, each daughter received an independent added-volume target, and the total cell count increased by one. The growth coefficient $k$ and the added-volume target are the only quantities that set the dynamics; the total cell count $N(t)$ and the net population growth rate $r(t)$ are outcomes of the simulated population and were supplied to DRIFT as data.

**Initialization and equilibration for the Figs. 2 and 3 baseline.** Each of the 10 simulations started from 2,000 cells. We drew each cell's initial volume from a Gaussian distribution with a mean of 780 $\mu\text{m}^3$ and a standard deviation of 100 $\mu\text{m}^3$, truncated below at 400 $\mu\text{m}^3$, set its birth volume to half of that volume, and assigned an added-volume target equal to that birth volume plus an independent draw with a mean of

780 µm³ and a relative standard deviation of 5%. This seed population only started the simulation and entered no analysis. We then advanced the population under the rules above until its volume distribution stopped changing. Every 2 h we histogrammed the volumes over 0–4,500 µm³ in 20-µm³ bins and compared the histogram with the one from the preceding check using the symmetric Kullback–Leibler divergence $D_{\mathrm{sym}}(p,q) = \frac{1}{2}D_{\mathrm{KL}}(p \parallel q) + \frac{1}{2}D_{\mathrm{KL}}(q \parallel p)$. We accepted the population as being at steady state at the first check for which $D_{\mathrm{sym}}$ stayed below $2 \times 10^{-4}$ on four consecutive comparisons. Across the 10 simulations this occurred between 78 and 84 h, within the 90-h limit we allowed.

**Prescribed single-cell growth profiles (Supplementary Fig. 4).** The stochastic-adder populations in Supplementary Fig. 4 used four single-cell growth profiles. Each profile replaced the exponential growth law of the baseline population while the initialization, equilibration, added-volume targets, and daughter partition were unchanged. With normalized cycle coordinate $u \in [0,1]$, birth volume $x_B = 780\ \mu\mathrm{m}^3$, added-volume target $\Delta = 780\ \mu\mathrm{m}^3$, and cycle time $T = 8\mathrm{h}$, volume was prescribed as $x(u) = x_B + \Delta y(u)$. The normalized profiles were $y(u) = 2^u - 1$ (exponential), $y(u) = u$ (linear), $y(u) = 0.10u + 0.90\frac{u^2}{u^2+(1-u)^2}$ (sigmoidal Hill), and $y(u) = \log_2(1+u)$ (inverse exponential). The velocity for each growth profile was $w(x) = \frac{dx}{dt} = \frac{\Delta}{T}\frac{dy}{du}$.

**Growth-rate and added-volume perturbations (Fig. 3).** The equilibrated population was recorded for a further 5 h before the perturbation, and this pre-perturbation record is shared by the three branches. Both cases branched from the baseline population at the perturbation time $t = 0$. We saved the current volume, the birth volume, and the remaining added-volume target of every live cell, together with the state of the random-number generator, and we started three branches from that identical point: the untreated baseline of Fig. 2, which ran for a further 18 h, and Cases 1 and 2, which each ran for a further 25 h. The three branches of one simulation begin from the same $f(x,0)$ and the same $N(0)$ and are therefore paired, whereas the baseline populations of different simulations are independent. In Case 1, the single-cell volume-growth rate was halved at $t = 0$, whereas the added-volume target and the division and daughter-partition rules remained unchanged ($k = 0.5k_0 = 0.04332\ \mathrm{h}^{-1}$; $\Delta = 780\ \mu\mathrm{m}^3$). In Case 2, the added-volume target was increased 1.5-fold at $t = 0$, whereas the single-cell volume-growth rate remained unchanged ($k = k_0$; $\Delta = 1.5\Delta_0 = 1{,}170\ \mu\mathrm{m}^3$). The Case 1 and Case 2 settings generated Fig. 3 and Supplementary Fig. 7 and are summarized in Supplementary Table 1, and their first-division times in Section Table 1 below.

**Section Table 1. First-division timing in the stochastic-adder simulations (mean ± standard error of the mean (SEM), *n* = 10 independent simulations).**

| simulated population | birth-state mean (µm³) | first assigned division (h) | simulated first division (h) |
|---|---|---|---|
| Steady-state stochastic-adder population | 807.559 ± 0.058 | 7.915 ± 0.002 | 8.110 ± 0.044 |
| Case 1: volume-growth rate 0.5× | 807.559 ± 0.058 | 15.607 ± 0.008 | 16.201 ± 0.100 |
| Case 2: added-volume target 1.5× | 807.559 ± 0.058 | 10.657 ± 0.001 | 10.714 ± 0.044 |

The Case 2 update multiplied each live cell's existing added-volume target by 1.5 ($\Delta_{\mathrm{post}} = 1.5\Delta_{\mathrm{pre}}$), preserving the existing cell-to-cell variation in the target. Daughters born after treatment received new added-volume targets with a mean of 1,170 µm³ and a relative standard deviation of 5%. Cell-specific target volumes and deterministic fixed points were not provided to the inverse. Instead, DRIFT used the time-varying median of the projected division-state density $\tilde{d}$.

The target update did not change cell states. The early rightward redistribution therefore reflected growth carrying cells to larger volumes together with a temporary reduction in daughter-cell entry, rather than an instantaneous shift in the state distribution or division-state density (Supplementary Fig. 7).

**Nonmonotonic-coordinate benchmark (Supplementary Fig. 16).** To isolate the limitation imposed by a nonnegative scalar velocity field, we generated a paired stochastic-adder population in which the latent size followed the growth and division rules of the main simulations while the measured volume carried a transient excursion. With latent size $z$, birth size $z_{\mathrm{birth}}$, division target $\theta$, and normalized log-size progress $\psi = \frac{\log\left(\frac{z}{z_{\mathrm{birth}}}\right)}{\log\left(\frac{\theta}{z_{\mathrm{birth}}}\right)}$, the measured volume was $x = z[1 + A_{\mathrm{exc}}\, h(\psi)]$, where the piecewise half-cosine $h(\psi)$ rose from zero over $\psi$ = 0.84–0.90 and returned to zero over $\psi$ = 0.90–0.97, and the benchmark shown in Supplementary Fig. 16 used $A_{\mathrm{exc}}$ = 0.12. Division was triggered by the latent target alone. The excursion therefore never advanced or delayed a simulated division, and the paired control without the volume excursion used identical latent sizes, targets, partitions, counts, and event times. The estimator was applied without retuning, and the mean of the 30 simulated lineages shown in Supplementary Fig. 16D, each followed from birth to its first division, provided the reference trajectory for the characteristic-trajectory comparison.

Cells on the increasing and decreasing branches can occupy the same measured volume. A single-valued single-cell velocity does not exist in that region. The two truth curves in Supplementary Fig. 16B therefore show velocities conditional on branch. The nonnegative DRIFT field should be interpreted as an effective population-level velocity whose characteristic trajectory is nondecreasing between division events.

The overall characteristic trajectory remained accurate, but DRIFT did not recover the localized late-cycle decrease because the inferred nonnegative field cannot represent both directional branches at the same measured state (Supplementary Fig. 16D).

The same limitation appears in the experimental projected-area records. Because cells round up before division, the tracked mean projected-area dips and rebounds near the end of the cycle, and the nonnegative DRIFT characteristic trajectory follows the net increase without reproducing this dip (Supplementary Fig. 11E). Recovering such nonmonotonic motion requires cell-cycle phase, a signed flux, or an additional state coordinate.

### Note 4. Data processing and statistics

**Coulter volume threshold selection and audit.** Each Coulter record contains a low-volume population of debris and dead cells in addition to the cell population. Following *ref. 1*, we removed this low-volume population by placing a lower volume threshold at the valley between the two peaks of the volume histogram. For every replicate and snapshot, we histogrammed the finite positive raw events in 20-µm³ bins over 0–6,000 µm³ and smoothed the histogram with a Gaussian kernel of $\sigma$ = 40 µm³. Smoothing was used only to locate the peaks and the valley. The cell peak was the tallest smoothed bin between 500 and 4,800 µm³, and the low-volume peak was the tallest smoothed bin below 500 µm³, or below 0.45 times the cell-peak volume when that value was smaller.

We searched for the valley strictly between the two peaks, excluding 80 µm³ ($2\sigma$) inward from each. The lowest smoothed bin in that interval defined the valley, with ties resolved toward the lower volume, and its bin center became the lower volume threshold. Events at or above the threshold were retained from the unbinned raw data (Supplementary Fig. 17A and B). Each replicate and time point received its own threshold, taken directly from its measured valley without smoothing or interpolation across snapshots. No upper threshold was applied, and the low-volume population was excluded on size alone, without an independent viability or debris label.

We required each valley to be well-resolved: the two peaks separated by at least 240 µm³, at least 8 smoothed counts in the cell peak, and a valley depth of at least 0.35. Valley depth is one minus the ratio of the smoothed valley count to the smaller smoothed peak count; a depth of 0.35 therefore means that the valley is 65% as high as the smaller peak. All 126 thresholds met these criteria and were retained. Supplementary Figure 17C and D and Section Table 2 show the variation of the selected thresholds across replicates.

**Section Table 2. Coulter counter lower-volume thresholds and population-balance fit by treatment condition.**

| treatment condition | selected lower threshold, min–max (µm³) | retained-event fraction, mean [range] | population-balance mismatch (%) |
|---|---|---|---|
| Untreated | 350–470 | 0.897 [0.883–0.912] | 3.639 |
| Cycloheximide | 270–390 | 0.886 [0.800–0.912] | 3.474 |
| Nocodazole | 310–930 | 0.794 [0.647–0.871] | 9.985 |
| Barasertib | 270–990 | 0.866 [0.781–0.931] | 19.257 |

**Volume density reconstruction and fit settings.** Volumes were spherical-equivalent volumes (Methods), binned on the same 20-µm³ grid used to choose the threshold. A small number of events fell outside the grid used by the inverse; these were kept in the raw records and still counted in the denominator when we measured the retained fraction. For each replicate and snapshot, we turned the retained events into a state probability density by dividing each bin count by the bin width and by the total number of retained events; each replicate density therefore integrates to one. We also recorded what fraction of all finite positive raw events survived the threshold. We then averaged the replicate densities, smoothed them across time with a five-snapshot Gaussian filter, set any negative values to zero, and renormalized.

The inverse also needs the total cell count $N(t)$. Rather than using the raw counts, we fitted a smooth curve to $\log N(t)$ that was not allowed to decrease (Supplementary Note 2). We then scaled fitted counts by the fraction of events that passed the threshold, averaged over replicates and smoothed with a three-snapshot moving median; $N(t)$ therefore refers to the same cells as the densities. We did this because the net population growth rate is the difference in $\log N$ between snapshots per unit time, and any fluctuation in the counts passes directly into the growth rate. In the simulations, fluctuations in the counts degraded recovery more than errors in the measured volumes did (Supplementary Fig. 5 and Supplementary Table 5).

The four conditions were fitted with the same basis functions and regularization settings, and with the same rule for choosing between the weak and the moderate penalty (Supplementary Note 1); that rule chose the moderate penalty in all four. Supplementary Figure 9 shows the inferred fields and the characteristic trajectories for each condition. Section Table 2 gives, for each condition, the range of thresholds selected, the fraction of events retained, and the mean residual $R$ of the population-balance equation. Section Table 3 gives the division timing for the untreated and CHX records and the change in division state under CHX.

**Section Table 3. CHX-induced changes in division time and division-state mean.**

| metric | reference | comparison | change |
|---|---|---|---|
| first division time | Untreated: 7.240 h | CHX: 10.055 h | +38.9% |
| division-state mean | CHX (0 h): 1,705 μm³ | CHX (17 h): 1,547 μm³ | −9.3% |

**Single-cell buoyant-mass traces.** Single-cell buoyant-mass trajectories were from previously published suspended microchannel resonator (SMR) measurements of L1210 cells, in which individual cells were passed repeatedly through the resonator and buoyant mass was calculated from the resonant-frequency shift[2,3]. We analyzed the published traces as recorded, without refitting or averaging. Because buoyant mass and Coulter volume were measured in separate experiments, the traces were used only to compare response direction and temporal form with the volume-based characteristic trajectories (Supplementary Fig. 10).

**Projected-area imaging and track analysis.** For projected-area, each biological replicate contributed 100 complete single-cell tracks of HeLa cells expressing free red fluorescent protein (RFP), selected as described below. Because the cohort was restricted by quality control to cells that were successfully segmented and followed in every frame, the same cells were present at every snapshot and no division or loss entered the cohort. The retained cell count was therefore constant by construction, and we set $r = B = D = 0$.

We generated masks with Cellpose cyto3[4] using a diameter of 85 pixels, a flow threshold of 0.8, a cell-probability threshold of −0.7, and a minimum object size of 160 pixels, and converted mask area to μm² using a pixel size of 0.625 μm. We restricted tracking to fields with no more than 20% confluency at 0 h and excluded objects that touched an image boundary or the scale bar and field-of-view label printed in the lower-left corner of each image or had projected-area outside 40–50,000 μm². Within each retained field, we linked objects one to one between consecutive hourly frames using centroid displacement and area change. Candidate links required a displacement of at most 70 pixels and an area ratio between 0.18 and 5.5; we accepted links greedily in order of increasing $C = \frac{\Delta r}{70} + \left|\log\left(\frac{A_{\text{current}}}{A_{\text{previous}}}\right)\right|$, allowing each object and each track to be used at most once per frame transition. We then checked how far cells moved within each field, using only tracks with at least 30 observations. A field was retained when it contained at least five such tracks, when the median of their mean frame-to-frame displacements was at most 16 pixels, when the 90th percentile of their maximum frame-to-frame displacements was at most 65 pixels, and when the median of their maximum displacements from the starting centroid was at most 150 pixels. Tracks from these fields that were present in all 44 hourly frames from 0 to 43 h formed a pool of 2,718 tracks for ranking.

We applied a five-frame trailing median to each complete projected-area track and ranked the tracks by a score $Q$ that combined seven measured properties of the track: $Q = 18\log_2[\max(F,1)] + 22\rho + 12P_+ + 0.04D_{\text{NN}} - 2\bar{D} - 18J_- - 10R_{90}$. Four terms reward the behavior we wanted to keep. $F$ is the maximum median-filtered area divided by the initial median-filtered area; tracks that grew more therefore score higher. $\rho$ is the Pearson correlation between time and median-filtered area, clipped to $[-1,1]$, and rewards a steady increase. $P_+$ is the fraction of consecutive area changes that are nonnegative, and rewards a smooth increase. $D_{\text{NN}}$ is the median nearest-neighbor distance, clipped to 0–300 pixels, and rewards cells that are well separated from their neighbors and therefore less likely to be confused when frames are linked. Three terms penalize signs of tracking failure. $\bar{D}$ is the mean interframe centroid displacement, which is large when a cell moves far between frames or the link is unstable. $J_{-}$ is the largest fractional decrease between consecutive median-filtered areas, which is large when a mask is lost or the track jumps to a different cell. $R_{90}$ is the 90th percentile of $\frac{|A-A_{\text{med}}|}{A_{\text{med}}}$, where $A$ is the measured area at a frame and $A_{\text{med}}$ is the five-frame median-filtered area at the same frame. This term is large when the measured area fluctuates from frame to frame. The coefficients are empirical weights fixed in the analysis code to place the terms on comparable numerical scales; they were not fitted or optimized against reference data. We ordered tracks by decreasing $Q$ within each biological replicate, resolved ties by ascending track identifier, and retained the 100 highest-scoring complete tracks from each replicate (Supplementary Table 6). The projected-area analysis used frames from 0 to 40 h, giving 100 observations at each of 41 hourly snapshots per replicate and 20,500 complete-frame observations in total. The score did not use any inferred field, characteristic-trajectory RRMSE, or agreement with DRIFT. Each retained cohort was therefore enriched for long, spatially stable trajectories with sustained, internally consistent area increase rather than being a random sample of segmented cells.

Within each replicate, these 100 tracks defined the time-series population distributions used to infer the velocity field and characteristic trajectory. Because there was no birth or division term to infer, the fit used the velocity basis alone, together with the spatial, temporal, and specific-rate penalties defined in Supplementary Note 1. We fitted the data with both the weak and the moderate penalty, and the selection rule retained the weak penalty ($\lambda_g = 3 \times 10^{-4}$). Supplementary Tables 6 and 7 report the retained cohorts and the corresponding fit results.

**Projected-area biological-replicate analysis.** We constructed the time-series population distributions and fitted the velocity field separately for each of the five biological replicates using 41 hourly snapshots from 0 to 40 h. For each replicate, we ran a separate steady-state fit as defined in Supplementary Note 2, using that replicate's measured $t = 0$ state probability density as the fixed input, an approximately 20-h HeLa free-RFP cycle to set the imposed count growth, and the projected-area division map as the daughter-state map. We used this proxy only to define the birth-state density and its mean, 25th-percentile, and 75th-percentile starting states.

We initialized the central projected-area characteristic trajectory at the mean of this birth-state density, and the shaded band spans the characteristic trajectories initialized at its 25th and 75th percentiles. We integrated each characteristic trajectory with the velocity held constant within each interval, linear interpolation in state, and 0.02-h Heun/RK2 steps, without post hoc smoothing. Replicate-level characteristic-trajectory RRMSEs and their mean ± SEM are reported in Supplementary Fig. 12 and Supplementary Tables 6 and 7.

**DNA-content inference from PI flow cytometry.** We reconstructed transient DNA-content dynamics from time-series PI distributions of BARA-treated L1210 cells, one from each biological replicate ($n$ = 3 biological replicates) (Supplementary Fig. 14). Sample preparation, staining, and acquisition are described in Methods. We located the 2C peak as the dominant mode of the PI fluorescence histogram and the 4C peak as the second mode near twice that intensity. PI fluorescence was mapped to the calibrated state coordinate by a single affine transformation whose offset and scale were chosen to place these two peaks at exactly 1 and 2; the offset is a calibration intercept determined by the two peak positions rather than a separately measured background. Within each file, a small residual peak-alignment shift constrained to at most 0.35 DNA/2C in absolute value was then applied. We report the calibrated state coordinate $x$ as DNA/2C, with the 2C, 4C, and 8C states at $x$ = 1, 2, and 4, respectively.

Missing snapshots in biological replicate #2 (19 h) and biological replicate #3 (6 and 20 h) were interpolated only to maintain numerical continuity and were not treated as observations. To initialize the trajectories, we ran a separate $t = 0$ steady-state fit as defined in Supplementary Note 2, using the initial state probability density as fixed input, an approximately 8-h L1210 cycle to set the imposed count growth, and a deterministic mapping from 4C to 2C as the daughter-state map. This proxy defined the birth-state density only. Flow cytometry recorded 10,000 events per sample, providing normalized DNA-content distributions. For the transient DNA analysis, we assumed that no cell divided or was lost over the fitted window and set $r = B = D = 0$. We therefore did not infer a division-state density or a post-division return to a daughter state. We fitted the data with both the weak and the moderate penalty separately for each biological replicate, and the selection rule retained the moderate penalty ($\lambda_g = 3 \times 10^{-3}$) for all three biological replicates (Supplementary Table 7).

For each biological replicate, we initialized the central characteristic trajectory at the mean of the birth-state density and the band boundaries at its 25th and 75th percentiles, then propagated all three starts through the velocity field of that replicate. Within each biological replicate, the band therefore represents dependence on the initial state rather than sampling uncertainty.

**Independent EdU validation in untreated cells.** For the untreated PI–EdU comparison, we processed each of the four acquisitions and the two EdU-negative controls separately; the data set is described in Methods. We first filtered events with a robust forward-scatter area (FSC-A)-versus-forward-scatter height (FSC-H) singlet gate and calibrated PI within each file by mapping the estimated 2C and 4C peaks to DNA/2C = 1 and 2. In each DNA bin, we calculated the EdU profile as the 10% trimmed mean of the linear signal in the allophycocyanin-area channel (APC-A) after subtracting the mean of the two negative controls in the same bin. We then averaged the four background-corrected EdU profiles with equal file weight. Finally, we normalized the resulting EdU profile and the stationary DRIFT velocity field of each of the three replicates separately to unit area on the same 0.90–2.14 DNA/2C support. We did not shift, rescale, or nonlinearly warp the DNA axis.

The equal-weight mean EdU profile and the pooled stationary DRIFT velocity profile shown in Supplementary Fig. 13F gave Pearson $r = 0.976$. Correlations with the stationary DRIFT velocity fields of the three replicates were 0.975 ± 0.003 (mean ± SEM, $n = 3$ biological replicates; Section Table 4). The EdU and $w(x)$ profiles peaked at 1.58 and 1.54 DNA/2C, respectively. The comparison does not determine how long cells spend in each phase, and it does not calibrate EdU fluorescence to the absolute magnitude of $w(x)$.

**Geminin-based cross-modal comparison in BARA-treated cells.** The PI–Geminin data comprise one untreated and one barasertib-treated experiment (Methods). Both were analyzed using the compensated channel values stored at acquisition, without additional compensation. We kept events with finite positive FSC-A, FSC-H, and side-scatter area (SSC-A) values, trimmed each scatter channel to its central 99.8% within each file, and selected singlets within 3.5 median absolute deviations (MAD) of an iteratively fitted $\log_{10}$(FSC-H)-versus-$\log_{10}$(FSC-A) line. PI and Geminin were not used for gating. Within each experiment, the dominant 2C peak and a second peak constrained to 1.65–2.25 times that peak were mapped by a linear rescaling to DNA/2C = 1 and 2; this calibration was fixed for the mixed acquisitions. Analyses used the common support 0.75–8.0 DNA/2C. Supplementary Figure 15A pools four acquisitions from the barasertib-treated experiment, each containing a mixture of samples collected at 0, 11, and 22 h after treatment, whereas Supplementary Fig. 15B and C use six separately labeled files from the same experiment (two at each treatment time).

To obtain an EdU-like readout for treated cells, in which EdU was not measured, we transferred the untreated PI–EdU relationship onto treated events using Geminin as an intermediate. Untreated measurements defined a PI–Geminin reference $G_0(u)$ and a PI–EdU reference $E_0(u)$ over a local 2C-to-4C cycle coordinate $u$. Each treated event lies in the 2C-to-4C, 4C-to-8C, or 8C-to-16C interval ($k$ = 0, 1, and 2, respectively). Before transfer, the total Geminin signal of each treated event was divided by $2^k$ to bring the low-Geminin branches of successive cycles onto a common baseline. For each barasertib-treated event with DNA state $x$ and normalized Geminin value $G$, the DNA-derived prior was $u_0 = \frac{x}{2^k}$, and the effective local coordinate $u^*$ minimized $(u-u_0)^2+\lambda[G_0(u)-G]^2$ with $\lambda=1$. The event-level EdU-equivalent signal was $\hat{E}=2^k E_0(u^*)$; the transferred signal therefore scales with DNA-content across successive replication cycles. We summarized these event-level predictions as 10% trimmed means in bins of width 0.12 DNA/2C, and line segments were not joined across replication-cycle boundaries.

For comparison with the transient DRIFT field, both $\hat{E}(x,t)$ and $w(x,t)$ were area-normalized over the DNA range covered at that time, and agreement was summarized by Pearson correlation and peak position (Supplementary Fig. 15C). The 0-h Geminin-based EdU-equivalent profile is the pretreatment reference. Because both reference curves came from untreated cells, and because the Geminin-based EdU-equivalent profile is not a direct EdU measurement in treated cells, the BARA profile comparison is a cross-modal consistency analysis rather than an independent validation.

The pooled PI–Geminin map in Supplementary Fig. 15A shows the recurring relationship between DNA-content and Geminin and was used only for descriptive visualization; it does not assign treatment time to individual events or measure velocity or DNA-synthesis rate.

For the time-specific comparison, we used the separately labeled 11- and 22-h files from the BARA-treated experiment. Within the experiment, we transformed Geminin with an inverse hyperbolic sine function and normalized it between the first and 99th percentiles for display, without comparing absolute fluorescence across experiments. Geminin did not enter DRIFT fitting or timing; in the transfer analysis it only refined the local cycle coordinate at which the untreated EdU reference was evaluated. Pearson $r$ was $0.568 \pm 0.016$ at 11 h and $0.681 \pm 0.151$ at 22 h (mean ± SEM across the three replicate DRIFT fields; Section Table 4), with a larger spread across replicates at 22 h. Each comparison used the DRIFT field of each of the three replicates and one Geminin-based EdU-equivalent profile at the matched time.

**Section Table 4. Replicate-level Pearson correlations for DNA cross-modal comparisons.**

| comparison | time | replicate #1 | replicate #2 | replicate #3 | mean ± SEM |
|---|---|---|---|---|---|
| Untreated stationary EdU versus DRIFT velocity | — | 0.977 | 0.968 | 0.979 | 0.975 ± 0.003 |
| BARA EdU-equivalent versus DRIFT velocity | 11 h | 0.579 | 0.590 | 0.536 | 0.568 ± 0.016 |
| BARA EdU-equivalent versus DRIFT velocity | 22 h | 0.886 | 0.772 | 0.386 | 0.681 ± 0.151 |

**Stationary-input controls.** We used two controls to test whether the inverse remained internally consistent with the measured state probability density fixed in time. For projected-area, we used a single untreated well and pooled the objects segmented and tracked in its 81 imaging fields, which are not independent replicates. The projected-area probability density at 20 h was held fixed and repeated at every time point (Supplementary Fig. 11A). We built $N(t)$ by counting all objects that passed quality control at each hourly time point in the same well (Supplementary Fig. 11B). We used the 10–40-h window, excluding the first 10 h because the cells had only just been seeded, and took the 10-h time point as the internal origin, giving 31 time points on a 0–30-h axis. The counts rose from 1,957 objects at 10 h to 5,261 at 40 h. Only the stationary-input projected-area control used the empirical partition kernel measured from paired daughter and parent projected-areas.

For DNA, we held the pooled pretreatment 0-h state probability density fixed from 0 to 8 h, imposed an 8-h doubling of $N(t)$, and mapped each 4C mother cell deterministically to a 2C daughter cell using a fixed one-half mapping rather than a Beta partition kernel (Supplementary Fig. 13A). The pooled density was built from three separately processed PI/RNase acquisitions as described below.

We fitted each control with both the weak and the moderate penalty and applied the Supplementary Note 1 selection rule separately for each coordinate. We projected each inferred division-state density with the CDF-targeted procedure defined in Supplementary Note 2. In both controls, the mean of the inferred birth-state density defined the initial state, and the characteristic trajectory used the elapsed-time and division-state criteria defined for the volume analysis. In both controls, the median division-state criterion was satisfied before the count-derived elapsed-time criterion. Each control was fitted once, on a single input distribution, and therefore gives one set of outputs rather than a replicate-level mean (Section Table 5). These controls do not provide measured longitudinal trajectories: the DNA count curve was imposed rather than measured, and the tail of the projected-area division-state density remained sensitive to the state boundary.

The pretreatment DNA densities were processed with the same PI/RNase pipeline as the main DNA analysis. After scatter gating, we evaluated event counts on a 900-point grid over 0.75–8.0 DNA/2C and convolved the counts with a fixed Gaussian kernel of standard deviation 0.06 DNA/2C. For the stationary-input control, we restricted each of the three 0-h densities to 0.75–2.20 DNA/2C, renormalized them, and averaged them into the single pooled density used as the input. Supplementary Figure 13A shows the histogram of gated events on fixed bins together with the curve supplied to the inverse; that curve received no further display smoothing.

We compared the projected-area characteristic trajectory from this control, taken as fitted, with a newborn-aligned cohort of 29 cycles defined independently of DRIFT, which gave a common-window RRMSE of 5.09% over the prespecified 0–6-h window (Supplementary Fig. 11E). Section Table 5 summarizes the timing and state outputs from both controls.

**Section Table 5. Timing and state outputs from stationary-input controls for projected-area and DNA-content.**

| **stationary-input control** | **count-derived cycle period / first assigned division (h)** | **birth-state mean** | **division-state median** | **characteristic-trajectory fold change over one cycle** |
|---|---|---|---|---|
| Projected-area, μm²: untreated 20-h snapshot | 21.678 / 21.678 | 576.890 | 1079.310 | 1.9305 |
| DNA-content, DNA/2C: pooled pretreatment density | 8.000 / 8.000 | 0.959 | 1.924 | 2.1064 |

**Error metrics.** We calculated velocity RRMSE over the central 90% of the *f(x,t)* probability mass at each time, with equal weights across the retained state bins:

$$\mathrm{RRMSE}_w = 100\sqrt{\frac{\sum_t \sum_{x\in S_{90}(t)}[\hat{w}(x,t) - w(x,t)]^2}{\sum_t \sum_{x\in S_{90}(t)} w(x,t)^2}},$$

Here, $\hat{w}(x,t)$ and $w(x,t)$ denote the inferred and simulated velocity fields, respectively, and $S_{90}(t)$ is the central 90% state support at time $t$. For each snapshot interval, we then compared the inferred and simulated division-state densities with the Hellinger distance, calculated as

$$H_d(t) = 100\sqrt{\frac{1}{2}\int\left[\sqrt{\tilde{d}(x,t)} - \sqrt{d_{\mathrm{event}}(x,t)}\right]^2 dx},$$

where $\tilde{d}(x,t)$ is the projected inferred division-state density and $d_{\mathrm{event}}(x,t)$ is the raw, unsmoothed simulated event density. We averaged $H_d(t)$ over the fitted snapshot intervals to obtain the projected Hellinger distance. We separately calculated the Hellinger distance from the direct inferred $d(x,t)$ before projection as an inverse-stage audit. For complete synthetic cycles, whose inferred and simulated cycle durations differ, we rescaled the simulated and the inferred birth-to-division trajectory each to phase $u \in [0,1]$ before comparison. We calculated characteristic-trajectory RRMSE as

$$\mathrm{RRMSE_{char}} = 100\sqrt{\frac{\sum_u [\hat{x}(u) - x(u)]^2}{\sum_u x(u)^2}},$$

where $\hat{x}(u)$ and $x(u)$ are the inferred and simulated characteristic trajectories. We calculated cycle-duration error as

$$E_T = 100\frac{|\hat{T} - T|}{T},$$

where $\hat{T}$ and $T$ are the inferred and simulated cycle durations, respectively. For experimental trajectories and for any trajectory without a complete division event, we could not define a full cycle and instead compared the two curves on the fixed absolute-time grid over the window in which both were defined, without time warping. We call this the common-window RRMSE.

To check how much the result depended on the choice of support, we recalculated the velocity RRMSE over the central 98% of the probability mass and over a fixed support. The Hellinger distance calculated from the fixed projected $\tilde{d}$ was the division-state recovery metric. The projection coefficients (24) and smoothing penalty (0.01) were fixed in advance and were not selected from the recovery errors of individual cases.

For a complete synthetic cycle we report a characteristic-trajectory RRMSE and a separate cycle-duration error. For a trajectory without a complete division event we report a common-window RRMSE, and its duration is reported as not estimated. A trajectory that left the state support of $f$ was not counted as a valid fit.

**Repeatability and independent units.** Simulation repeatability was evaluated with 10 independent simulations, each of which generated the steady-state population, Case 1, and Case 2. For every simulation we used a new random seed and reran the entire workflow: snapshots, counts, inverse, penalty selection, division outputs, and characteristic trajectories. The independent unit was the simulation ($n$ = 10), not simulated cells, lineages, histogram bins, or time points (Supplementary Fig. 3). In experiments, the independent unit was the biological replicate.

We report estimates as mean ± SEM across these independent units. Supplementary Tables 3 and 4 report recovery metrics for the steady-state population, Cases 1 and 2, and the four prescribed growth profiles.

**Sensitivity and noise sweeps.** We varied six estimator settings: the specific-rate penalty; the velocity and division-density basis sizes; the projection applied to the division-state output; the quantile of the projected division-state density used as the division criterion; how the birth state was initialized; and the settings used when no division term was inferred. Supplementary Figure 1 and Supplementary Table 2 report the tested grid, the number of valid fits, and the recovery metrics.

All simulation sweeps used the error definitions above. In Supplementary Fig. 5 and Supplementary Table 5 we varied the two noise sources separately: error in the measured state probability density $f(x,t)$ with exact counts, and noise in the total cell count $N(t)$ with exact state probability densities, at coefficients of variation (CVs) of 0, 1, 2.5, 5, 10, and 15%. Every noise level was applied to the same simulations, giving 10 paired simulations per level. At every CV, including CV = 0, the counts were passed through the monotone log-count fit described below before inverse estimation. We added the state-measurement noise by convolving $f(x,t)$ with a Gaussian kernel, and we did not additionally subsample to a finite number of events.

For Supplementary Fig. 6, we sampled exact subsets of the same frozen 0.2-h latent histories at 0.4-, 1-, 2-, and 4-h snapshot intervals and crossed these intervals with count-noise CVs of 0, 1, 2, 5, and 10% for the steady-state population, Case 1, and Case 2. Noise was applied as $N_{\text{obs}}(t) = \max\{1, N_{\text{true}}(t)[1 + \text{CV}\, z_t]\}$. Within each condition and simulation, one master noise vector $z_t$ was scaled across CVs and exactly subset across snapshot intervals, preserving pairing across all cells. No additional measurement noise was added to $f(x,t)$. As in Supplementary Fig. 5, the counts at every CV, including CV = 0, were passed through the monotone log-count fit before inverse estimation and characteristic integration.

Before inverse estimation, we obtained a single nondecreasing fitted log-count curve $g$ by minimizing $\|g - \log N_{\text{obs}}\|_2^2 + \lambda_{\text{count}}\|D_2 g\|_2^2$ subject to $\text{diff}(g) \geq 0$ . The count-fit penalty was not retuned: $\lambda_{\text{count}}(\Delta t) = 10^{-0.75}(1\,\text{h}/\Delta t)^4$ . Fitted counts were $\exp(g)$ , and inverse interval rates were $\max[\text{diff}(g)/\Delta t, 0]$. The instantaneous characteristic clock used a PCHIP representation of this same fitted log-count curve and its nonnegative derivative; no second count fit was performed. The inverse-specific-rate Tikhonov mapping was also held fixed: the steady-state baseline used the weak value $\lambda_g = 3 \times 10^{-4}$, whereas Case 1 and Case 2 used the moderate value $\lambda_g = 3 \times 10^{-3}$. We did not reselect $\lambda_g$ for individual count-noise/snapshot-interval cells. Projected division-state density was primary; direct division-state density was retained as an audit.

For each primary, birth-proxy, and alternative-regularization fit, we ran two cold-start constrained least-squares solves with identical physical constraints. Scaling the complete objective system by $\beta$ changes the squared objective scale by $\beta^2$ without changing its mathematical minimizer. The $\beta = 100000$ solve was the frozen production solve, and $\beta = 10000$ was a mandatory independent convergence audit; manuscript metrics always came from $\beta = 100000$. Both solves had to terminate successfully, satisfy the configured physical equality and bound tolerances, and agree within the configured objective, coefficient, velocity-field, division-density, and primary-metric thresholds. Any failure suppressed manuscript-facing output, and no fallback solver was used.

Each of the 12 division-density basis columns was normalized to unit discrete mass, and its nonnegative coefficients were constrained to sum to one independently in every interval. Before optimization, each equality row was divided by its Euclidean norm, $\sqrt{12}$. The internal constrained-least-squares tolerance was $10^{-10}$, matching the Tikhonov and isolated-noise analyses. Thus, the corresponding physical-coordinate equality-residual envelope was $\sqrt{12} \times 10^{-10} = 3.4641 \times 10^{-10}$; we used a rounded, scale-aware physical QA ceiling of $5 \times 10^{-10}$. This ceiling was an acceptance check only and did not enter the objective or alter any inferred field.

For Supplementary Fig. 8 we started from the same steady-state population and crossed volume-growth-rate multipliers of 0.5, 0.8, 1, 1.2, and 1.5 with added-volume-target multipliers of 0.8, 1, 1.2, and 1.5, simulating and refitting each of the 20 combinations independently. The heat maps of Supplementary Figs. 6 and 8 report the mean ± SEM across simulations. All 600 cadence-by-count-noise design rows and all primary, birth-proxy, and alternative-regularization fit pairs passed the prespecified checks, so every Supplementary Fig. 6 cell contains 10 valid paired simulations.

**Experimental and cross-modal comparisons.** We compared projected-area characteristic trajectories with the directly tracked mean of the same retained cells, using the common-window RRMSE defined above. None of the experimental comparisons — projected-area, SMR, untreated PI–EdU, and BARA PI–Geminin — entered DRIFT fitting. For the cross-modal comparisons we calculated Pearson correlation over DNA-state bins after normalizing each profile separately to unit area on a common support. Because the two signals do not share an absolute scale, the coefficient measures profile-shape consistency rather than pointwise prediction accuracy. In the steady-state comparison we used the 32 bins for which the EdU profile and the interpolated $w(x)$ were finite. In the barasertib comparison we used the DNA support shared by the two profiles at the matched time. The three biological-replicate DRIFT fields were the independent units for the mean ± SEM reported in the main text, and the data and transfer model are described under Independent EdU validation in untreated cells and Geminin-based cross-modal comparison in BARA-treated cells.

### Supplementary Figures

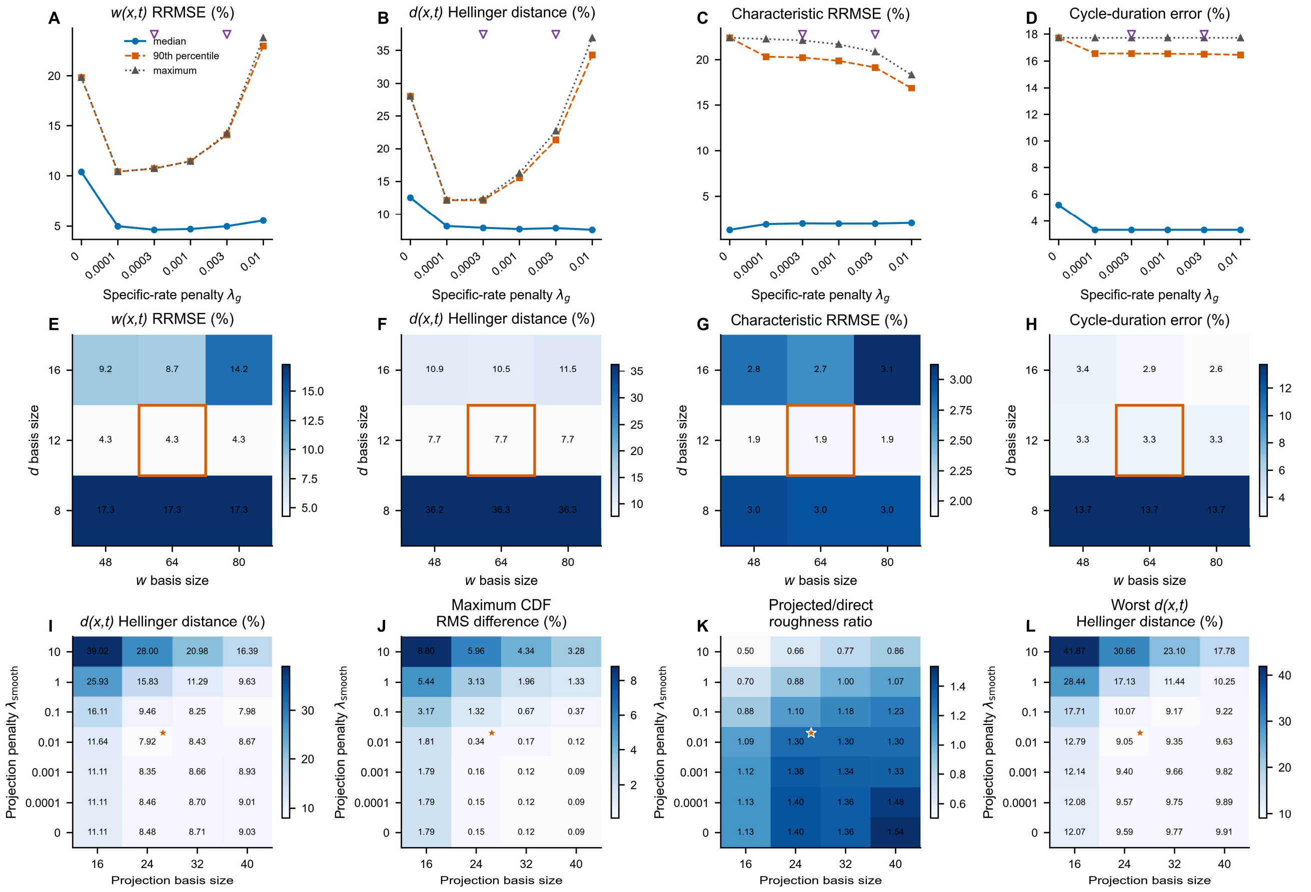


**Supplementary Fig. 1. Recovery errors are stable across the regularization, basis, and projection settings tested.** (**A**–**D**) Velocity field *w(x,t)* RRMSE, division-state density *d(x,t)* Hellinger distance, characteristic-trajectory RRMSE, and cycle-duration error, respectively, as the specific-rate penalty $\lambda_g$ was varied. Curves show the median (blue circles), 90th percentile (orange squares), and maximum (gray triangles) across four simulated growth profiles and Cases 1 and 2, and purple inverted triangles mark the weak and moderate penalties. (**E**–**H**) Median errors of the same four metrics for combinations of 48, 64, or 80 velocity coefficients and 8, 12, or 16 division coefficients; orange outlines mark the 64/12 basis used in the final analysis. (**I**) Interval-averaged Hellinger distance of the projected division-state density against the raw simulated event density, averaged over the three simulated conditions. (**J**) Largest interval-wise root-mean-square difference between the projected and the direct inferred cumulative distributions across the three conditions. (**K**) Ratio of projected to direct roughness, averaged over the three conditions, where roughness is the summed absolute second difference divided by the summed absolute first difference. (**L**) Largest mean Hellinger distance among the three conditions. In **I**–**L**, the axes are 16, 24, 32, or 40 projection coefficients and smoothing penalties from 0 to 10, and stars mark the selected setting of 24 coefficients and 0.01. Supplementary Table 1 lists the fixed settings, and Supplementary Table 2 reports the numerical values.

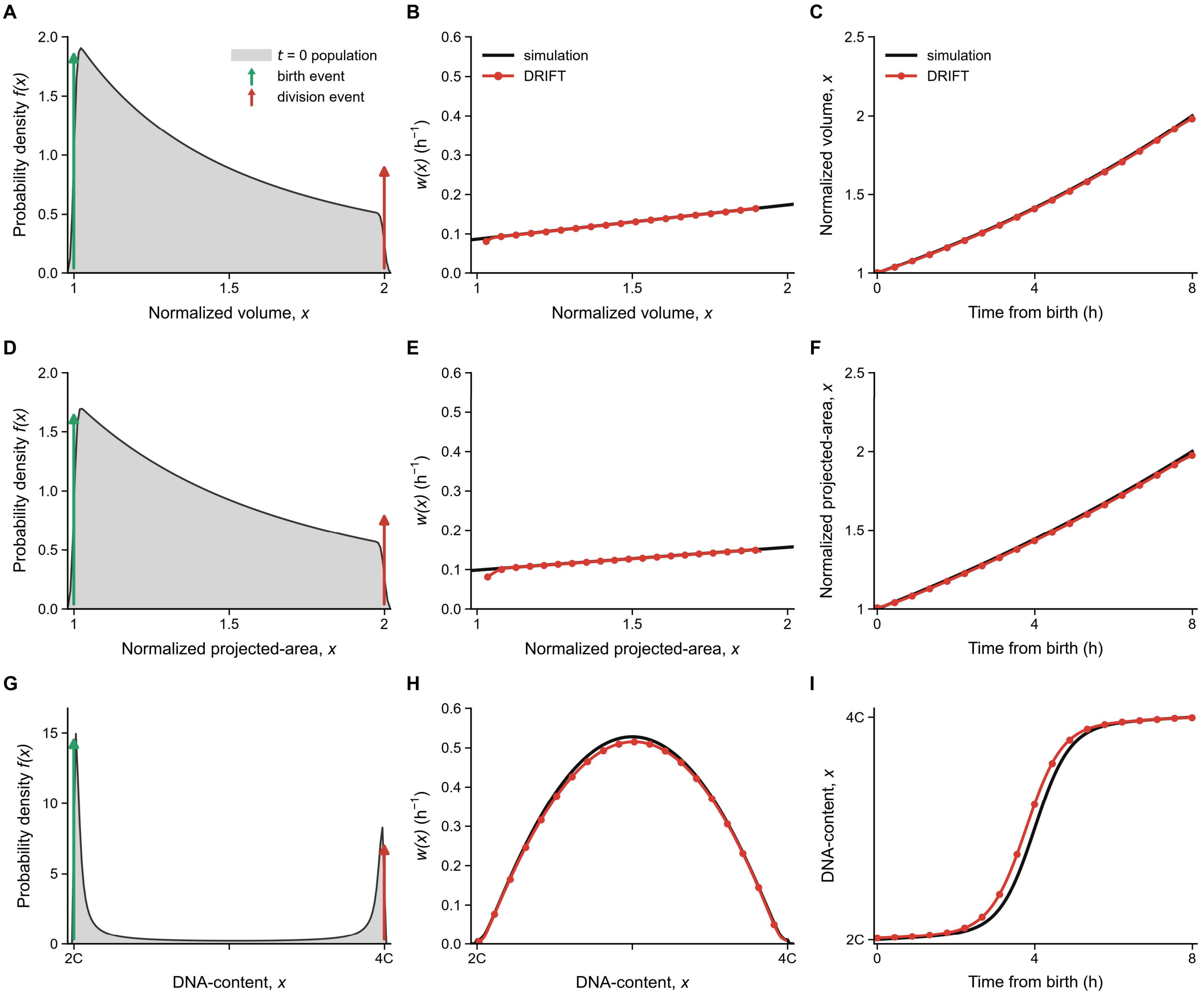


**Supplementary Fig. 2. DRIFT recovers the velocity field and the characteristic trajectory in deterministic, stationary simulations of volume, projected-area, and DNA-content.** Rows show volume (**A**–**C**), projected-area (**D**–**F**), and DNA-content (**G**–**I**). Every cell followed the same 8-h cycle. Birth and division therefore occurred at single states, and the state distribution did not change with time. (**A**, **D**, and **G**) Stationary probability densities $f(x)$; green and red arrows mark the simulated birth and division states, respectively. (**B**, **E**, and **H**) Simulated velocity fields $w(x)$ are shown in black and DRIFT estimates as red points connected by solid lines. (**C**, **F**, and **I**) Simulated birth-to-division characteristic trajectories are shown in black and DRIFT characteristic trajectories in red over the 8-h cycle. Each simulated curve starts at the deterministic birth state, whereas each DRIFT characteristic trajectory starts at the mean of the DRIFT-inferred birth-state density. Projected-area is displayed on a birth-to-division axis rescaled from 1 to 2, but its errors were calculated in the native coordinate. Velocity field $w(x)$ RRMSEs were 0.776%, 2.259%, and 2.389%, and characteristic-trajectory RRMSEs were 0.616%, 0.904%, and 2.872% for volume, projected-area, and DNA-content, respectively (Supplementary Note 3).

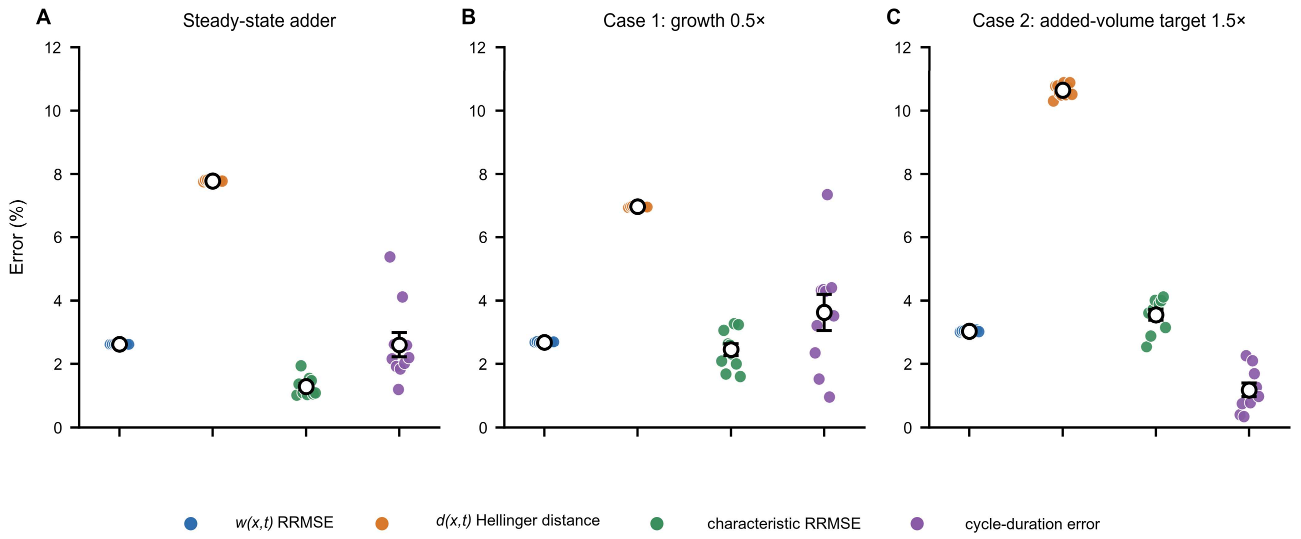


**Supplementary Fig. 3. DRIFT recovery is consistent across 10 independent simulations in each condition.** (**A**) Steady-state stochastic-adder population. (**B**) Case 1 with a 0.5-fold volume-growth rate. (**C**) Case 2 with a 1.5-fold added-volume target. Each panel shows velocity field *w(x,t)* RRMSE, division-state density *d(x,t)* Hellinger distance, characteristic-trajectory RRMSE, and cycle-duration error; colors identify the four metrics. Each colored point represents one independent simulation ($n$ = 10); open black circles and bars show mean ± SEM. Supplementary Table 3 reports the numerical values.

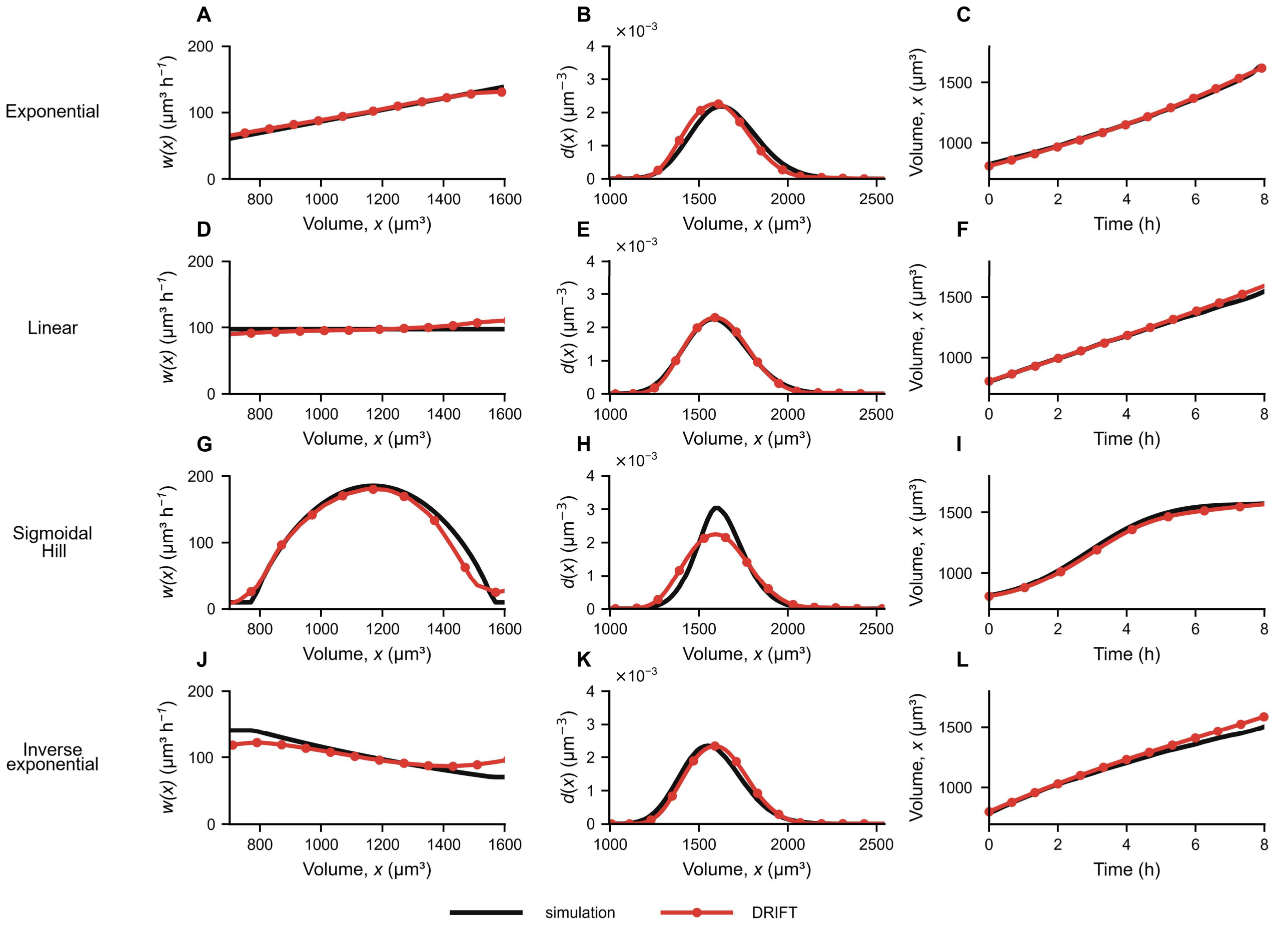


**Supplementary Fig. 4. DRIFT recovers four prescribed single-cell growth profiles in stochastic-adder simulations.** Rows show simulations with exponential (**A**–**C**), linear (**D**–**F**), sigmoidal Hill (**G**–**I**), or inverse exponential (**J**–**L**) growth profiles. Columns show the velocity field $w(x)$ (**A**, **D**, **G**, and **J**), the division-state density (**B**, **E**, **H**, and **K**), and the characteristic trajectory (**C**, **F**, **I**, and **L**). Black curves are simulated references; red points connected by solid lines are DRIFT estimates. Curves show one representative of the $n$ = 10 independent simulations per profile. Supplementary Table 4 reports the numerical values.

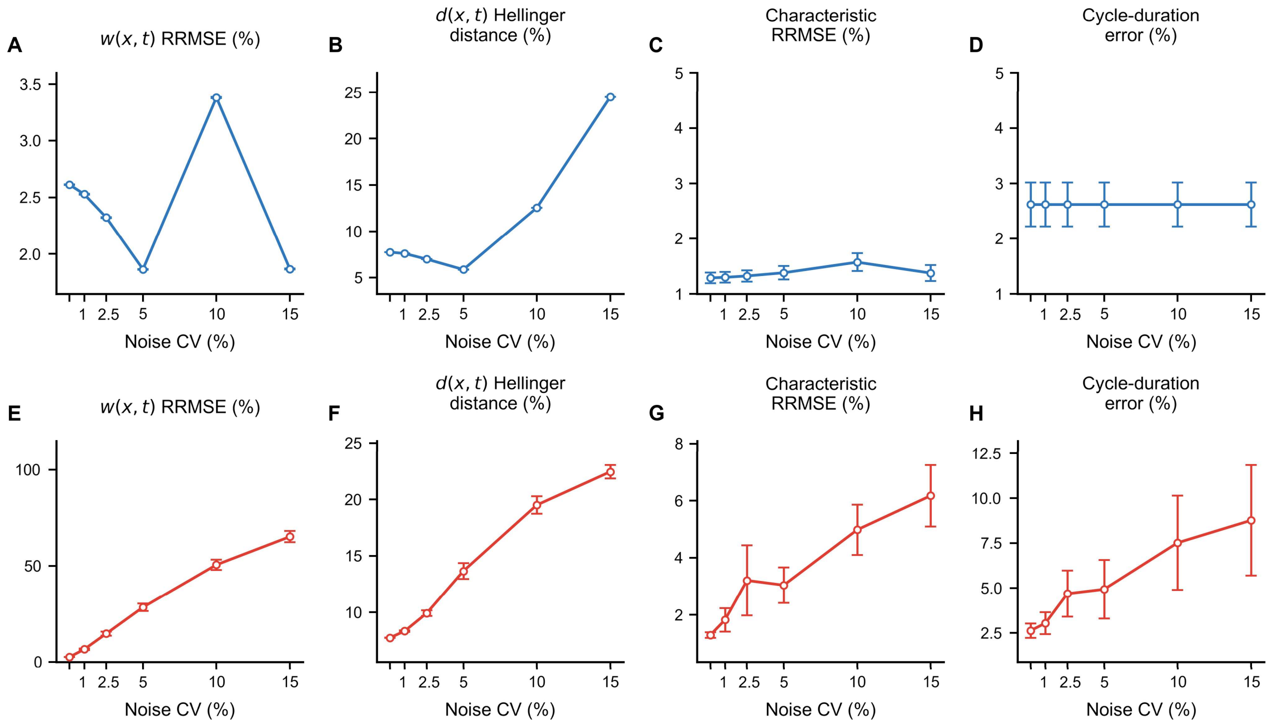


**Supplementary Fig. 5. DRIFT is more sensitive to population-count noise than to state-measurement noise.** (**A**–**D**) State-measurement noise was varied while $N(t)$ remained exact, and (**E**–**H**) population-count noise was varied while the state distributions remained exact, both in the steady-state stochastic-adder population of Fig. 2. (**A** and **E**) Velocity field *w(x,t)* RRMSE. (**B** and **F**) Division-state density *d(x,t)* Hellinger distance. (**C** and **G**) Characteristic-trajectory RRMSE. (**D** and **H**) Cycle-duration error. Open circles and bars show mean ± SEM across $n$ = 10 paired simulations at CV = 0, 1, 2.5, 5, 10, and 15%. Supplementary Table 5 reports the numerical values.

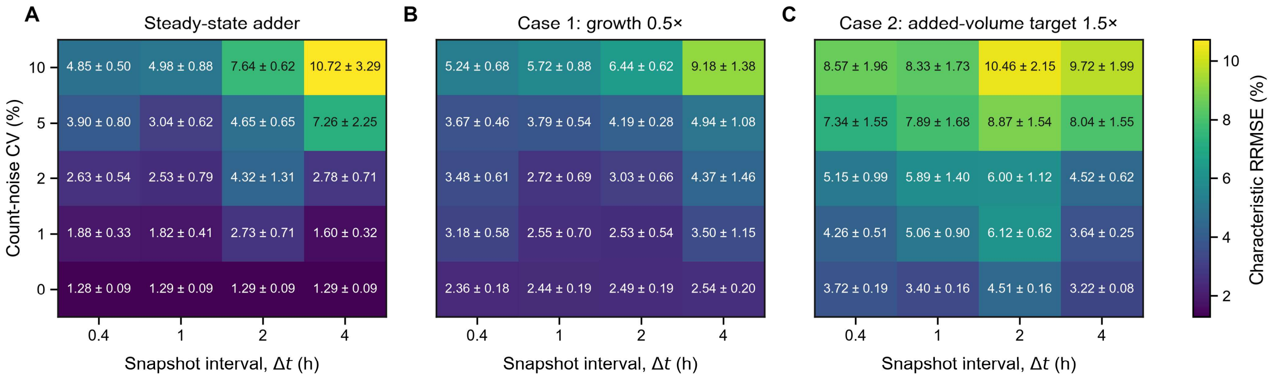


**Supplementary Fig. 6. Count-noise level and snapshot interval affect characteristic-trajectory RRMSE.** Heat maps show the characteristic-trajectory RRMSE for (**A**) the steady-state stochastic-adder population, (**B**) Case 1, in which the single-cell volume-growth rate in a stochastic-adder population was halved at $t = 0$ h while the division-size rule remained unchanged, and (**C**) Case 2, in which the single-cell volume-growth rate remained unchanged and the added-volume target was increased to 1.5 times its pre-perturbation value at $t = 0$ h. Columns show snapshot intervals of 0.4, 1, 2, and 4 h; rows show count-noise CVs of 0, 1, 2, 5, and 10%. Count noise was added only to *N(t)*. The color bar and numbers in the cells indicate mean and mean ± SEM across $n = 10$ paired simulations, respectively.

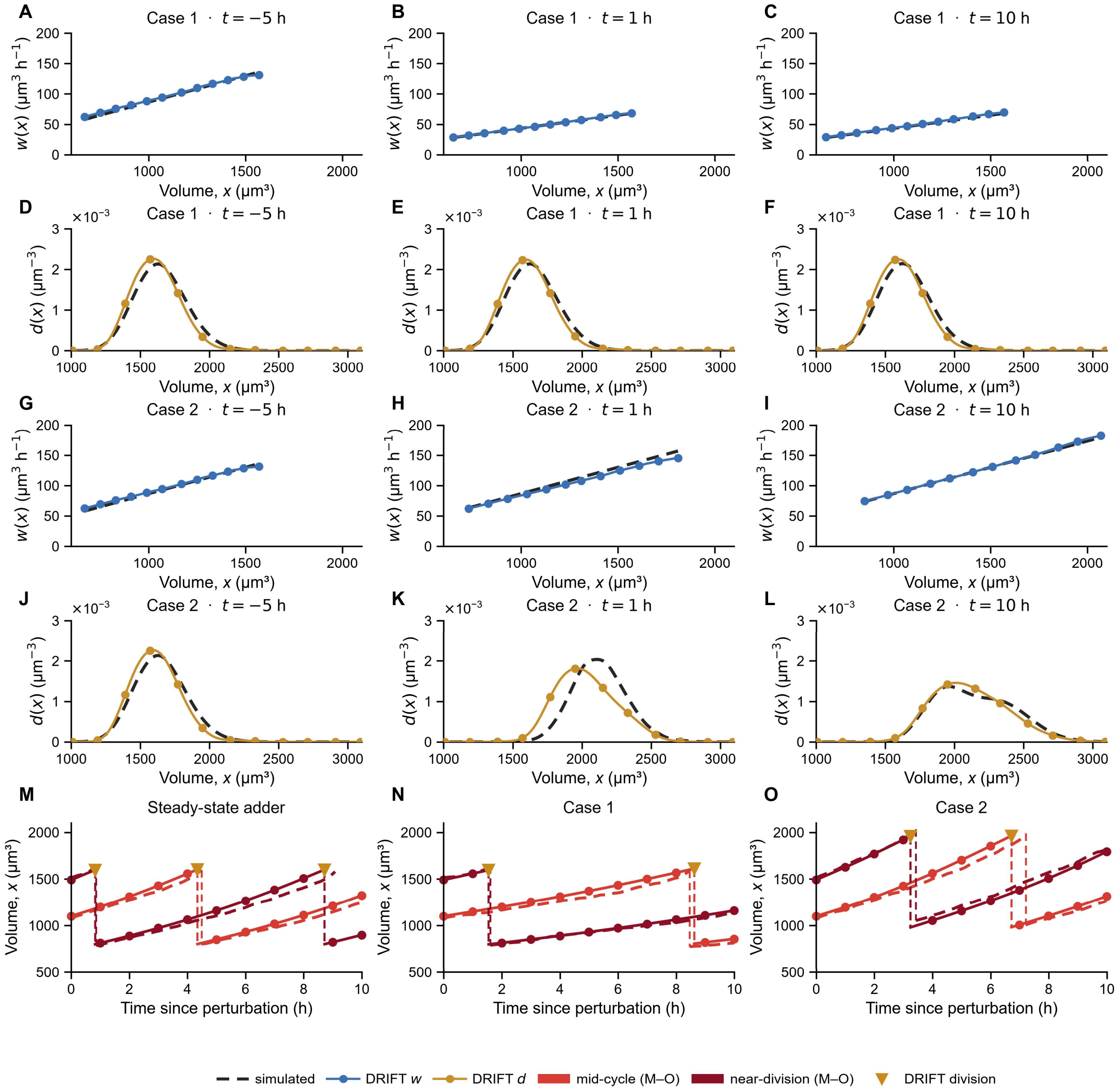


**Supplementary Fig. 7. DRIFT recovers perturbed volume dynamics across the cell cycle.** Dashed curves show simulations, and solid curves with markers show DRIFT estimates. (**A**–**C** and **G**–**I**) Velocity fields *w(x,t)* (blue) at −5, 1, and 10 h relative to the perturbation for Case 1, in which the single-cell volume-growth rate in a stochastic-adder population was halved at $t$ = 0 h while the division-size rule remained unchanged (**A**–**C**) and Case 2, in which the single-cell volume-growth rate remained unchanged and the added-volume target was increased to 1.5 times its pre-perturbation value at $t$ = 0 h (**G**–**I**). (**D**–**F** and **J**–**L**) Division-state densities *d(x,t)* (ochre) at the same time points for Case 1 (**D**–**F**) and Case 2 (**J**–**L**). (**M**–**O**) Characteristic trajectories initialized from the mid-cycle (red) and near-division (dark red) states (pretreatment phases 0.45 and 0.90; see Supplementary Note 2 for additional details) for the steady-state stochastic-adder population (**M**), Case 1 (**N**), and Case 2 (**O**); ochre triangles mark DRIFT division events. All panels show one representative of the $n$ = 10 independent simulations.

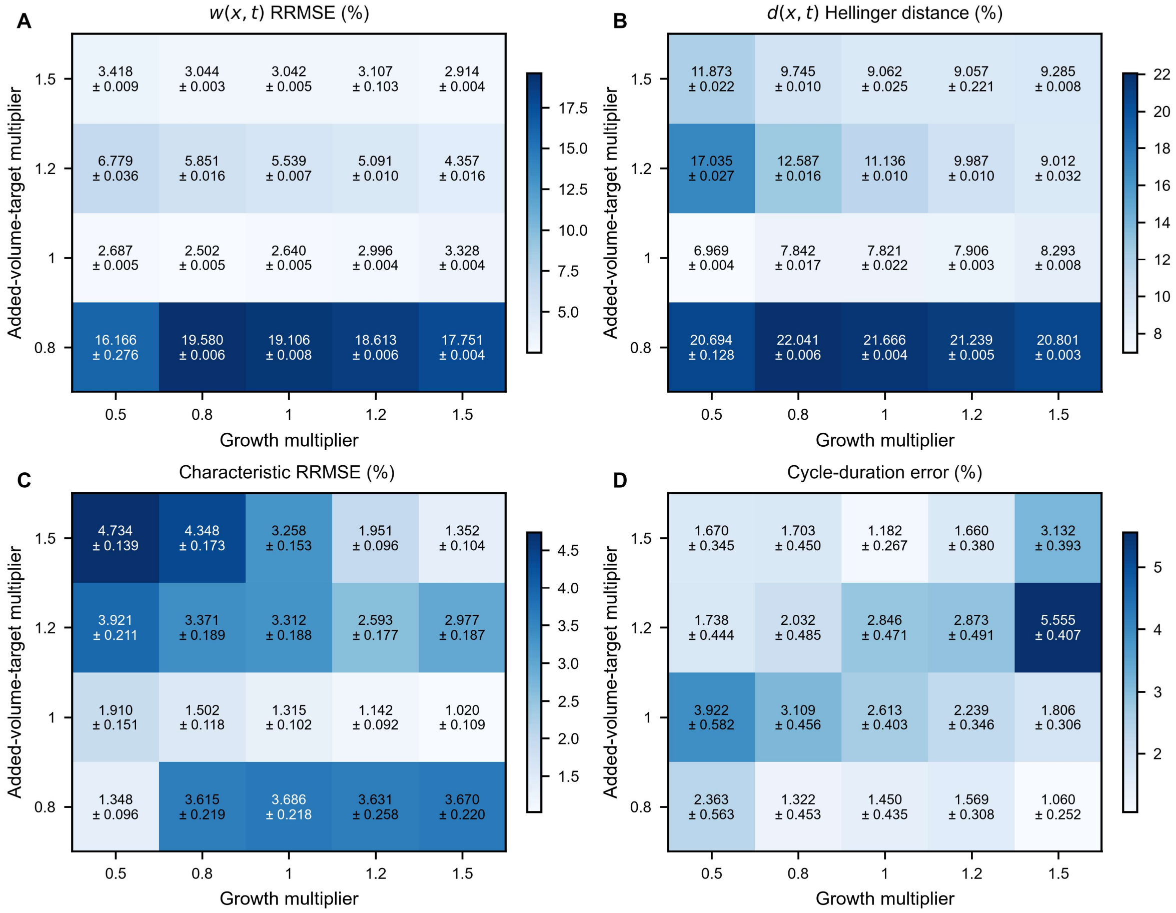


**Supplementary Fig. 8. DRIFT recovers volume dynamics across combined changes in the volume-growth rate and the added-volume target.** Heat maps show recovery metrics for combined perturbations applied to the stochastic-adder population at $t = 0$ h. (**A**) Velocity field *w(x,t)* RRMSE. (**B**) Division-state density *d(x,t)* Hellinger distance. (**C**) Characteristic-trajectory RRMSE. (**D**) Cycle-duration error. In **A**–**D**, columns are volume-growth-rate multipliers of 0.5, 0.8, 1, 1.2, and 1.5 and rows are added-volume-target multipliers of 0.8, 1, 1.2, and 1.5. The color bar and numbers in the cells indicate mean and mean ± SEM across 10 independent simulations, respectively.

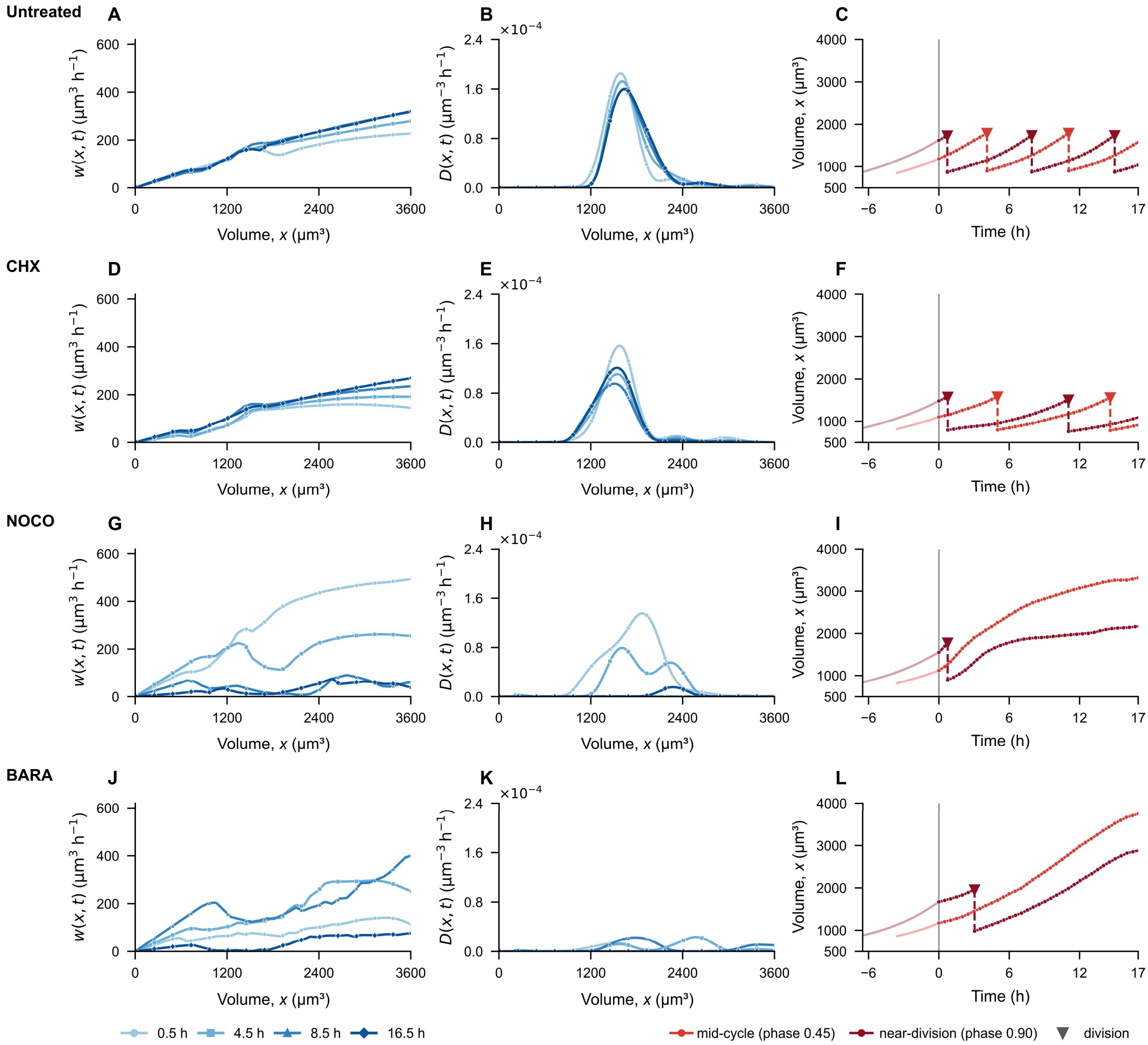


**Supplementary Fig. 9. Whether an inferred division occurs within the measurement window depends on the initial state under nocodazole and barasertib.** Rows show untreated (**A**–**C**), cycloheximide-treated (CHX; **D**–**F**), nocodazole-treated (NOCO; **G**–**I**), and barasertib-treated (BARA; **J**–**L**) L1210 cells. Columns show velocity fields *w(x,t)* (**A**, **D**, **G**, and **J**), mother-division-loss densities *D(x,t)* (**B**, **E**, **H**, and **K**; Supplementary Note 2), and characteristic trajectories (**C**, **F**, **I**, and **L**). The velocity fields and mother-division-loss densities are shown at 0.5, 4.5, 8.5, and 16.5 h, and color progresses from light to dark blue. The trajectories were initialized at the mid-cycle or the near-division pretreatment state (red and dark red; pretreatment phases 0.45 and 0.90, respectively; Supplementary Note 2). Pale segments show pretreatment progression, inverted triangles mark completed divisions, and dashed vertical segments connect the mother and daughter states. In **C** and **F**, both initializations complete a division within 17 h, whereas in **I** and **L** only the near-division initialization does. Each condition was fitted once from the replicate-averaged volume probability density ($n$ = 1, 1, 2, and 3 independent experiments for untreated, CHX, NOCO, and BARA, respectively; Supplementary Note 4).

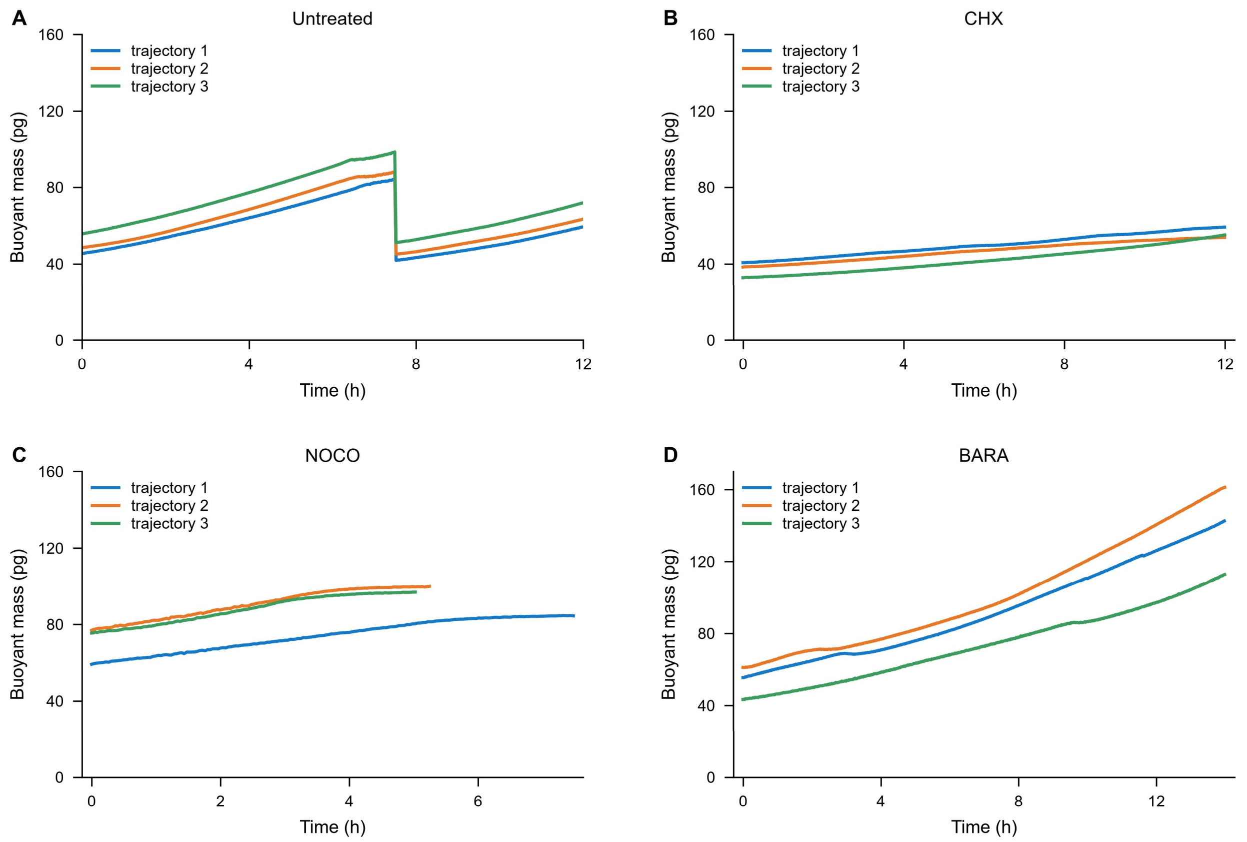


**Supplementary Fig. 10. Single-cell buoyant-mass traces show cell-to-cell variation within each condition.** (**A**–**D**) Untreated, cycloheximide-treated (CHX), nocodazole-treated (NOCO), and barasertib-treated (BARA) L1210 cells, respectively. Each panel shows three separately measured cells, and each colored line is one longitudinal buoyant-mass trace recorded with a suspended microchannel resonator. The CHX and NOCO traces are from *ref. 2*, and the untreated and BARA traces from *ref. 3*.

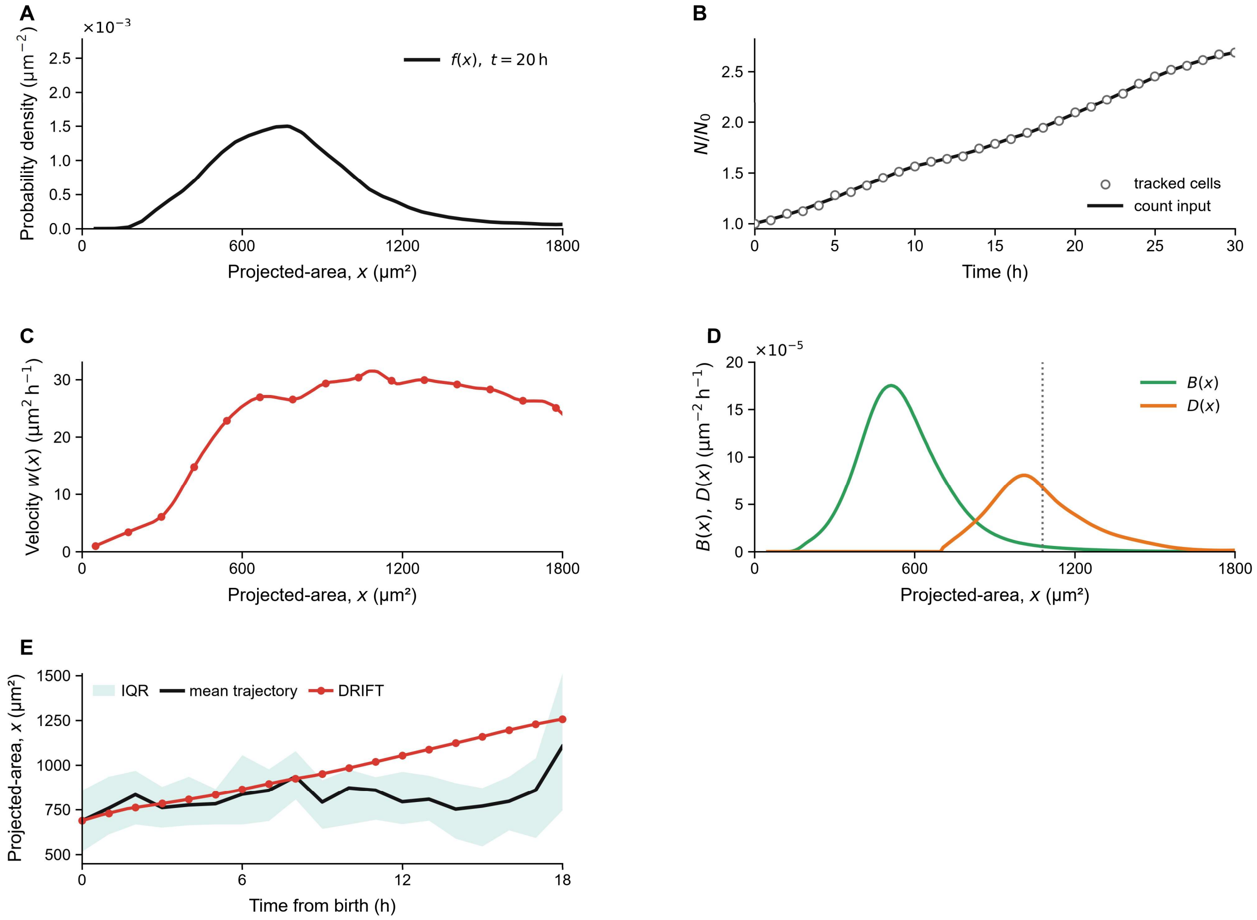


**Supplementary Fig. 11. The stationary projected-area characteristic trajectory follows the independently tracked newborn cohort except during mitotic rounding.** (**A**) The projected-area probability density measured at 20 h in a single untreated well, held fixed as the stationary input *f(x)* (black). (**B**) Number of objects that passed quality control at each hourly time point in the same well (open circles) and the fitted count curve supplied to the inverse (black), both normalized to the first snapshot and plotted on the internal 0–30-h axis. (**C**) Inferred velocity field *w(x)*, taken as the median across the fitted intervals. (**D**) Physical daughter-birth density *B(x)* (green) and mother-division-loss density *D(x)* (orange), shown as pointwise medians over the same fitted intervals; the dotted line marks the median division state (Section Table 5). (**E**) Mean (black) and interquartile range (pale blue) of 29 newborn-aligned tracked cycles together with the DRIFT-inferred characteristic trajectory (red).

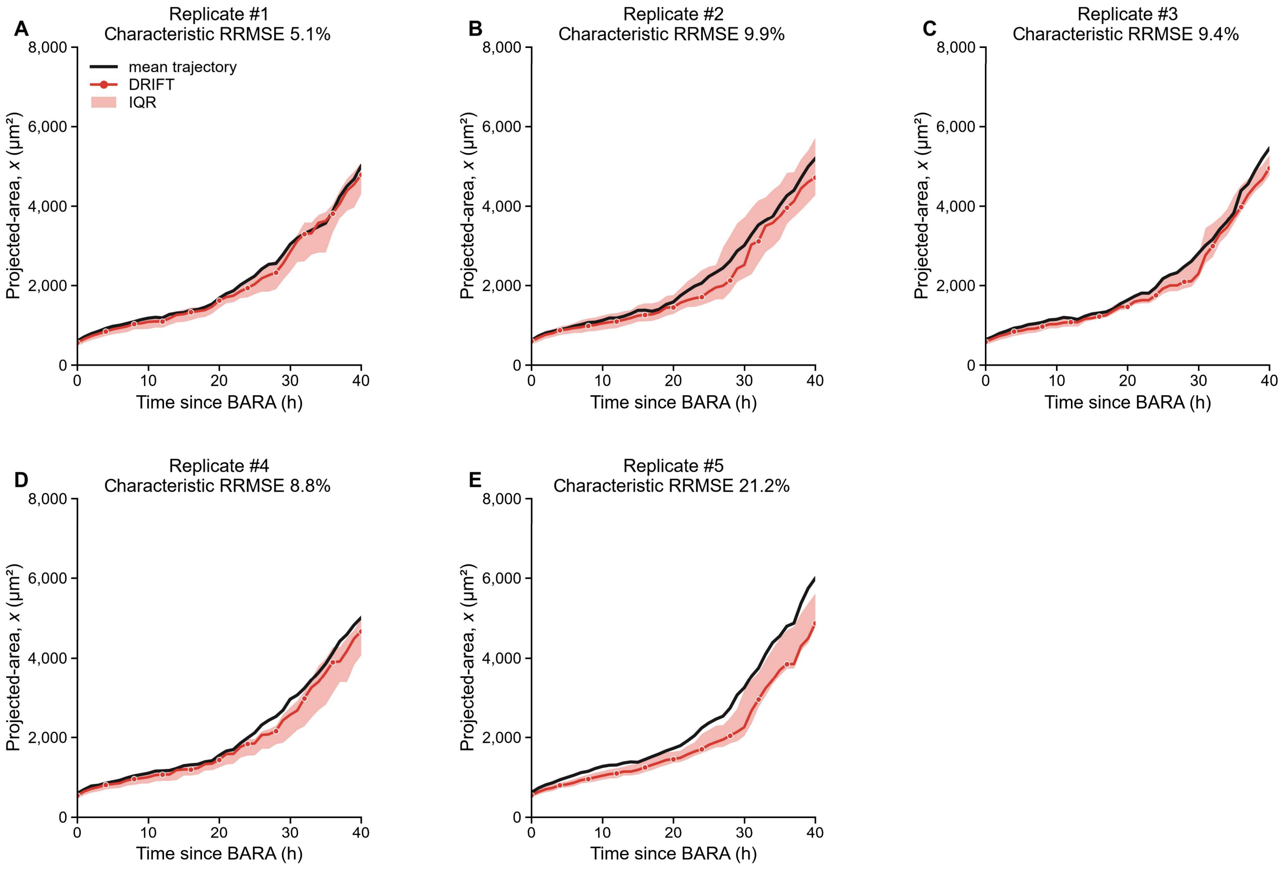


**Supplementary Fig. 12. DRIFT characteristic trajectories follow the tracked mean projected-area increase.** (**A**–**E**) Mean tracked trajectories (black) and DRIFT characteristic trajectories (red) for biological replicates #1–#5, respectively, each comprising 100 retained tracks from 0 to 40 h and fitted with its own velocity field. The birth-state density of each replicate was inferred from its own $t$ = 0 snapshot; its mean initialized the central DRIFT characteristic trajectory, and its 25th and 75th percentiles initialized the pink band. The band shows dependence on the initial state. Supplementary Table 7 reports the values.

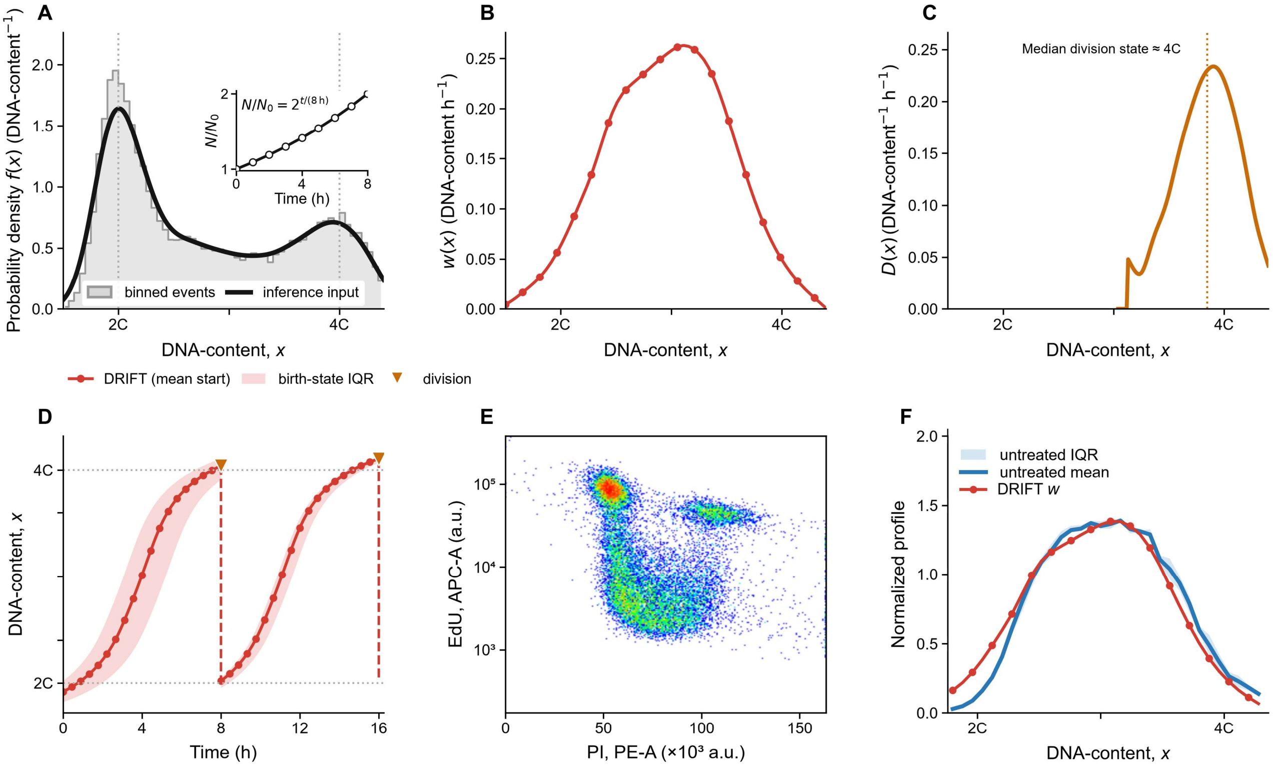


**Supplementary Fig. 13. The stationary DNA velocity field reproduces a closed 2C-to-4C cycle and matches the independent EdU-incorporation profile.** (**A**) Pooled DNA-content distribution from three separately processed pretreatment PI/RNase acquisitions. The gray histogram shows the equal-weight distribution of gated events, the black curve is the stationary input supplied to the inverse, and the inset shows the assumed count increase $N/N_0$ for an 8-h doubling time. (**B**) Pooled stationary DRIFT-inferred velocity field *w(x)*. (**C**) Pooled mother-division-loss density *D(x)*; the dotted line marks the median division state. (**D**) Two-cycle characteristic trajectories obtained by integrating the same pooled stationary *w(x)* in panel **B** from the mean (red) and from the 25th and 75th percentiles (pink band) of the birth-state density *b(x)*. Ochre triangles mark division events, and dashed drops connect division and daughter states. (**E**) PI–EdU event map from a single acquisition after a 30-min EdU pulse. (**F**) Equal-weight mean and interquartile range of background-corrected EdU profiles from four untreated acquisition files compared with pooled stationary DRIFT *w(x)* from $n = 3$ biological replicates.

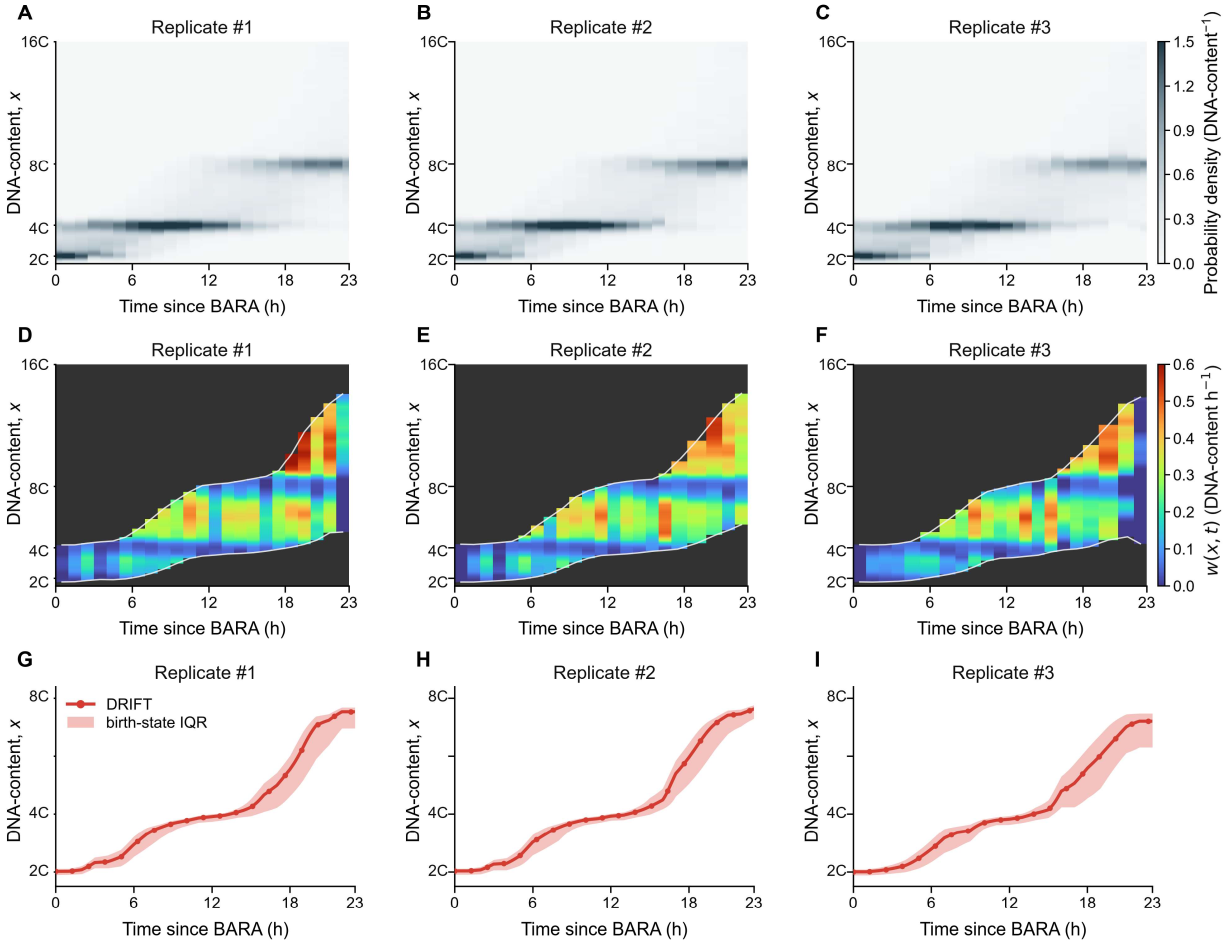


**Supplementary Fig. 14. The staircase-like DNA-content increase after barasertib treatment.** Columns correspond to biological replicates #1–#3 of barasertib-treated L1210 cells measured by PI flow cytometry. (**A**–**C**) Time-resolved DNA-content distributions; grayscale encodes probability density. (**D**–**F**) Velocity fields $w(x,t)$; color encodes velocity magnitude and is shown only between the 5th and 95th cumulative-mass states. Dark-gray regions lie outside this displayed support, and white curves mark its limits. (**G**–**I**) Characteristic trajectories initialized at the mean of each replicate's $t = 0$ birth-state density (red), with bands initialized at its 25th and 75th percentiles (pink). Supplementary Table 7 reports the selected penalty for these records.

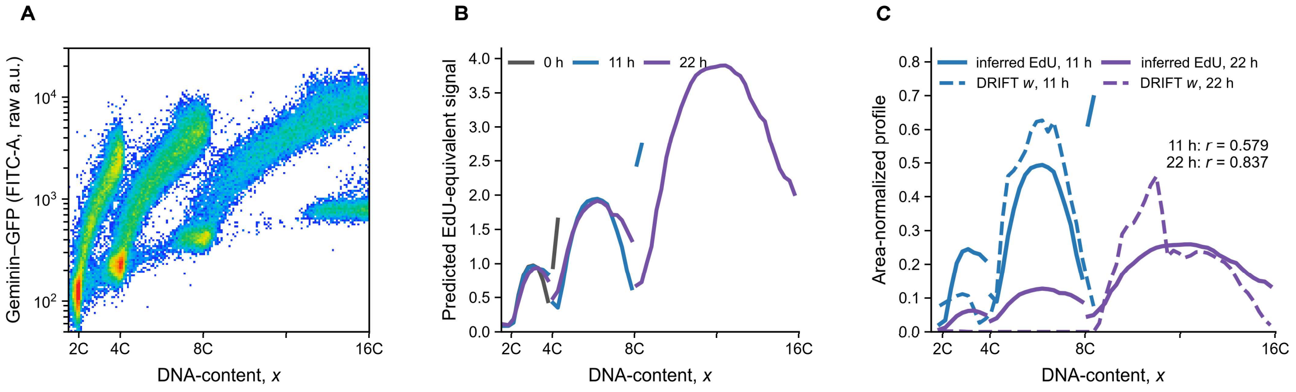


**Supplementary Fig. 15. Geminin-based EdU-equivalent profiles and the DRIFT DNA velocity field are positively correlated at both treatment times.** (**A**) Pooled PI–Geminin-GFP event-density map from four acquisitions of one barasertib-treated experiment with L1210 cells expressing the fluorescent ubiquitination-based cell cycle indicator (FUCCI). Each acquisition contained a mixture of samples collected at 0, 11, and 22 h after treatment. PI fluorescence reports DNA-content and Geminin-GFP fluorescence reports cell-cycle position. (**B**) Geminin-based EdU-equivalent profiles from six separately labeled files of the same experiment, two at each of the treatment times 0, 11, and 22 h. (**C**) Time-matched EdU-equivalent and DRIFT velocity profiles *w(x,t)* at 11 and 22 h. Gray, blue, and purple denote 0, 11, and 22 h, respectively, and in **C** solid and dashed lines denote the EdU-equivalent and the DRIFT velocity profiles, respectively. Each profile was normalized over the measured DNA range. The DRIFT velocity profiles in **C** are the pointwise medians of the three replicate-specific profiles. The *r* values in **C** are the Pearson correlations between the displayed profiles.

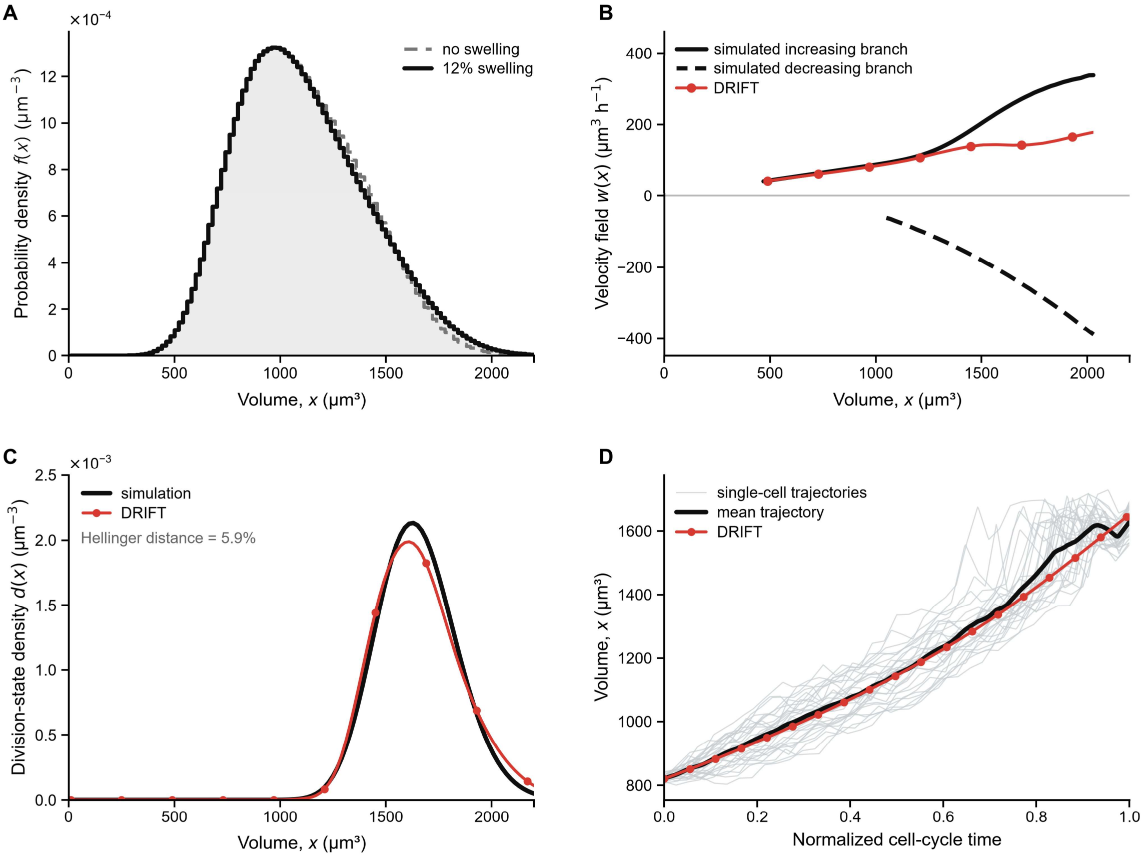


**Supplementary Fig. 16. Effect of the nonnegative-velocity constraint on reconstruction of mitotic swelling dynamics.** (**A**) Stationary volume distributions from simulations with no swelling (gray dashed) and 12% swelling (black), in which the measured volume rose by up to 12% late in the cycle and returned to its unswollen value before division. (**B**) Simulated increasing (solid black) and decreasing (dashed black) velocity branches together with the single DRIFT velocity field *w(x)* (red). (**C**) Simulated division-state density *d*(*x*) (black) and DRIFT projected division-state density (red). (**D**) Individual simulated cell-volume trajectories (light gray; 30 lineages shown), the mean trajectory of the same 30 lineages (black), and the DRIFT-inferred characteristic trajectory (red) over one normalized cell cycle. The characteristic-trajectory RRMSE was 1.96%, and the first-cycle duration error was 0.38% (0.03 h).

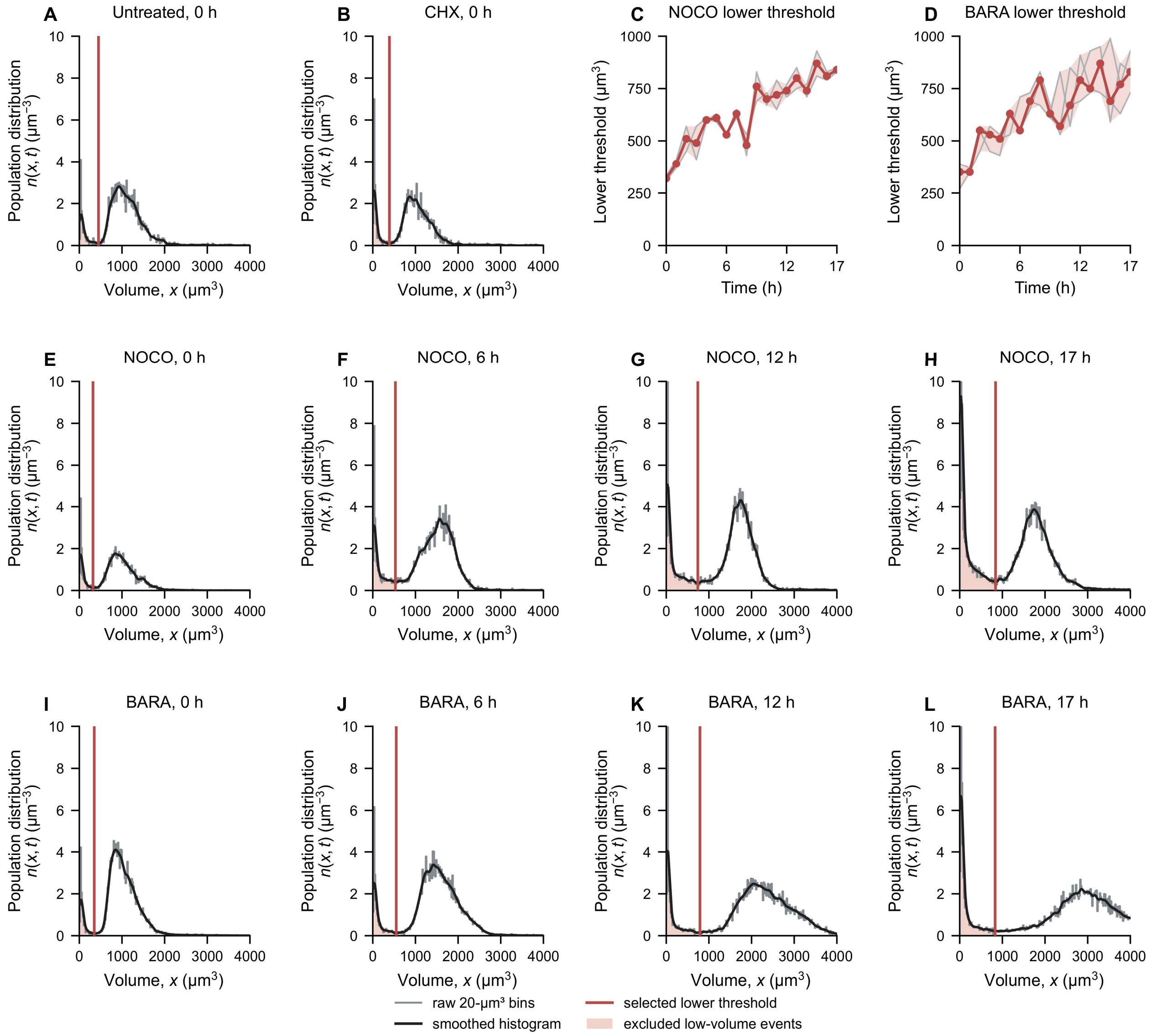


**Supplementary Fig. 17. A well-resolved valley between the low-volume and cell peaks defines the lower volume threshold.** Histograms in **A**, **B**, and **E**–**L** were averaged over the $n$ = 1, 1, 2, and 3 independent experiments for untreated, CHX, NOCO, and BARA, respectively, and each experiment was thresholded separately. (**A** and **B**) Untreated and cycloheximide-treated (CHX) 0-h volume population distributions $n(x,t)$, respectively. Gray lines show the raw 20-µm³ bins, black curves show the smoothed histogram used to locate the two peaks and the intervening valley, red lines mark the selected lower thresholds (median across replicates), and pale red shading marks the excluded low-volume events attributed to debris and dead cells. (**C** and **D**) Lower thresholds for the nocodazole (NOCO) and barasertib (BARA) records from 0 to 17 h, respectively. Gray lines show individual replicates, red points and lines show the median, and shading shows the replicate range. (**E**–**H**) Nocodazole histograms and (**I**–**L**) barasertib histograms at 0, 6, 12, and 17 h with the selected thresholds. The low volume threshold detection method is adapted from *ref. 1*. See Supplementary Note 4 and Section Table 2 for more details.

### Supplementary Tables

### Supplementary Table 1. Notation and fixed settings for DRIFT inference and characteristic-trajectory construction.

| notation or setting | definition | value |
|---|---|---|
| $x$, $t$ | measured state coordinate and time | state unit; h |
| $N(t), r_q, N_{\mathrm{fit}}(t), r_{\mathrm{fit}}(t)$ | cell count, interval log-count slope, monotone count fit, and its instantaneous rate | count; $h^{-1}$ |
| $f(x,t)$, $F(x,t)$ | unit-mass state probability density and its cumulative distribution | f: $\mathrm{state}^{-1}$; F: dimensionless |
| $w(x,t)$ | inferred state velocity | state unit $h^{-1}$ |
| $g = \frac{T_0 w}{x}$ | dimensionless specific rate | $T_0 = 1$ h |
| $d$, $\tilde{d}$ | unit-mass division-state density and its projected output | $\mathrm{state}^{-1}$ |
| $b(x,t)$ | birth-state density; integrates to two daughters per division | $\mathrm{state}^{-1}$ |
| $D = rd$ | physical division-loss density, plotted from the projected density $\tilde{d}$ | $\mathrm{state}^{-1}$ $h^{-1}$ |
| $B(x,t)$ | daughter-birth density, $B = 2KD$ | $\mathrm{state}^{-1}$ $h^{-1}$ |
| $K$ | single-daughter partition operator | column mass 1 |
| $M = 2K - I$ | net binary-fission source operator | dimensionless |
| $T_{\mathrm{inst}}(t)$ | instantaneous count-derived period | $\frac{\ln(2)}{r_{\mathrm{fit}}(t)}$; h |
| $\varphi_0$ | initial pretreatment cycle fraction | 0; 0.45; 0.90 for newborn-like, mid-cycle, and near-division starts |
| elapsed-time criterion | elapsed-time threshold for first and later events | $(1-\varphi_0)T_{\mathrm{inst}}(t)$; $T_{\mathrm{inst}}(t)$ |
| $x_{D,0.5}(t)$ | median of the projected division-state density | state unit |
| $K_w$, $K_d$ | velocity and division-density basis dimensions | 64; 12 |
| $K_{proj}$ | projection dimension; smoothing penalty | 24; 0.01 |
| $\lambda_g$ | specific-rate penalty (weak; moderate) | $3 \times 10^{-4}$; $3 \times 10^{-3}$ |
| other penalties | spatial and temporal difference penalties | $\lambda_{w,x} = 10^{-6}$; $\lambda_{d,x} = 10^{-12}$; $\lambda_{w,t} = 10$; $\lambda_{d,t} = 10^{-2}$ |
| penalty selection | moderate retained if both conditions hold | $\frac{R_{\mathrm{mod}}}{R_{\mathrm{weak}}} \leq 1.25$; $R_{\mathrm{mod}} - R_{\mathrm{weak}} \leq 0.25\, pp$ |
| partition kernel | daughter-fraction distribution (volume analyses) | symmetric Beta(15,15) |
| partition kernel | daughter-state map (projected-area analyses) | deterministic projected-area division map; empirical daughter–parent kernel for the stationary control |
| partition kernel | daughter-state map (DNA-content analyses) | deterministic one-half map |
| Case 1: volume-growth rate 0.5× | post-treatment growth coefficient | k: 0.08664 → 0.04332 $h^{-1}$ |
| Case 2: added-volume target 1.5× | post-treatment added-volume target | Δ: 780 → 1,170 μm³ |
| synthetic grid | state grid and standard snapshot interval | 0–4,500 μm³; 20-μm³ bins; 1 h (0.4, 1, 2, and 4 h exact subsets in Supplementary Fig. 6) |
| initial population | seed cells per simulation | 2,000 |
| equilibration | steady-state criterion | symmetric KL divergence < $2 \times 10^{-4}$ on four consecutive 2-h checks; 78–84 h reached |
| Coulter grid | state grid and division support | 10–5,990 μm³; 210–3,990 μm³ |
| integration | characteristic-trajectory solver | Heun (RK2); 0.005-h step |
| scoring support | velocity; division-state density | central 90% of f mass; declared division support |

**Supplementary Table 2. Sensitivity of recovery metrics to the specific-rate penalty and velocity/division basis sizes (median across valid simulated records).**

| estimator component | tested setting | valid fits | velocity-field RRMSE (%) | projected Hellinger distance (%) | characteristic-trajectory RRMSE (%) | cycle-duration error (%) |
|---|---|---|---|---|---|---|
| Specific-rate penalty | $\lambda g = 0$ | 5/6 valid | 10.377 | 12.538 | 1.296 | 5.202 |
| | $\lambda g = 1 \times 10^{-4}$ | 6/6 valid | 4.976 | 8.175 | 1.916 | 3.325 |
| | $\lambda g = 3 \times 10^{-4}$ | 6/6 valid | 4.618 | 7.906 | 1.997 | 3.325 |
| | $\lambda g = 1 \times 10^{-3}$ | 6/6 valid | 4.696 | 7.703 | 1.978 | 3.325 |
| | $\lambda g = 3 \times 10^{-3}$ | 6/6 valid | 4.974 | 7.851 | 1.979 | 3.325 |
| | $\lambda g = 1 \times 10^{-2}$ | 6/6 valid | 5.547 | 7.596 | 2.061 | 3.325 |
| Basis size | Velocity: 48; division-state density: 8 | 6/6 valid | 17.270 | 36.200 | 2.991 | 13.710 |
| | Velocity: 48; division-state density: 12 | 6/6 valid | 4.275 | 7.698 | 1.875 | 3.325 |
| | Velocity: 48; division-state density: 16 | 6/6 valid | 9.197 | 10.884 | 2.810 | 3.376 |
| | Velocity: 64; division-state density: 8 | 6/6 valid | 17.301 | 36.252 | 2.968 | 13.710 |
| | Velocity: 64; division-state density: 12 | 6/6 valid | 4.289 | 7.690 | 1.884 | 3.325 |
| | Velocity: 64; division-state density: 16 | 6/6 valid | 8.725 | 10.466 | 2.673 | 2.940 |
| | Velocity: 80; division-state density: 8 | 6/6 valid | 17.299 | 36.273 | 2.968 | 13.740 |
| | Velocity: 80; division-state density: 12 | 6/6 valid | 4.303 | 7.680 | 1.883 | 3.325 |
| | Velocity: 80; division-state density: 16 | 5/6 valid | 14.174 | 11.469 | 3.126 | 2.620 |

**Supplementary Table 3. Recovery metrics for the simulated populations in Figs. 2 and 3 (mean ± SEM, *n* = 10 independent simulations).**

| simulated population | velocity-field RRMSE (%) | kernel-mapped birth-state Hellinger distance (%) | projected Hellinger distance (%) | characteristic-trajectory RRMSE (%) | cycle-duration error (%) |
|---|---|---|---|---|---|
| Steady-state stochastic-adder population | 2.621 ± 0.003 | 3.608 ± 0.007 | 7.787 ± 0.008 | 1.283 ± 0.095 | 2.608 ± 0.391 |
| Case 1: volume-growth rate 0.5× | 2.687 ± 0.005 | — | 6.969 ± 0.004 | 2.180 ± 0.163 | 3.633 ± 0.571 |
| Case 2: added-volume target 1.5× | 3.042 ± 0.005 | — | 10.640 ± 0.060 | 3.502 ± 0.152 | 1.109 ± 0.191 |

**Supplementary Table 4. Recovery metrics across four prescribed single-cell growth profiles (mean ± SEM, *n* = 10 independent simulations per growth profile).**

| single-cell growth profile | velocity-field RRMSE (%) | projected Hellinger distance (%) | characteristic-trajectory RRMSE (%) | cycle-duration error (%) |
|---|---|---|---|---|
| Exponential | 2.621 ± 0.003 | 7.787 ± 0.008 | 1.283 ± 0.095 | 2.608 ± 0.391 |
| Linear | 5.536 ± 0.004 | 4.553 ± 0.010 | 1.523 ± 0.269 | 2.234 ± 0.454 |
| Sigmoidal Hill | 10.446 ± 0.002 | 12.364 ± 0.013 | 3.647 ± 0.417 | 8.614 ± 1.370 |
| Inverse exponential | 10.716 ± 0.003 | 7.619 ± 0.006 | 1.261 ± 0.071 | 2.835 ± 0.520 |

**Supplementary Table 5. Recovery metrics under independently varied state-measurement noise or population-count noise at 1-h snapshot intervals (mean ± SEM, *n* = 10 paired simulations per setting).**

| error source | CV (%) | velocity-field RRMSE (%) | projected Hellinger distance (%) | characteristic-trajectory RRMSE (%) | cycle-duration error (%) |
|---|---|---|---|---|---|
| Population-count noise | 0 | 2.612 ± 0.003 | 7.734 ± 0.003 | 1.288 ± 0.094 | 2.619 ± 0.398 |
| | 1 | 6.602 ± 0.388 | 8.319 ± 0.063 | 1.819 ± 0.413 | 3.034 ± 0.609 |
| | 2.5 | 14.744 ± 1.021 | 9.901 ± 0.257 | 3.205 ± 1.232 | 4.678 ± 1.290 |
| | 5 | 28.435 ± 1.982 | 13.650 ± 0.700 | 3.038 ± 0.620 | 4.923 ± 1.631 |
| | 10 | 50.587 ± 2.683 | 19.496 ± 0.771 | 4.977 ± 0.882 | 7.512 ± 2.621 |
| | 15 | 65.201 ± 2.941 | 22.440 ± 0.595 | 6.168 ± 1.074 | 8.758 ± 3.077 |
| State-measurement noise | 0 | 2.612 ± 0.003 | 7.734 ± 0.003 | 1.288 ± 0.094 | 2.619 ± 0.398 |
| | 1 | 2.528 ± 0.005 | 7.597 ± 0.003 | 1.298 ± 0.097 | 2.619 ± 0.398 |
| | 2.5 | 2.322 ± 0.004 | 6.988 ± 0.003 | 1.323 ± 0.101 | 2.619 ± 0.398 |
| | 5 | 1.864 ± 0.003 | 5.868 ± 0.005 | 1.380 ± 0.120 | 2.619 ± 0.398 |
| | 10 | 3.381 ± 0.005 | 12.540 ± 0.007 | 1.571 ± 0.160 | 2.619 ± 0.398 |
| | 15 | 1.867 ± 0.004 | 24.485 ± 0.006 | 1.375 ± 0.143 | 2.619 ± 0.398 |

**Supplementary Table 6. Projected-area track summaries by biological replicate.**

| biological replicate | n | projected-area at 0 h (μm²) | projected-area at 40 h (μm²) | 40-h fold change |
|---|---|---|---|---|
| #1 | 100 | 603 [497–724] | 4,832 [3,987–5,687] | 7.90 [6.89–9.76] |
| #2 | 100 | 622 [487–763] | 4,878 [4,042–5,978] | 7.96 [6.75–9.89] |
| #3 | 100 | 627 [516–754] | 5,032 [4,088–6,200] | 7.84 [6.58–9.73] |
| #4 | 100 | 573 [480–696] | 4,727 [3,485–5,798] | 7.77 [6.41–9.71] |
| #5 | 100 | 589 [501–727] | 5,424 [4,277–7,007] | 8.83 [7.52–11.97] |

**Supplementary Table 7. Projected-area and DNA-content inference metrics.**

| state coordinate | selected specific-rate penalty | weak / moderate population-balance residual (%) | characteristic-trajectory RRMSE (%) |
|---|---|---|---|
| Projected-area, replicate #1 | weak ($3 \times 10^{-4}$) | 25.93 / 26.47 | 5.12 |
| Projected-area, replicate #2 | weak ($3 \times 10^{-4}$) | 32.23 / 32.78 | 9.91 |
| Projected-area, replicate #3 | weak ($3 \times 10^{-4}$) | 31.22 / 31.91 | 9.38 |
| Projected-area, replicate #4 | weak ($3 \times 10^{-4}$) | 28.63 / 29.12 | 8.77 |
| Projected-area, replicate #5 | weak ($3 \times 10^{-4}$) | 29.98 / 30.40 | 21.18 |
| Projected-area, mean ± SEM | weak in 5 of 5 replicates | 29.60 ± 1.10 / 30.13 ± 1.11 | 10.87 ± 2.71 |
| DNA-content | moderate ($3 \times 10^{-3}$) in all three biological replicates | — | — |
